# Genes for ash tree resistance to an insect pest identified via comparative genomics

**DOI:** 10.1101/772913

**Authors:** Laura J. Kelly, William J. Plumb, David W. Carey, Mary E. Mason, Endymion D. Cooper, William Crowther, Alan T. Whittemore, Stephen J. Rossiter, Jennifer L. Koch, Richard J. A. Buggs

## Abstract

Genome-wide discovery of candidate genes for functional traits within a species typically involves the sequencing of large samples of phenotyped individuals^1^, or linkage analysis through multiple generations^2^. When a trait occurs repeatedly among phylogenetically independent lineages within a genus, a more efficient approach may be to identify genes via detection of amino acid residues shared by species possessing that trait^3,4^. Here, by taking this approach, we identify candidate loci in the genus *Fraxinus* (ash trees) for resistance to the emerald ash borer beetle (EAB; *Agrilus planipennis*), a pest species that appears innocuous to otherwise healthy ash in its native East Asian range^5^ but is highly destructive in North America^6^ and poses a threat to ash trees in Europe^7^. Assembling whole genome sequences for 24 diploid species and subspecies of ash, and estimating resistance to EAB for 26 taxa from egg bioassays, we find 53 genes containing amino acid variants shared between two or more independent *Fraxinus* lineages with EAB-resistant species, that are unlikely to be due to chance or undetected paralogy. Of these, seven genes have putative roles relating to the phenylpropanoid biosynthesis pathway and 17 are potentially connected to herbivore recognition, defence signalling or programmed cell death. We also find that possible loss-of-function mutations among our 53 candidate genes are more frequent in susceptible species, than in resistant ones. Patterns of polymorphism for the EAB-associated amino acid variants in ash trees representing different European populations suggest that selection may be able to enhance their resistance to EAB.

---

EAB has proved more costly than any other invasive forest insect within the USA to date^8^. We assessed resistance to EAB for 26 *Fraxinus* taxa (Supplementary Table 1). Tree resistance was scored according to the instar, health and weight of EAB larvae in the stems of artificially infested trees eight weeks after infestation^9^ (Methods, Supplementary Table 1). In *Fraxinus baroniana*, *F. chinensis*, *F. floribunda*, *F. mandshurica*, *F. platypoda* and *Fraxinus* sp. D2006-0159, least squares means (LSM) of the proportion of host killed larvae (number of larvae killed by tree defence response divided by total larvae entering the tree) were >0.75 (Fig. 1a, Supplementary Table 2) and no surviving larvae reached the last developmental instar (L4; Fig. 1b, Supplementary Table 2), indicating that these species are resistant to EAB. In contrast, all other taxa tested had a LSM proportion of larvae killed of 0.58 or less (Fig. 1a) and had LSM for L4 larvae proportion between 0 and 0.89 (Fig. 1b).

**Figure 1.**
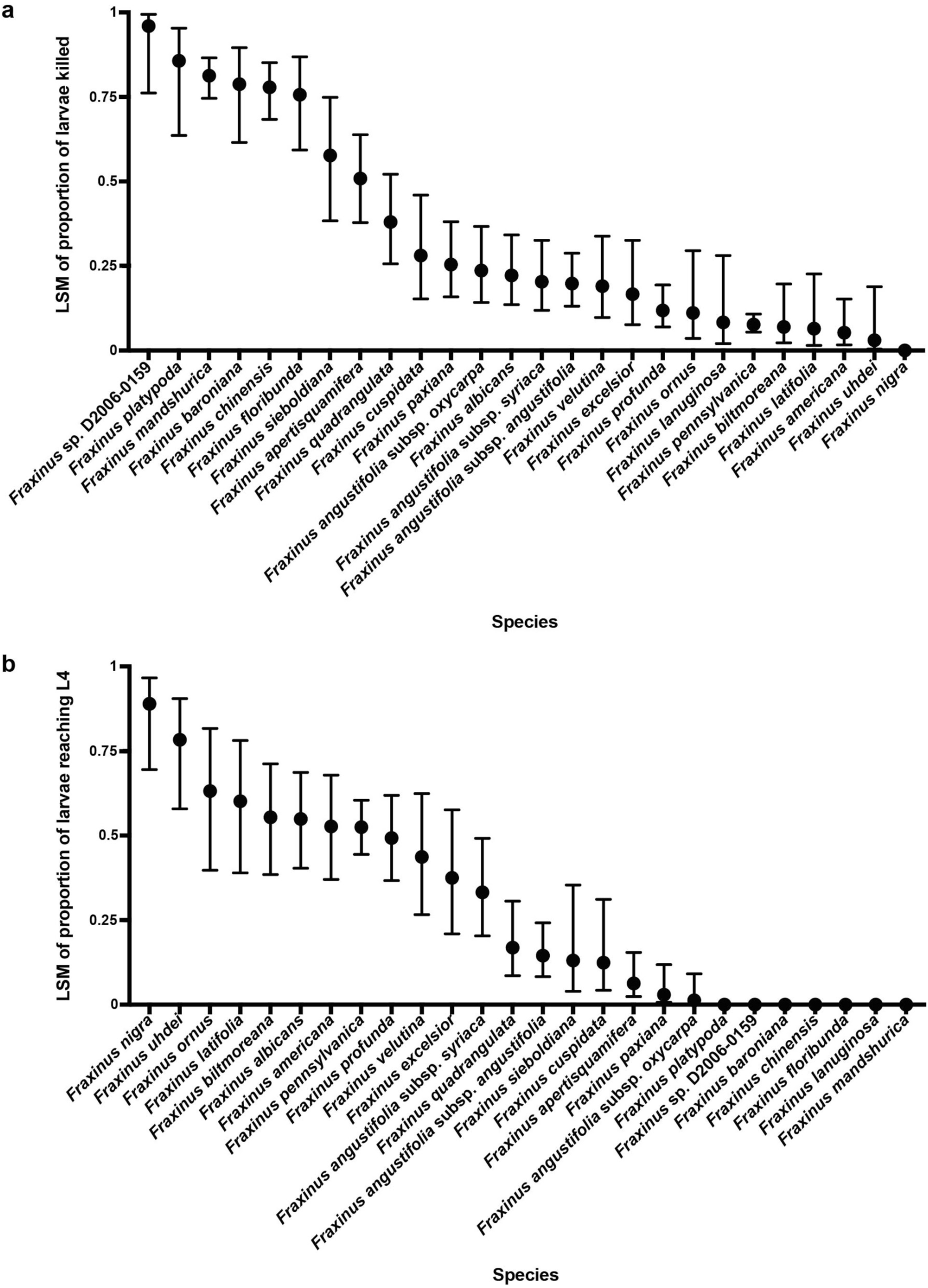
*Fraxinus* species’ resistance to EAB in bioassays. **a, b,** Measures of resistance of different *Fraxinus* taxa to EAB larvae. The *x* axis shows taxa tested; the *y* axis shows least squares means (LSM) of the proportion of larvae successfully entering the tree that were killed by a host defence response (**a**) or LSM of the proportion of larvae successfully entering the tree that reached the L4 instar (**b**). The error bars represent 95% confidence intervals. *Fraxinus* sp. D2006-0159 is a genotype from China for which we could not determine a recognised species name. *Fraxinus biltmoreana*, *F. chinensis*, *F. lanuginosa*, *F. profunda* and *F. uhdei* are polyploids and were not included in the genomic analyses; *F. apertisquamifera* was also not included.

To search for candidate genes related to defence response against EAB, we sequenced and assembled the genomes of 28 individuals from 26 diploid taxa representing all sections within the genus^10^, including a common EAB-susceptible accession and a rare putatively EAB-resistant accession^9^ for *F. pennsylvanica* (Supplementary Table 3). For all individuals we generated c. 35 to 85X whole genome shotgun coverage with Illumina sequencing platforms (Methods; Supplementary Table 4). On assembly (Methods) these data generated 133,719 to 715,871 scaffolds for each individual, with N50s ranging from 1,987 to 50,545bp (Supplementary Table 4). We annotated genes in these assemblies via a reference based approach (Methods) using the published genome annotation of *F. excelsior*^11^. We clustered the protein sequences of these genes into putative orthologue groups (OGs; Methods), also including protein sequences from the *F. excelsior* reference genome and the published genome annotations of *Olea europaea*^12^, *Erythranthe guttata*^13^ and *Solanum lycopersicum*^14^. We found a total of 87,194 OGs, each containing sequences from between two and 32 taxa; 1,403 OGs included a sequence from all 32 taxa.

We generated multiple sequence alignments for the 1,403 OGs including all taxa and inferred gene-trees for each (Methods). In order to generate a species-tree estimate for the genus *Fraxinus*, we conducted Bayesian concordance analysis (Methods). This resulted in a tree based on 272 phylogenetically informative low copy genes (Fig. 2 and Supplementary Note 1). Within this tree, the EAB-resistant taxa identified from our bioassays occurred in three independent lineages. (1) *F. baroniana*, *F. chinensis*, *F. floribunda* and *Fraxinus* sp. D2006-0159 clustered together, within a larger clade that included most species in section *Ornus*, including susceptible *F. ornus*. (2) *F. mandshurica* occurred within a clade corresponding to section *Fraxinus* that also included susceptible taxa. (3) *F. platypoda* was sister to a clade corresponding to section *Melioides*, which includes most of the susceptible American species.

**Figure 2.**
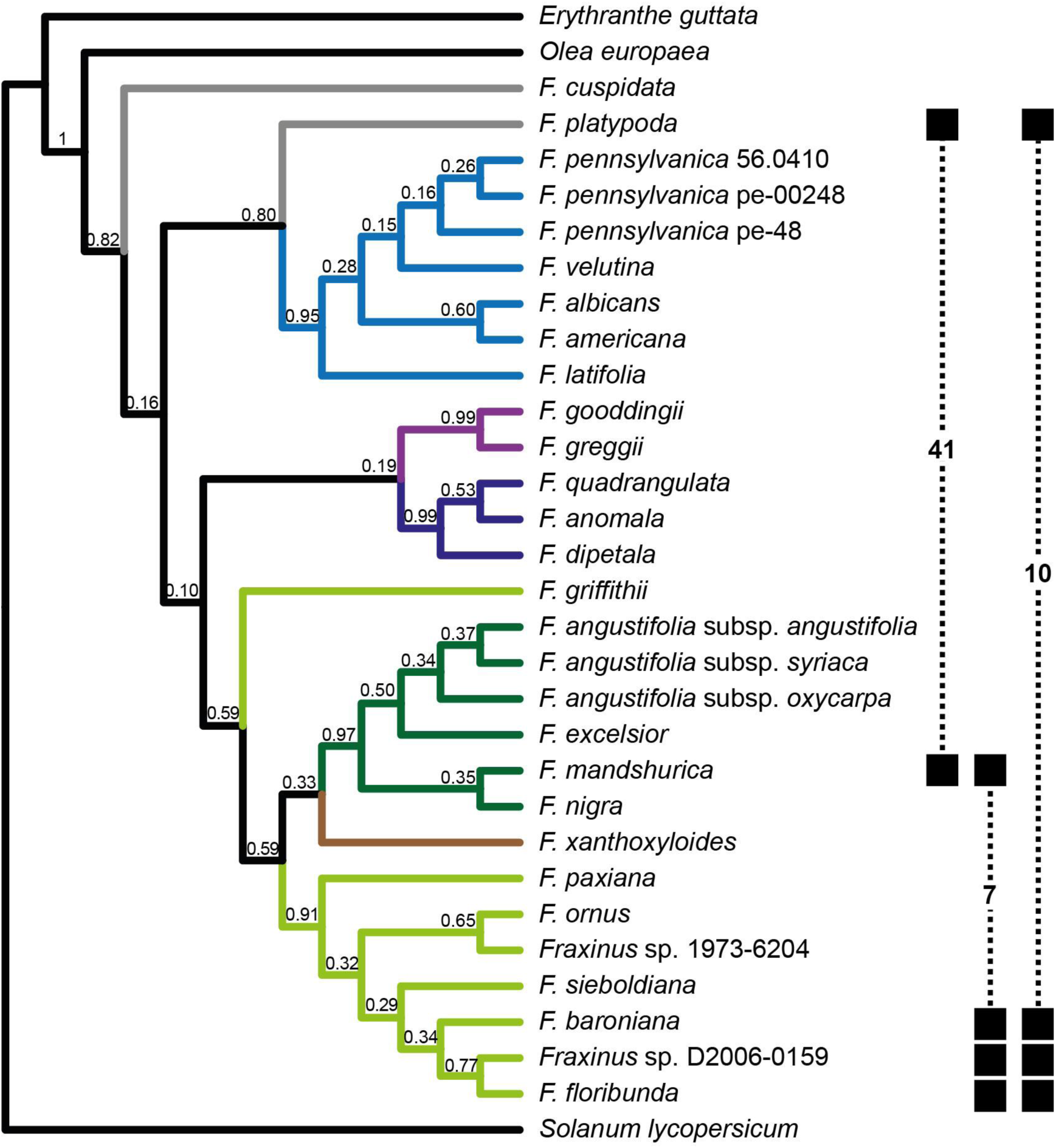
Species-tree for the genus *Fraxinus*. Primary concordance tree inferred from 272 phylogenetically informative loci found in all taxa, inferred via Bayesian concordance analysis with BUCKy. Taxonomic sections within *Fraxinus*, according to Wallander^10^, are shown in different colours: dark blue - section *Dipetalae*; dark green - section *Fraxinus*; light blue - section *Melioides*; light green - section *Ornus*; purple - section *Pauciflorae*; brown - section *Sciadanthus*. *Fraxinus* species not placed into a specific section (*incertae sedis*) are coloured grey and outgroups black. Numbers above the branches are sample-wide concordance factors. Filled squares (linked by dashed lines) indicate the resistant taxa included in the three pairwise convergence analyses, with the number of candidate genes found from that comparison shown; numbers do not sum to 53 (i.e. the total number of candidate genes) because some genes were identified by more than one pairwise comparison.

We searched for amino acid variants putatively convergent between the resistant lineages using an approach that identifies loci with a level of convergence in excess of that likely to be due to chance alone (grand-conv; see Methods). We conducted three pairwise analyses of lineages: (1) *F. mandshurica* versus *F. platypoda*, (2) *F. mandshurica* versus *F. baroniana*, *F. floribunda* and *Fraxinus* sp. D2006-0159, (3) *F. platypoda* versus *F. baroniana*, *F. floribunda* and *Fraxinus* sp. D2006-0159. In all these analyses we included three outgroups and five *Fraxinus* species with high susceptibility (Methods). Each analysis was based on alignments of OGs found in all of the included taxa: 3,454 OGs in analysis 1, 3,097 OGs in analysis 2 and 3,026 OGs in analysis 3. Our candidate amino acid variants were those identified by grand-conv as convergent (minimum posterior probability of 0.90) within loci predicted to have the highest excess of convergent over divergent substitutions in the resistant lineages (Methods). These loci were then checked for the possible confounding effect of paralogy, as well as gene model and alignment errors, leaving a total of 67 amino acid sites in 53 genes (Supplementary Note 2 and Supplementary Table 5). Phylogenetic analysis of the CDS alignments for the 53 genes revealed that, in all but one case (OG20252; Supplementary Fig. 1a), the pattern of amino acid variation at candidate sites is better explained by a hypothesis of convergent point mutations, rather than introgressive hybridisation or incomplete lineage sorting (Supplementary Fig. 1). For four loci (OG11013, OG20859, OG37870 and OG41448) the state identified as convergent by grand-conv appears more likely to be ancestral within *Fraxinus*, with change occurring in the other direction (i.e. from the “convergent” state identified by grand-conv to the “non-convergent” state; Supplementary Table 5).

Three of the 67 candidate amino acids are predicted to be presence/absence variants for phosphorylation sites (Supplementary Fig. 2a, 2b and Supplementary Table 5), a post-translational modification that plays a key role in regulating plant immune signalling^15^. We also looked for evidence of loss-of-function of the 53 candidate genes, based on the presence of frameshifts, stop codon gains and start codon losses, in any of the *Fraxinus* individuals included in our convergence analyses. Six of our 53 candidate genes appear to lack a fully functional allele in a susceptible taxon, compared with one for resistant taxa (Supplementary Note 3 and Supplementary Table 6), suggesting these susceptible taxa may have impaired function of some genes related to defence against EAB.

Among our 53 candidate genes, seven have putative roles relating to the phenylpropanoid biosynthesis pathway (Supplementary Note 4). This pathway generates antifeedant and cytotoxic compounds, as well as products involved in structural defence, such as lignin^16^; it can contribute to indirect defence by producing volatiles which attract parasitoids or predators^17^. Loci OG15551, OG853 and OG16673 are of particular interest. Four convergent amino acids were identified in OG15551 (Fig. 3), a paralogue of *CYP98A3* (Supplementary Fig. 3a), which encodes a critical phenylpropanoid pathway enzyme^18^. Three of the four residues fall within CYP98A3 putative substrate recognition sites, with two at positions predicted to contact the substrate^19^ including a leucine (sulphur containing)/methionine (non-sulphur containing) variant (Fig. 3). OG853 is apparently orthologous to *MED5a*/*RFR1* (Supplementary Fig. 3b), a known regulator of the phenylpropanoid pathway^20,21^ that seems to be involved in regulation of defence response genes^22^. OG16673 is a likely glycoside hydrolase; putative *Arabidopsis thaliana* homologues belong to glycoside hydrolase family 1 and have beta-glucosidase activity, with functions such as chemical defence against herbivory, lignification and control of phytohormone levels^23^. A role for beta-glucosidases in defence against EAB in individual *Fraxinus* species has been previously suggested on the basis of chemical^24^ and transcriptomic^25^ data, and several metabolomic studies have indicated that products of the phenylpropanoid pathway could be involved^26^.

**Figure 3.**
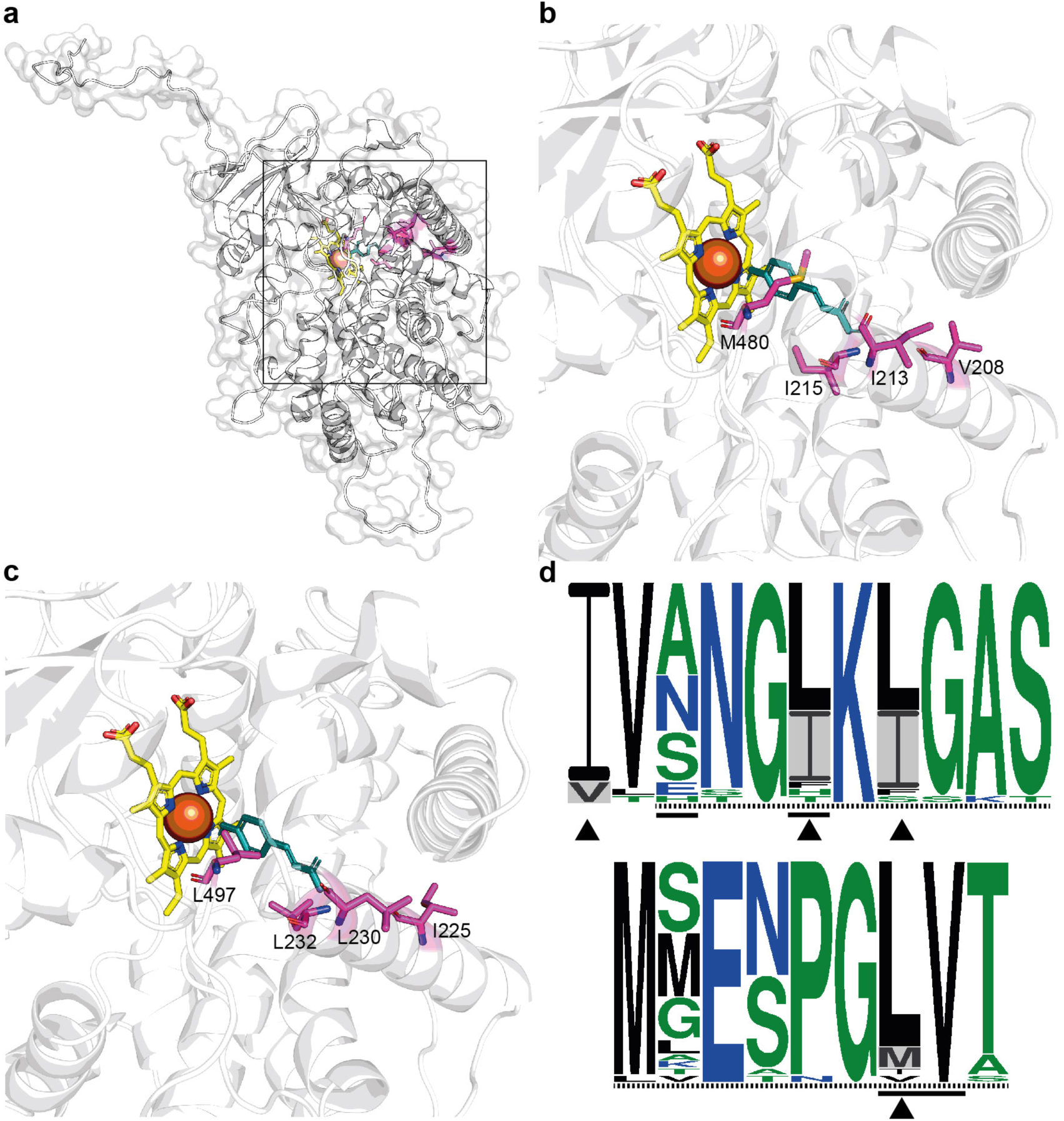
Predicted protein structure for OG15551. **a**, Predicted structure for OG15551, modelled using the protein sequence for the EAB-resistant species *Fraxinus mandshurica*. The black box indicates the region containing the active site, which is enlarged in **b** and **c**. **b**, Region containing the predicted active site in *F. mandshurica*, showing the four amino acid sites at which evidence for convergence between EAB-resistant species was detected. The putative substrate, p-Coumarate, is shown in blue and the heme cofactor in yellow. **c**, Region containing the predicted active site in the EAB-susceptible *F. pennsylvanica* pe-48, showing the amino acid states found at the four sites at which evidence for convergence between EAB-resistant species was detected; the putative substrate and cofactor are shown as in **b**. **d**, Sequence logos for OG15551 and putatively homologous sequences from other angiosperms for regions containing sites at which evidence of convergence was detected (positions 208-218 (top) and 474-482 (bottom) in the *F. excelsior* reference protein), showing the degree of sequence conservation across 30 genera. The height of each residue indicates its relative frequency at that site; amino acids are coloured according to their hydrophobicity (blue = hydrophilic; black = hydrophobic; green = neutral). Dashed lines indicate substrate recognition sites and solid lines residues that are predicted to contact the substrate in the *A. thaliana* CYP98A3 protein^19^; arrowheads indicate sites at which evidence of convergence between EAB-resistant taxa was detected and grey shading shows the amino acid states associated with resistance.

We found 15 candidate genes (Supplementary Note 4) with possible roles in perception and signalling relevant to defence response against herbivorous insects^17^. OG4469 is a probable orthologue of AtG-LecRK-1.6, a G-type lectin receptor kinase (LecRK) with ATP binding activity (Supplementary Note 4.3). G-type LecRKs can act as pattern recognition receptors (PRRs) in the perception of feeding insects^27^; extracellular ATP is a damage-associated molecular pattern (DAMP) whose perception can trigger defence response related genes^27^. OG38407 appears orthologous to *SNIPER4* (Supplementary Fig. 1), a F-box protein encoding gene involved in regulating turnover of defence-response related proteins, for optimal defence activation^28^; the convergent site is in a leucine rich repeat (LRR) region (Supplementary Fig. 2c), which is involved in recognition of substrate proteins for ubiquitination^28,29^.

Several genes appear to relate to phytohormone biosynthesis and signalling, including those with putative functions in jasmonate (JA; OG41448), brassinosteroid (OG43828), cytokinin (OG39275) and abscisic acid (ABA; OG47560) biosynthesis, and GO terms associated with hormone metabolism and biosynthesis are significantly enriched among our set of candidate genes (Supplementary Note 5; Supplementary Table 7). JA signalling is the central regulatory pathway for defence response against insect herbivores^27,30^ whereas brassinosteroids and cytokinins can play important roles in insect resistance, via modulation of the JA pathway^30,31^. ABA is induced by herbivory and is a known modulator of resistance to insect herbivores^27,30^. OG11720 is putatively orthologous to *NRT1.5*/*NPF7.3*., a member of the NRT1/PTR family^32^ which is involved in transport of multiple phytohormones (Supplementary Note 4.3); a transcript matching this gene family had decreased expression in response to both mechanical wounding and EAB feeding in *F. pennsylvanica*^25^. Putative functions of further candidates relate to other signalling molecules involved in triggering defence response (Supplementary Note 4.5), including calcium (OG50989)^17,33^, nitric oxide (NO; OG21033)^34,35^ and spermine (OG33348)^36^. Increased resistance to EAB can be artificially induced in *Fraxinus* species with otherwise high susceptibility^37^, leading to the suggestion that susceptible species may fail to recognise, or respond quickly enough to, early signs of EAB attack^26^. Our identification of candidate genes putatively involved in perception and signalling underlines the possibility of differences between EAB-resistant and susceptible *Fraxinus* species in both their ability to sense and react to attacking insects.

Hypersensitive response (HR), involving programmed cell death (PCD), is associated with effector-triggered immunity in response to microbial pathogens^38^ but can also be induced by insect herbivory^39^ and oviposition^40^. OG16739 and OG37870 are candidates with putative roles related to HR-like effects and PCD. OG16739 has homologues that control cell death in response to wounding, via the induction of ethylene and the expression of defence and senescence related genes^41^. OG37870 may be orthologous to genes that seem to play a role in controlling PCD of xylem elements^42^. Candidate loci whose putative functions lack an obvious link to plant defence response (Supplementary Note 4), could be involved in other phenotypic traits shared between EAB resistant species or may play a role in defence response that is not yet understood. We found that 19 of our 53 candidates match the same *A. thaliana* genes as transcripts that are differentially expressed in response to elm leaf beetle (either in response to simulated egg deposition, or larval feeding)^43^, including genes, such as OG24969, whose putative *A. thaliana* homologues lack a clear defence-related function.

We analysed allelic variation at the 67 amino acid sites within the 53 candidate genes for all sequenced taxa assessed for resistance to EAB. Of the 67 sites, seven only have the EAB-resistance associated state in resistant taxa, and another is only homozygous for the EAB-resistance associated state in resistant taxa (Supplementary Table 8). Of the 53 candidate genes, four are only homozygous in resistant taxa for the EAB-resistance associated state at the candidate amino acid site(s) detected within them (OG853, OG21449, OG36502 and OG37560; Supplementary Table 8). If we omit the genomes of *F. nigra*, *F. excelsior* and the three *F. angustifolia* subspecies (sect. *Fraxinus*), for 24 of the 53 candidate genes we only find the EAB-resistance associated states in resistant taxa, for 48 genes they are only found in taxa with a LS mean proportion of larvae killed of ≥0.25 and the remaining five genes are only homozygous for the EAB-resistance associated states in taxa with a LS mean proportion of larvae killed of ≥0.25 (Supplementary Table 8).

Analysis of previously generated whole genome sequence data for 37 *F. excelsior* individuals from different European provenances revealed that for 50 of the 67 candidate amino acid sites (occurring in 41 of the 53 genes) the EAB-resistance associated state was present, with evidence for polymorphism at seven of these sites (Supplementary Table 9). None of the EAB-resistance associated amino acid states were found in the putatively resistant *F. pennsylvanica* genotype (Supplementary Table 8), suggesting that different genes, or different variants within these genes, are involved in the intraspecific variation in susceptibility of this species. Despite this, transcripts inferred to be from 11 of our candidate genes showed evidence for differential expression subsequent to EAB-feeding in *F. pennsylvanica* (Supplementary Note 6 and Supplementary Table 10) and two gene families that were highlighted as potentially important for response to tissue damage in *F. pennsylvanica*^25^ are also represented among our candidates (see above).

We have provided the first evidence based on genome-wide analyses for the types of genes involved in resistance to EAB in the genus *Fraxinus*, indicating that multiple loci, contributing to different defence responses, underlie this trait. Our genome level data may help to target future efforts to increase the resistance of American and European ash species to EAB via breeding or gene editing.

## METHODS

### Data reporting

For the emerald ash borer resistance assays, experiments were conducted using a randomised block design. No statistical methods were used to predetermine sample size. The investigators were not blinded to allocation during experiments and outcome assessment.

### Plant material

All plant materials used in this study were sourced from living or seed collections in the UK or USA. Due to biosecurity measures, we were not able to move living materials between the two countries. In our initial selection of material we relied upon species identifications that had already been made in the arboreta or seed banks within which the materials were held. For each of the accessions included in this study, we PCR amplified and Sanger sequenced the nuclear ribosomal internal transcribed spacer (ITS) region, following standard methods; forward and reverse sequences were assembled into contigs using CLC Genomics Workbench v8.5.1 (QIAGEN Aarhus, Denmark). As far as possible the identity of all materials was verified by Eva Wallander using morphology and ITS sequences, according to her classification of the genus^10^. This led to some re-designation of samples: of particular note, one of the three accessions of *F. pennsylvanica* that we genome sequenced was originally sampled as *F. caroliniana*, and the accession that we originally sampled and genome sequenced as *F. bungeana* was determined to be *F. ornus* (Supplementary Table 3). However, subsequent phylogenetic analysis including allele sequences for this latter individual (see below - Distinguishing between different underlying causes of convergent patterns) indicated it to likely be a hybrid between the *F. ornus* lineage and another lineage within sect. *Ornus* and therefore we designate it as *Fraxinus* sp. 1973-6204. We also designated an accession that was originally sampled for genome sequencing as *F. chinensis* as *Fraxinus* sp. D2006-0159 due to uncertainty regarding species delimitation. Furthermore, for genotype vel-4, that was redetermined as *F. pennsylvanica*, we have maintained its original species name (*F. velutina*). A list of all materials used in the study is shown in Supplementary Table 3, including initial identifications and subsequent identifications by Eva Wallander, as well as details of voucher specimens.

### Emerald ash borer resistance assays

Twenty six *Fraxinus* taxa (species, subspecies and one taxon of uncertain status) were collected for egg bioassay experiments (Supplementary Table 3). We aimed to test three clonal replicates (grafts or cuttings) of at least two genotypes of each species. For some taxa less than two genotypes were available in the US, and occasionally genotypes did not propagate well by graft or cutting so seedlings from the same seedlot were used instead (details for each taxon are included in Supplementary Table 1). The majority of egg bioassays were conducted in 2015 and 2016 and groups of approximately 20 genotypes were conducted in each set (week within year; Supplementary Table 1) with one grafted ramet, cutting or seedling in each block and all ramets, cuttings and seedlings within the block randomised to location/order of assay. To facilitate comprehensive analysis, the same controls were repeated in each week (susceptible *F. pennsylvanica* genotype pe-37 and/or pe-39 and resistant *F. mandshurica* genotype ‘mancana’).

Trees were treated as uniformly as possible prior to inoculation. Adult beetles were reared and used for egg production as previously described^9^. Inoculations were performed in a greenhouse to keep conditions uniform for the duration of the assay, and to minimise predation of the eggs. We followed the EAB egg transfer bioassay method reported by Koch et al.,^9^ that had previously been used on genotypes of *F. pennsylvanica* and *F. mandshurica* with the changes noted below. The egg dose for each tree was determined according to the method of Duan et al.,^44^, which takes into account the bark surface area. A target density of 400 eggs per m^2^ was used; this density is above that reported to allow host defences to kill larvae in green ash, but is in the range where competition and cannibalism are minimised^44^. Twelve individual eggs, on a small strip cut from the coffee filter paper on which they were laid, were taped to each tree. The spacing was varied between eggs to maintain a consistent target dose (e.g. eggs placed 7.5 cm apart on stem 1.0-1.1cm diameter to eggs placed 3 cm apart on stem 2.5-2.6cm diameter). The portion of the tree where eggs were placed was wrapped in medical gauze to protect from jostling and egg predation. Past experiments have shown that egg assay results are not consistent on stems less than 1 cm in diameter (JLK, unpublished data). Due to size differences between some species, to achieve the target dose and avoid placing eggs where the stem diameter was <1 cm, occasionally less than 12 eggs were placed. A total of 2199 egg bioassays (each egg represents a bioassay) were conducted on 61 different genotypes and a total of 206 ramets, cuttings or seedlings.

Occasional ramets, cuttings and/or seedlings were considered as assay failures if less than three larvae successfully entered the tree (i.e. the effective egg dose was too low), or if there were other problems with the tree (too small diameter overall, cultivation issues, etc.), and that replicate was excluded from analysis (data not shown). Four weeks after egg attachment each egg was inspected to determine if it had successfully hatched, and if there were signs of the larva entering the tree. Larval entry holes, when detected, were marked to assist with future dissection. At eight weeks, dissection of the entry site was performed and galleries made by larval feeding were carefully traced using a grafting knife to determine the outcome of each hatched egg. Health (dead or alive) and weight (in cases when larvae could be recovered intact) was recorded for each larva, and developmental instar was determined using measurements of head capsule and length^45,46^

Preliminary exploratory data analysis indicated that the proportion of “tree killed” (i.e. larvae killed by tree defence response) and the proportion of live L4 larvae (number divided by the number of larvae that entered the tree) were the best variables to distinguish resistance versus susceptibility at the species level. We fitted a generalised linear mixed model to the proportion tree killed and proportion L4 using the GLIMMIX procedure in SAS. The model specification is proportion as a binomial distribution with a logit link function, species as a fixed effect, and block/replicate nested within sequential week (week within year) as a random effect (this allowed for comprehensive analysis over years and weeks with correct variance/covariance restrictions). Non-significant predictors were eliminated from the final model. Least squares means of tree killed or L4 proportion were calculated with confidence intervals on the data scale (proportion).

### Genome size estimation by flow cytometry (FC)

We used FC to estimate the genome size of individuals used for whole genome sequencing (Supplementary Table 4). *Fraxinus* samples from UK collections were prepared and analysed as described in Pellicer et al. (2014), with the exception that ‘general purpose isolation buffer’ (GPB^47^) without the addition of 3% polyvinylpyrrolidone (PVP-40) and LB01 buffer^48^ were used for some samples. *Oryza sativa* (‘IR-36’; 1C = 0.50 pg^49^) was used as an internal standard. For each individual analysed, two samples were prepared (from separate leaves or different parts of the same leaf) and two replicates of each sample run. *Fraxinus* samples from US collections were analysed using a Sysmex CyFlow Space flow cytometer, as described in Whittemore and Xia^50^*; Pisum sativum* (‘Ctirad’; 1C = 4.54 pg^51^) and *Glycine max* (‘Williams 82’; 1C = 1.13 pg^52^) were used as internal standards. For each individual analysed, six samples were prepared (from separate leaves or different parts of the same leaf) and three samples run with each size standard.

### DNA extraction

Total genomic DNA was extracted from fresh, frozen or silica-dried leaf or cambial material, using either a CTAB extraction protocol modified from^53^ or using a Qiagen DNeasy^®^ Plant Mini or Maxi kit.

### Genome sequencing and assembly

For each of the 28 diploid individuals selected for whole genome sequencing (Supplementary Table 3), sufficient Illumina sequence data were generated to provide a minimum of c. 30x coverage of the 1C genome size, based on the C-value estimates obtained for the same individuals (see above - Genome size estimation by flow cytometry (FC)), or those of closely related taxa. Libraries with average insert sizes of 300 or 350bp, 500 or 550bp and 800bp were prepared from total genomic DNA by the Genome Centre, at Queen Mary University of London, and the Centre for Genomic Research, at the University of Liverpool. Paired-end reads of 125, 150 or 151 nucleotides were generated using the Illumina NextSeq 500, HiSeq 2500 and HiSeq 4000 platforms (Illumina, San Diego, California, USA); see Supplementary Table 4 for the exact combination of libraries, read lengths and sequencing platforms used for each individual. For selected taxa, chosen to represent different sections within the genus, we also generated data from long mate-pair (LMP) libraries (Supplementary Table 4). LMP libraries with average insert sizes of 3kb and 10kb were prepared from total genomic DNA by the Centre for Genomic Research, at the University of Liverpool, and sequenced on an Illumina HiSeq 2500 to generate reads of 125 nucleotides to a depth of c. 10x coverage of the 1C genome size.

Initial assessment of sequence quality was performed for all read pairs from the short-insert libraries (300-800bp inserts) using FastQC v.0.11.3 or v.0.11.5 (www.bioinformatics.babraham.ac.uk/projects/fastqc/). Reads were clipped using the fastx_trimmer tool in the FASTX-Toolkit v.0.0.14 (http://hannonlab.cshl.edu/fastx_toolkit/index.html) to remove the first 5-10 nucleotides of each read; for the NextSeq reads, the last 5 nucleotides were also clipped. Adapter trimming was performed using cutadapt v.1.8.1^54^ with the “O” parameter set to 5 (i.e. minimum overlap of five bases) and using option “b” (i.e. adapters allowed to occur on both the 5’ and 3’ end of reads); default settings were used for all other parameters. Quality trimming and length filtering was performed using Sickle v.1.33^55^ with the “pe” option and the following parameter settings: -t sanger -q 20 −l 50 and default settings for other parameters. This yielded quality trimmed paired and singleton reads with a minimum length of 50 nucleotides; only intact read pairs were used for downstream analyses.

For the LMP libraries, duplicate reads were removed using NextClip v.1.3.1 (https://github.com/richardmleggett/nextclip) with the --remove_duplicates parameter specified and default settings for all other parameters. Adapter trimming was performed using cutadapt v.1.10^54^; junction adapters were removed from the start of reads by running option “g”, with the adapter sequence anchored to the beginning of reads with the “^” character, and the following settings for other parameters: -O 10 -n 2 -m 25. Other adapter trimming was performed using option “a”, with the further parameters set to the same values as specified above. Quality trimming was performed with PRINSEQ-lite v.0.20.4^56^, with the following parameter settings: -trim_qual_left 20 -trim_qual_right 20 -trim_qual_window 20 -trim_tail_left 101 -trim_tail_right 101 -trim_ns_left 1 - trim_ns_right 1 -min_len 25 -min_qual_mean 20 -out_format 3.

*De novo* genome assembly was performed for each individual using CLC Genomics Workbench v8.5.1 (QIAGEN Aarhus, Denmark). All trimmed read pairs from the short-insert libraries were used for assembly under the following parameter settings: automatic optimization of word (k-mer) size; maximum size of bubble to try to resolve=5000; minimum contig length=200bp. Assembled contigs were joined to form scaffolds using SSPACE (version 3.0^57^) with default parameters, incorporating data from mate-pair libraries with 3kb and 10kb insert sizes where available. Library insert lengths were specified with a broad error range (i.e. ±40%). Gaps in the SSPACE scaffolds were filled using GapCloser (version 1.12^58^) with default parameters. The average library insert lengths were specified using the estimates produced by SSPACE during scaffolding. Scaffolding and gap filling was not performed for individuals that lacked data from libraries with insert size of ≥500bp (only a single insert size library was available for these taxa; Supplementary Table 4). We did not attempt to extract sequences of organellar origin from the assemblies, or to separately assemble the plastid and mitochondrial genomes.

Sequences within the assemblies that correspond to the Illumina PhiX control library were identified via BLAST. A PhiX bacteriophage reference sequence (GenBank accession number CP004084) was used as a query for BLASTN searches, implemented with the BLAST+ package (v.2.5.0+^59^), against the genome assembly for each taxon with an *E* value cut-off of 1×10^−10^. Sequences that matched the PhiX reference sequence at this threshold were removed from the assemblies. We used the assemblathon_stats.pl script (https://github.com/ucdavis-bioinformatics/assemblathon2-analysis/blob/master/assemblathon_stats.pl) with default settings to obtain standard genome assembly metrics, such as N50. BUSCO v2.0^60^ was used to assess the content of the genome assemblies. The “embryophyta_odb9“ lineage was used and analyses run with the following parameter settings: --mode genome -c 8 -e 1e-05 -sp tomato.

### Gene annotation and orthologue inference

To annotate genes in the newly assembled *Fraxinus* genomes, we used a similarity based approach implemented in GeMoMa^61^, with genes predicted in the *F. excelsior* BATG0.5 assembly as a reference set. We used the “Full Annotation” gff file for BATG0.5 (Fraxinus_excelsior_38873_TGAC_v2.longestCDStranscript.gff3; available from http://www.ashgenome.org/transcriptomes), which contains the annotation for the single longest splice variant for each gene model. This annotation file also includes preliminary annotations for genes within the organellar sequences (gene models FRAEX38873_v2_000400370-FRAEX38873_v2_000401330) which were not reported in the publication of the reference genome^11^; none of the sets of putative orthologues used for the species-tree inference or molecular convergence analysis (see below) include these preliminary organellar models from the BATG0.5 reference assembly. The Extractor tool from GeMoMa v.1.3.2 was used to format the data from the reference genome (gff and assembly files), with the following parameter settings: v=true f=false r=true Ambiguity=AMBIGUOUS. To obtain information on similarity between the reference gene models and sequences in the newly assembled *Fraxinus* genomes, we performed TBLASTN searches of individual exons (i.e. the “cds-parts” file generated by Extractor) against the assembly file for each individual with BLAST+ (v.2.2.29+^59^). makeblastdb was used to format each assembly file into a BLAST database with the following parameter settings: -out ./blastdb -hash_index -dbtype nucl. tblastn was then run with the “cds-parts.fasta” file as the query, with the following parameter settings: - num_threads 24 -db ./blastdb -evalue 1e-5 -outfmt “6 std sallseqid score nident positive gaps ppos qframe sframe qseq sseq qlen slen salltitles” -db_gencode 1 -matrix BLOSUM62 -seg no -word_size 3 -comp_based_stats F -gapopen 11 -gapextend 1 - max_hsps 0. Finally, the GeMoMa tool itself was run for each individual, with the TBLASTN output, cds-parts file and *de novo* assembly file as input, with the “e” parameter set to “1e-5” and default settings for all other parameters. Because GeMoMa generates predictions for each reference gene model separately, the output may contain gene models that are at identical, or overlapping, positions, especially for genes that belong to multi-gene families (http://www.jstacs.de/index.php/GeMoMa#FAQs). As the presence of these redundant gene models does not prevent the correct inference of sets of orthologues with OMA (see below), we opted to retain the predicted proteins from all gene model predictions generated by GeMoMa for input into OMA. The gffread utility from cufflinks v.2.2.1^62^ was used to generate the CDS for each gene model; getfasta from bedtools v.2.26.0^63^ used to generate full-length gene sequences (i.e. including introns where present), with the “-name” and “-s” options invoked.

To identify sets of putatively orthologous sequences, we used OMA standalone v.2.0.0^64,65^ to infer OMA groups (OGs) and hierarchical orthologous groups (HOGs). To the protein sets from the 29 diploid *Fraxinus* genome assemblies (the 28 newly generated assemblies, plus the existing reference assembly for *F. excelsior*), we added proteomes from three outgroup species: *Olea europaea* (olive), which belongs to the same family as *Fraxinus* (Oleaceae); *Erythranthe guttata* (monkey flower; formerly known as *Mimulus guttatus*), which belongs to the same order as *Fraxinus* (Lamiales); *Solanum lycopersicum* (tomato), which belongs to the same major eudicot clade as *Fraxinus* (lamiids). For *O. europaea*, we used the annotation for v6 of the genome assembly^12^; the file containing proteins for the single longest transcript per gene (OE6A.longestpeptide.fa) was downloaded from: http://denovo.cnag.cat/genomes/olive/download/?directory=.%2FOe6%2F. For *E. guttata* we used the annotation for v2.0 of the genome assembly^13^; the file containing proteins for the primary transcript per gene (Mguttatus_256_v2.0.protein_primaryTranscriptOnly.fa) was downloaded from Phytozome 12^66^. For *S. lycopersicum* we used the annotation for the vITAG2.4 genome assembly; the file containing proteins for the primary transcript per gene (Slycopersicum_390_ITAG2.4.protein_primaryTranscriptOnly.fa) was downloaded from Phytozome 12.

Fasta formatted files containing the protein sequences from all 32 taxa were used to generate an OMA formatted database. An initial run of OMA was performed using the option to estimate the species-tree from the OGs (option ‘estimate’ for the SpeciesTree parameter); we set the InputDataType parameter to ‘AA’ and left all other parameters with the default settings. The species-tree topology from the initial run was then modified in FigTree v.1.4.2 (http://tree.bio.ed.ac.uk/software/figtree/) to reroot it on *S. lycopersicum*; nodes within the main clades of *Fraxinus* species (which corresponded to sections recognised in the taxonomic classification^10^) were also collapsed and relationships between these major clades and any individual *Fraxinus* not placed into a clade, were collapsed. OMA was then rerun with the modified species-tree topology specified in Newick format through the SpeciesTree parameter; the species-tree topology is used during the inference of HOGs (https://omabrowser.org/standalone/), and does not influence the OGs obtained.

### Species-tree inference

To obtain a more robust estimate of the species-tree for *Fraxinus* (compared with that estimated by OMA, see above, or existing species-tree estimates based on very few independent loci^67,68^), we selected clusters of putatively orthologous sequences from the results of the OMA analysis. OGs containing protein sequences from all 32 taxa (29 diploid *Fraxinus* and three outgroups) were identified and corresponding CDSs aligned with MUSCLE^69^ via GUIDANCE2^70^ with the following parameter settings: --program GUIDANCE --msaProgram MUSCLE --seqType codon --bootstraps 100, and default settings for other parameters. Datasets where sequences were removed during the alignment process (identified by GUIDANCE2 as being unreliably aligned) or which failed to align due to the presence of incomplete codons (i.e. the sequence length was not divisible by three) were discarded. Alignment files with unreliably aligned codons removed (i.e. including only codons with GUIDANCE scores above the 0.93 threshold for the “colCutoff” parameter) were used for downstream analyses. A custom Perl script was used to identify alignments shorter than 300 characters in length or which included sequences with <10% non-gap characters; these datasets were excluded from further analysis.

The remaining alignment files were converted from fasta to nexus format with the seqret tool from EMBOSS v.6.6.0^71^. MrBayes v. 3.2.6^72^ was used to estimate gene-trees with the following parameter settings: lset nst=mixed rates=gamma; prset statefreqpr=dirichlet(1,1,1,1); mcmc nruns = 2 nchains = 4 ngen = 5000000 samplefreq = 1000; Sumt Burninfrac = 0.10 Contype = Allcompat; Sump Burninfrac = 0.10. Diagnostics (average standard deviation of split frequencies (ASDSF) and for post burnin samples, potential scale reduction factor (PSRF) for branch and node parameters and effective sample size (ESS) for tree length) were examined to ensure that runs for a given locus had reached convergence and that a sufficient number of independent samples had been taken. We discarded datasets where the ASDSF was ≥0.010; all remaining datasets had an ESS for tree length in excess of 500.

We used BUCKy v.1.4.4^73,74^ to infer a species-tree for *Fraxinus* via Bayesian concordance analysis, which allows for the possibility of gene-tree heterogeneity (arising from biological processes such as hybridisation, that is reported to occur within *Fraxinus*^67^) but which makes no assumptions regarding the reason for discordance between different genes^74^. First, mbsum v.1.4.4 (distributed with BUCKy) was run on the MrBayes tree files for each locus, removing the trees sampled during the first 500,000 generations of each run as a burn-in. We then used the output from mbsum to select the most informative loci on the basis of the number of distinct tree topologies represented in the sample of trees from MrBayes; loci with a maximum of 2000 distinct topologies were retained. BUCKy was then run on the combined set of mbsum output files for all retained loci under ten different values for the α parameter (0.1, 1, 2, 5, 10, 20, 100, 500, 1000, ∞), which specifies the *a priori* level of discordance expected between loci. Each analysis with a different α setting was performed with different random seed numbers (parameters -s1 and -s2) and the following other parameter settings: -k 4 -m 10 -n 1000000 -c 2. Run outputs were checked to ensure that the average standard deviation of the mean sample-wide concordance factors (CF; the sample-wide CF is the proportion of loci in the sample with that clade^73^) was <0.01. The same primary concordance tree (PCT; the PCT comprises compatible clades found in the highest proportion of loci and represents the main vertical phylogenetic signal^74^) and CFs for each node were obtained for all settings of α.

We also repeated the species-tree inference using full-length sequences (i.e. including introns, where present); alignment and gene-tree inference were carried out as described above, with the exception that the --seqType parameter in GUIDANCE2 was set to “nuc”. BUCKy analyses were performed as described above. The same PCT was obtained for all settings of α, with only minor differences in the mean sample-wide concordance factors. The PCT inferred from the full-length datasets was also identical to that obtained from the CDS analyses; we based our final species-tree estimate on the output of the full-length analyses due to the presence of a larger number of informative loci within these datasets.

One of the informative loci from the CDS analyses (i.e. those with ≤2000 distinct topologies within their gene-tree sample), and three of those from the full-length analyses, were subsequently found to be among our filtered set of candidate loci with evidence of convergence between EAB-resistant taxa (see below). To test whether the signal from these loci had an undue influence on the species-tree estimation, we excluded them and repeated the BUCKy analyses for the CDS and full-length datasets as described above, with the exception that only an α parameter setting of 1 was used. The PCTs obtained from these analyses were identical to those inferred when including all datasets, with minor (i.e. 0.01) differences in CFs.

In addition to the analysis including all taxa, we also performed BUCKy analyses for 13 taxa selected for inclusion in the grand-conv analyses and for the subsets of 10-12 taxa for each of the three grand-conv pairwise comparisons (see below - Analysis of patterns of sequence variation consistent with molecular convergence). OGs containing protein sequences from all 13 taxa (10 *Fraxinus* and three outgroups) were identified and corresponding CDSs for these 13 taxa aligned with MUSCLE via GUIDANCE2, as described above. We also identified and aligned CDSs for all additional OGs which did not include all 13 taxa, but which contained proteins for all taxa in one of the subsets used for the pairwise comparisons. Filtering of alignments and gene-tree inference were carried out as described above for the full set of 32 taxa. BUCKy was run separately with the MrBayes tree sample samples for loci including all 13 taxa and for additional loci including taxa for each of the three smaller subsets for the grand-conv pairwise comparisons. BUCKy analyses were performed as described above, with the exception that no filtering of loci on the basis of number of distinct gene-tree topologies was performed, a value of between 2 and 35 was used for the -m parameter, and only a single α parameter setting, of 0.1, was used. Topologies for the PCTs for the set of 13 taxa and subsets of 10-12 taxa were both congruent with each other and with the PCTs inferred from analyses including all taxa, for all nodes with a CF of ≥0.38.

### Analysis of patterns of sequence variation consistent with molecular convergence

To test for signatures of putative molecular convergence in protein sequences we used a set of diploid taxa representing the extremes of variation in susceptibility to EAB, as assessed by our egg bioassays. By limiting our analysis at this stage to this subset of taxa we could also maximise the number of genes analysed, as with increasing taxon sampling the number of OGs for which all taxa are represented decreases. This set comprised: five highly susceptible taxa (*F. americana*, *F. latifolia*, *F. ornus*, *F. pennsylvanica* [susceptible genotype], and *F. velutina*), five resistant taxa (*F. baroniana, F. floribunda, F. mandshurica*, *F. platypoda* and *Fraxinus* sp. D2006-0159) and three outgroups (*O. europaea, E. guttata* and *S. lycopersicum*).

We did not include *F. nigra* as one of the highly susceptible species, although it has been used in comparisons of resistant and susceptible species in several previous studies (e.g. ^75,76^), because detailed analysis of the phenotype of *F. nigra* individuals using EAB bioassays indicates evidence of a possible widespread hypersensitive-like response, which rather than being a beneficial defence response may actually accelerate tree death (JLK, unpublished data), similar to what has been proposed for *Tsuga canadensis* in response to hemlock woolly adelgid^39^. This is consistent with reports of mortality of *F. nigra* occurring at lower densities of EAB in the field compared with other highly susceptible North American species (N. Siegert and D. McCullough, pers. comm.).

Of the ten *Fraxinus* genome assemblies included in the convergence analysis, six were from genotypes that were also included in the EAB resistance assays outlined above (*F. americana* am-6, *F. baroniana* bar-2, *F. floribunda* flor-ins-12, *F. pennsylvanica* pe-48 and *F. platypoda* spa-1 and *Fraxinus* sp. D2006-0159 F-unk-1). For the other four, we could not test the exact individual that we sequenced the genome of, but relied upon results of bioassays of other individuals in the same species.

We used the following three pairwise comparisons to seek loci showing amino acid convergence between resistant taxa:

1. *F. mandshurica* (sect. *Fraxinus*) versus *F. platypoda* (*incertae sedis*)
2. *F. mandshurica* (sect. *Fraxinus*) versus *F. baroniana*, *F. floribunda* and *Fraxinus* sp. D2006-0159 (sect. *Ornus*)
3. *F. platypoda* (*incertae sedis*) versus *F. baroniana, F. floribunda* and *Fraxinus* sp. D2006-0159 (sect. *Ornus*)

The more divergent homologous amino acid sequences are between species, the more likely it is that a convergent amino acid state will occur by chance^4^. In order to account for this, we compared the posterior expected numbers of convergent versus divergent substitutions across all pairs of independent branches of the *Fraxinus* species-tree for the selected taxa using a beta-release of the software Grand-Convergence v0.8.0 (hereafter referred to as grand-conv; https://github.com/dekoning-lab/grand-conv). This software is based on PAML 4.8^77^ and is a development of a method used by Castoe et al^4^. It has also been recently used, for example, to detect convergence among *Laverania* species^78^.

For input into grand-conv, we used the same OGs that were the basis of the BUCKy analyses of the 13 taxa selected for the grand-conv analyses, and analyses of the subsets of taxa for the pairwise comparisons (see above - Species-tree inference). Therefore, as well as meeting the criterion of including all relevant taxa, the OGs analysed with grand-conv had also successfully passed the alignment, filtering and gene-tree inference steps. A set of input files was created for each of the three pairwise comparisons that, where present, removed any taxa from the other resistant lineage from the alignments generated by GUIDANCE2. Alignment files were then converted from fasta to phylip format using the Fasta2Phylip.pl script [https://github.com/josephhughes/Sequence-manipulation]. To ensure that sequences from each taxon appeared in a consistent order across all datasets (which is necessary for automating the generation of site-specific posterior probabilities for selected branch pairs with grand-conv), the phylip formatted files were sorted using the unix command “sort” prior to input into grand-conv. Species-tree files for each of the pairwise comparisons were created from the Newick formatted PCTs for the relevant taxon sets generated with BUCKy, with the trees edited to root them on *S. lycopersicum*.

For the grand-conv analysis, “gc-estimate” was first run on the full set of input alignment files for each pairwise comparison, with the following parameters settings: --gencode=0 - -aa-model=lg --free-bl=1, specifying the appropriate species-tree file for each of the pairwise comparisons. Next, “gc-discover” was run to generate site-specific values for the posterior probability (PP) of divergence or convergence for the branch pairs of interest; the numbers for the branch pairs relating to the resistant taxa were established from an initial run of “gc-estimate” and “gc-discover” on a single input file, and then specified when running “gc-discover” on all input files using the --branch-pairs parameter. A custom Perl script was then used to filter the output files containing the site-specific posterior probabilities to identify loci with at least one amino acid site where the PP of convergence was higher than divergence and passed a ≥0.9000 threshold. For this filtered set of datasets with significant evidence of convergence at at least one site, we checked if the “excess” convergence, as measured from the residual values from the non-parametric errors-in-variables regression calculated by grand-conv, was higher for the branch pair of interest than for any other independent pair of branches within the species-tree (i.e. the highest residual was found for the resistant branch pair). Only loci where the highest excess convergence was found in the resistant branch-pair were retained for further analysis.

### Refining the initial list of candidate loci identified with grand-conv

For the set of loci with evidence of convergence between at least one pair of resistant lineages from the grand-conv analyses, we applied additional tests to assess the robustness of the pattern of shared amino acid states. Specifically, we checked for the potential impact of alignment uncertainty and orthology/paralogy conflation. For each candidate locus, we identified the Hierarchical Orthologous Group (HOG) from the OMA analysis, to which the sequences for the candidate locus belong. These HOGs include sequences for an expanded set of taxa (see above - Species-tree inference) and may represent a single gene for all taxa (i.e. a set of orthologous sequences) or several closely related paralogues^64^. Protein sequences for HOGs were aligned with GUIDANCE and gene-tree inference conducted with MrBayes, as described above for the OMA putative orthologous groups with the exception that the --seqType parameter in GUIDANCE was set to “aa” and in MrBayes the prset parameter was set to “prset aamodelpr = mixed”. Any MrBayes analyses that had not converged after 5M generations (average standard deviation of split frequencies >0.01) were run for an additional 5M generations. The multiple sequence alignment and gene-tree estimates were then used to refine the initial list of candidate loci. Loci were dropped from initial list of candidates if either of the following applied:

1. If in the filtered MSA alignment generated by GUIDANCE, the site/sites where convergence was detected were not present, indicating they were in a part of the protein sequences that can not be aligned reliably.
2. If in the consensus gene-tree estimated by MrBayes, there was evidence that the sequences within which convergence was initially detected (i.e. those belonging to the ten *Fraxinus* species included in the grand-conv analysis) belong to different paralogues and that the pattern of convergence could be explained by sequences with the “convergent” state belonging to one paralogue and the “non-convergent” state belonging to another paralogue. We also excluded two loci that belong to large gene families (>10 copies) for which the MrBayes analyses failed to reach convergence within a reasonable time (≤10M generations) and for which orthology/paralogy conflation could therefore not be excluded.

Additionally, for the set of loci remaining, we checked for errors in the estimation of gene models (including in the reference models from *F. excelsior*) that might impact the results of the grand-conv analyses. Specifically, we dropped from our list any loci where the amino acid sites with evidence of convergence were found to be outside of an exon, or outside of the gene itself, following manual correction of the gene model prediction.

### Analysis of variants within candidate loci

To assess the possible impact of allelic variants (i.e. those not represented in the genome assemblies) on patterns of amino acid variation associated with the level of EAB susceptibility in *Fraxinus*, we called variants (SNPs and indels) and predicted their functional effects. For each sequenced *Fraxinus* individual, trimmed read pairs from the short insert Illumina libraries were mapped to the *de novo* genome assembly for the same individual using Bowtie 2 v.2.3.0^79^ with the “very-sensitive” preset and setting “maxins” to between and 1000 and 1400, depending on the libraries available for that individual. Read mappings were converted to BAM format and sorted using the “view” and “sort” functions in samtools v.1.4.1^80^. Prior to variant calling, duplicate reads were marked and read group information added to the BAM files using the “MarkDuplicates” and “AddOrReplaceReadGroups” functions in picard tools v.1.139 (http://broadinstitute.github.io/picard).

Variant discovery was performed with gatk v.3.8^81^. BAM files were first processed to realign INDELs using the “RealignerTargetCreator” and “IndelRealigner” tools. An initial set of variants was called for each individual using the “HaplotypeCaller” tool, setting the -stand_call_conf parameter to 30. VCF files from the initial variant calling were then hard filtered to identify low confidence calls by running the “VariantFiltration” tool with the -filterExpression parameter set as follows: “QD < 5.0 || FS > 20.0 || MQ < 30.0 || MQRankSum < −8.0 || MQRankSum > 8.0 || ReadPosRankSum < −2.0 || ReadPosRankSum > 2.0” (hard filtering thresholds were selected by first plotting the values for FS, MQ, MQRankSum, QD and ReadPosRankSum from the initial set of variant calls for selected individuals, representing the range of different sequence and library types used, to visualise their distribution and then modifying the default hard filtering thresholds in line with the guidance provided in the gatk document “Understanding and adapting the generic hard-filtering recommendations” (https://software.broadinstitute.org/gatk/documentation/article.php?id=6925).

Variants passing the gatk hard filtering step (excluding those where alleles had not been called; GT field = ./.) were further analysed using SnpEff v.4.3u^82^, in order to predict the impact of any variants within genes identified from the convergence analyses. Custom genome databases were built for each individual using the SnpEff command “build” with option “-gtf22”; a gtf file containing the annotation for all genes, as well as fasta files containing the genome assembly, CDS and protein sequences, were used as input. Annotation of the impact of variants was performed by running SnpEff with genes of interest specified using the -onlyTr parameter and the -ud parameter set to “0” to deactivate annotation of up or downstream variants. For each variant predicted to alter the protein sequence, the position of the change was checked to see whether it occurred at a site at which evidence for convergence had also been detected and, if so, whether it involved a change to or from the state identified as being convergent between resistant taxa.

We also used the SnpEff results to check for evidence of mutations that could indicate the presence of non-functional gene copies in certain taxa. Variants annotated as “stop gained”, “start lost” or “frameshift” in the 10 ingroup taxa included in the convergence analysis were manually examined to confirm that they would result in a disruption to the expected protein product, and that they were not false positives caused by errors in gene model estimation (e.g. misspecification of intron/exon boundaries). We checked further for evidence of truncation of sequences or errors in the GeMoMa gene model estimation that might be caused by loss-of-function mutations outside of the predicted exon boundaries (e.g. such as the loss of a start codon, which could cause GeMoMa to predict an incomplete gene model if an alternative possible start codon was present downstream). Such putative loss-of-function mutations would not be detected as such by SnpEff because they would be interpreted as low impact intergenic or intron variants.

We used WhatsHap v0.15^83^ to perform read-based phasing of alleles for loci with evidence of multiple variants within them. Input files for phasing in each taxon consisted of the fasta formatted genome assembly, VCF file containing variants passing the gatk hard-filtering step and BAM file from Bowtie 2 with duplicates marked and indels realigned (i.e. as input into variant calling with gatk - see above). The WhatsHap “phase” tool was run with the following parameter settings: --max-coverage 20 --indels; only contigs/scaffolds containing the genes of interest were phased (specified using the “-- chromosome” option). For loci with evidence of variants within the CDS, we used the output of WhatsHap to generate fully or partially phased allele sequences. The SnpSift tool from SnpEff v.4.3u^82^ was used to select variants that alter the CDS from the annotated VCF file generated by SnpEff and the positions of these variants checked against the WhatsHap output to see if they fell within phased blocks. For each gene, the number and size (i.e. number of phased variants encompassed) of each phased block was found and a custom Perl script used to select the largest (or joint largest) block for genes with at least one block spanning multiple phased variants within the CDS. Details of phased variants impacting the CDS within the selected blocks, and of variants for genes with only a single variant within the CDS (which were not considered for phasing with WhatsHap, but for which the CDS for separate alleles can be generated), were extracted from the WhatsHap output VCF file; these selected variants were then applied to the gene sequences from each genome assembly to generate individual alleles which are fully or partially phased within the CDS. The “faidx” function in samtools v.1.6^80^ was used to extract the relevant subsequences from the genome assembly files, and the “consensus” command in bcftools v.1.4 (http://www.htslib.org/doc/bcftools-1.4.html) used to obtain the sequence for each allele with the “-H 1” and “-H 2” options. In cases where the selected phased block also spans unphased variants, both sequences output by bcftools will have the state found in the original genome assembly at these sites, as they will for any variants outside of the selected phased block. The revseq and descseq tools from EMBOSS v.6.6.0^71^ were used to reverse complement the sequences for any genes annotated on the minus strand and to rename the output sequences. Phased alleles were used for further phylogenetic analysis of candidate loci (see below); phasing results were also used to check loci with multiple potential loss-of-function mutations within a single individual, to establish whether the mutations are on the same or different alleles. We discounted any cases of potential loss-of-function mutations where multiple frameshifts occurring in close proximity on the same allele resulting in the correct reading frame being maintained.

To check for polymorphism within *F. excelsior* at sites with evidence of convergence, we examined the combined BAM file generated from mapping Illumina HiSeq reads from 37 individuals from different European provenances (the European Diversity Panel) to the *F. excelsior* reference genome (BATG0.5) by Sollars et al^11^. Duplicate reads were removed from the BAM file using the “MarkDuplicates” function in picard tools v.1.139 (http://broadinstitute.github.io/picard), with the REMOVE_DUPLICATES option set to “true”. Selected contigs (containing the genes of interest) were extracted from the BAM file using the “view” function in samtools v.1.6^80^ and visualised with Tablet v.1.17.08.17^84^; evidence for polymorphism was observed directly from the reads and variants only recorded if supported by at least 10% of reads at that site. Loci OG39275 and OG46977 were excluded from this analysis due to errors in the reference gene models, possibly arising from misassembly, which meant the sites homologous to those with evidence of convergence between EAB-resistant taxa could not be identified (see Supplementary Table 5 for more details).

### Distinguishing between different underlying causes of convergent patterns

To test whether evidence of convergence found by grand-conv might actually be due to taxa sharing the same amino acid variant as a result of introgression or incomplete lineage sorting (ILS), we conducted phylogenetic analyses of coding DNA sequences for the candidate loci to infer their gene-trees. If introgression or ILS were the cause of the patterns detected by grand-conv, we would expect sequences from loci with apparently convergent residues to group together within their gene-tree, even when nucleotides encoding those residues are removed. The CDSs for the refined set of candidate loci were aligned with MUSCLE via GUIDANCE2 and alignment files with unreliably aligned codons removed were used for downstream analyses, as described above (see - Species-tree inference); none of datasets had sequences that were identified by GUIDANCE2 as being unreliably aligned. OG40061 failed to align due to the presence of an incomplete codon at the end of the reference gene model from *F. excelsior*; we trimmed the final 2bp from the *F. excelsior* sequence and reran GUIDANCE2 using this modified file.

Phased allele sequences generated using the WhatsHap results (see above - Analysis of variants within candidate loci) were added to the CDS alignments using MAFFT v.7.310^85^ with the options “--add” and “--keeplength”, in order to splice out any introns present in the phased sequences and maintain the original length of the CDS alignments. For any taxa for which phased sequences had been added, the original unphased sequence was removed from the alignment.

If intragenic recombination has taken place, gene-trees inferred from the CDS alignments may fail to group together the sequences with evidence of convergence even in cases where the convergent pattern is due to ILS or introgressive hybridisation. This is because the phylogenetic signal from any non-recombinant fragments of alleles derived from ILS or introgressive hybridisation may not be sufficiently strong to override that from fragments of alleles that have not been subject to these processes. To account for this possibility, we used hyphy v.2.3.14.20181030beta(MPI)^86^ to conduct recombination tests with GARD^87^ with the following parameter settings: 012345 “General Discrete” 3. Where GARD found significant evidence for a recombination breakpoint (*p*-value < 0.05), we partitioned the alignment into non-recombinant fragments for phylogenetic analysis.

Alignment files were converted to nexus format and gene-trees estimated with MrBayes, as described above (see - Species-tree inference). We checked the ASDSF and used Tracer v.1.6.0 (http://beast.bio.ed.ac.uk/Tracer) to inspect the ESS values for each parameter from the post burnin samples and to confirm that the burnin setting (i.e. discarding the first 10% of samples) was sufficient; in cases where runs had not converged after 5M generations (ASDSF ≥0.010), additional generations were run until an ASDSF of <0.010 was reached. We examined the consensus trees generated by MrBayes to look for evidence that sequences sharing the amino acid states inferred as convergent by grand-conv cluster together in the gene-tree, in conflict with relationships inferred in the species-tree for *Fraxinus*. In cases where evidence of such clustering was found, the codon(s) corresponding to the amino acid site(s) at which evidence of convergence was detected were excluded and the MrBayes analysis repeated. In cases where sequences that have the “convergent” amino acid group together in the gene-tree even after the codon(s) for the relevant site(s) have been excluded, we concluded that the evidence of convergence detected by grand-conv is more likely due to introgressive hybridisation or ILS. We also examined the gene-tree topologies to assess whether any of the amino acid states identified as convergent by grand-conv is more likely to be the ancestral state for *Fraxinus*.

### Further characterisation of candidate loci

To identify the gene from *Arabidopsis thaliana* that best matches each of the candidate loci in our refined set, we conducted a TBLASTN search of the *F. excelsior* protein sequence belonging to the relevant OGs against the *A. thaliana* sequences in the nr/nt database in GenBank^88^ and selected the hit with the lowest *E* value. In cases where the OG lacked a sequence from *F. excelsior*, we used the protein sequence from *F. mandshurica* as the query for the TBLASTN search instead.

We also checked for the presence of the *F. excelsior* sequences within the OrthoMCL clusters generated by Sollars et al.^11^ to see if they were associated with the same *A. thaliana* genes as identified by BLAST. We obtained information on the function of the best matching *A. thaliana* genes from The Arabidopsis Information Resource (TAIR; https://www.arabidopsis.org) and the literature. The OrthoMCL analysis conducted by Sollars et al.^11^ also included a range of other plant species, including *S. lycopersicum* (tomato) and the tree species *Populus trichocarpa* (poplar). As tomato is much more closely related to *F. excelsior* than is *A. thaliana*, and poplar is also a tree species, the function of the genes in these taxa may provide a better guide to the function of the *F. excelsior* genes. We therefore also checked the OrthoMCL clusters containing our candidate *F. excelsior* genes to identify putative orthologues, or close paralogues, from tomato and poplar. In cases where the OrthoMCL cluster included multiple tomato or poplar genes, we focused attention on the tomato sequence that also belonged to the OMA group, as the putative orthologue of our *F. excelsior* gene in that species. For poplar, we looked for information on all sequences, unless there were a large number in the cluster (>4). We searched for literature on the function of the tomato and poplar genes, using the gene identifiers from the versions of the genome annotations used for the OrthoMCL analysis^11^ and also looked for information on PhytoMine, in Phytozome 12^66^.

To further clarify the orthology/paralogy relationships between our candidates and genes from other species, we conducted phylogenetic analysis of the relevant OrthoMCL clusters from Sollars et al^11^ for selected loci. Protein sequences belonging to each OrthoMCL cluster were aligned and gene-trees inferred using GUIDANCE2 and MrBayes respectively, as described above for the OGs and HOGs. For the OrthoMCL cluster relating to OG15551, following an initial MrBayes analysis, we removed two incomplete sequences (Migut.O00792.1.p and GSVIVT01025800001, which were missing >25% of characters in the alignment) and two divergent sequences from *A. thaliana* (AT1G74540 and AT1G74550) which are known to derive from a Brassicales-specific retroposition event and subsequent Brassicaceae-specific tandem duplication^89^; the alignment and phylogenetic analysis was then repeated for the reduced dataset.

For OG15551, we generated a sequence logo for regions of the protein containing sites at which evidence of convergence was detected. We obtained putatively homologous sequences by downloading the fasta file for the OMA group (OMA Browser fingerprint YGPIYSF^90^) containing the *A. thaliana CYP98A3* gene (AT2G40890); the sequences were filtered to include only those from angiosperms, with a maximum of one sequence per genus retained (29 genera in total). To this dataset, we added the OG15551 protein sequences for *F. mandshurica* and *F. pennsylvanica* pe-48 and manually aligned the regions containing the relevant sites (positions 208-218 and 474-482 in the *F. excelsior* FRAEX38873_v2_000261700 reference protein). We used WebLogo v.3.7.3^91^ without compositional adjustment to generate logos for each of these regions.

### GO term enrichment analysis

To test for the possibility of overrepresentation of particular functional categories among the candidate loci in our refined set, compared with the complete set of genes used as input for the convergence analyses, we conducted gene ontology (GO) enrichment tests.

Fisher’s exact tests with the “weight” and “elim” algorithms, which take into account the GO graph topology^92^, were run using the topGO package^93^ (v.2.32.0) in R v.3.5.1^94^. We created a “genes-to-GOs” file for the complete set of *F. excelsior* gene models included in the grand-conv analyses, using GO terms from the existing functional annotation for the reference genome (Sollars et al.^11^); only the single longest transcript per gene (see http://www.ashgenome.org/transcriptomes) was included and for any OMA groups that lacked an *F. excelsior* sequence we used the reference model referred to by the majority of other *Fraxinus* sequences in the group (i.e. as indicated in the GeMoMa gene model names). We also created a list of *F. excelsior* reference model genes belonging to our refined set of candidate loci; again, for any OMA groups that lacked an *F. excelsior* sequence, we used the reference model referred to by the majority of other *Fraxinus* sequences in the group. The complete list of *F. excelsior* reference genes included in the grand-conv analyses, and their associated GO terms, was used as the background against which the list of gene models from the refined set of candidate loci was tested. Fisher’s exact test was run separately, with each of the algorithms, to check for enrichment of terms within the biological process (BP), molecular function (MF) and cellular component (CC) domains.

### Protein modelling

The SignalP 5.0 server^95^ and Phobius server^96^ (http://phobius.sbc.su.se/index.html) were used to detect the presence of signal peptides; for SignalP the organism group was set to “Eukarya” and for Phobius the “normal prediction” method was used. All *Fraxinus* sequences belonging to the OMA groups were used as input for the signal peptide analyses; we only concluded that a signal peptide was present if it was predicted by both methods. The NetPhos 3.1 Server (http://www.cbs.dtu.dk/services/NetPhos/) was used with default settings to identify candidate phosphorylation sites for loci where the amino acid variant observed at a site with evidence of convergence included a serine, threonine or tyrosine. The same protein sequences for low and high susceptibility taxa as used for protein modelling (see below) were input to an initial run of NetPhos 3.1; where evidence for phosphorylation site presence/absence was detected with this initial sequence pair (i.e. present in the sequence with the convergent state and absent from that with the non-convergent state, or vice versa) we reran NetPhos 3.1 on all *Fraxinus* sequences from the relevant OMA groups to test if this difference was consistently associated with the convergent/non-convergent state. We only counted as potential phosphorylation sites those for which the NetPhos score for phosphorylation potential was ≥0.900 for all sequences with the putative site.

RaptorX-Binding^97^ (http://raptorx.uchicago.edu/BindingSite/) was used to generate predicted protein models for each of the candidate genes in our refined set, as well as to outline possible binding sites and candidate ligands. Protein sequences for gene models from the *F. excelsior* reference genome were used for initial protein model and binding site prediction, except in cases where *F. excelsior* was not present in the OMA group or where comparison with the other ingroup and outgroup taxa indicated the *F. excelsior* gene model may be incorrect/incomplete; for these loci, the *F. mandshurica* sequences were used instead as, after the reference, the genome assembly for this taxon is one of the highest quality available. For loci for which a binding site could be successfully predicted (i.e. with at least one potential binding site with a pocket multiplicity value of ≥40), additional models were generated for representative resistant (*F. mandshurica* or *F. platypoda*) and susceptible (*F. ornus* or susceptible *F. pennsylvanica*) taxa using Swiss-model^98^ and Phyre2^99^ (intensive mode), with the exact taxon selection depending on which grand-conv pairwise comparison the locus was detected in (see above - Analysis of patterns of sequence variation consistent with molecular convergence) and which taxa had complete gene models. Where errors were detected in the predicted protein sequences for resistant or susceptible taxa (i.e. due to errors in the predicted gene model, which were detected through comparison with sequences from other species, including those from outgroups) these were corrected prior to modelling (e.g. by trimming extra sequence resulting from incorrect prediction of the start codon). Models predicted by the three independent methods (RaptorX-Binding, Swiss-modeller and Phyre2) were compared by aligning them using PyMOL v.2.0 with the align function to check for congruence; only those loci whose models displayed congruence and where the convergent site was located within/close to the putative active site were taken forward for predictive ligand docking analysis (using the Phyre2 and RaptorX-Binding models for the docking itself). In addition, any loci with congruent models where the site with evidence of convergence is also a putative phosphorylation site presence/absence variant, or which are within a putative functional domain, were analysed further. Ligand candidates were selected based on relevant literature and/or the RaptorX-Binding output, with SDF files for each of the molecules being obtained from PubChem (https://pubchem.ncbi.nlm.nih.gov). SDF files were converted to 3d pdb files using Online SMILES Translator and Structure File Generator (https://cactus.nci.nih.gov/translate/), so they could be used with Autodock. Docking analysis was carried out using Autodock Vina v.1.1.2^100^ with the GUI PyRx v.0.8^101^. Following docking, ligand binding site coordinates were exported as SDF files from Pyrex and loaded into PyMOL with the corresponding protein model file for the resistant and susceptible taxa. Binding sites were then annotated and the residues at which evidence for convergence had been detected with grand-conv were labelled.

### Evidence for differential expression of candidate loci in F. pennsylvanica

We used published transcriptome assembly and expression data from *F. pennsylvanica*^25^ to look for evidence of differential expression of our candidate loci in response to EAB larval feeding. This dataset comprised six genotypes of *F. pennsylvanica*, four putatively resistant to EAB and two susceptible to EAB. To identify the orthogues of our genes in the protein sequences of this independently assembled transcriptome^25^, we repeated the OMA clustering analysis (see above - Gene annotation and orthologue inference) with the addition of these data, available as “Fraxinus_pennsylvanica_120313_peptides” at the Harwood Genomics Project website (https://hardwoodgenomics.org). OMA was run as described above, with the SpeciesTree parameter set to ‘estimate’; because we only intended to use the results for the OGs, and not the HOGs, from this analysis, we did not repeat the clustering with a modified species-tree as was done for our main OMA analysis. Having identified the likely orthologous loci from the *F. pennsylvanica* transcriptome^25^, we used the results of the differential expression analysis^25^ to check whether our candidate loci had significantly increased or decreased expression post-EAB feeding in this dataset.

## Data availability

Underlying data for Figure 1 are available in Supplementary Tables 1 and 2. All Illumina sequence data and genome assemblies will be submitted to the European Nucleotide Archive, and the accession number provided, prior to publication. VCF files containing variants called for *Fraxinus* individuals will be submitted to the European Variation Archive, and the accession number provided, prior to publication. All other data are available from the corresponding authors upon reasonable request.

## Code availability

All unpublished code is available upon reasonable request from the corresponding authors.

## Acknowledgements

This research utilised Queen Mary’s Apocrita HPC facility, supported by QMUL Research-IT (http://doi.org/10.5281/zenodo.438045). We thank J. Carlson for providing *F. pennsylvanica* DNA; T. Baxter, S. Brockington, P. Brownless, D. Crowley, S. Honey, R. Irvine, R. Jinks, P. Jones, T. Kirkham, H. McAllister, I. Parkinson and S. Redstone for help with obtaining *Fraxinus* materials from UK collections; T. Poland for providing EAB eggs; M. Miller for propagating trees for the bioassays; R. Matko for preparation of voucher specimens; J. Pellicer for advice on flow cytometry; P. Howard and M. Struebig for advice on preparation of DNA extractions; J. Keilwagen for help with GeMoMa; K. Davies and J. Parker for help with software for convergence analyses; members of the Evolution Labchat group and Rossiter Lab at Queen Mary University of London, for discussions; R. Rose and J. Sayers for advice on protein modelling analyses. This project was funded by the Living with Environmental Change (LWEC) Tree Health and Plant Biosecurity Initiative - Phase 2 (grant number BB/L012162/1) funded jointly by the BBSRC, Defra, Economic and Social Research Council, Forestry Commission, NERC and the Scottish Government. R.J.A.B. acknowledges support from the DEFRA Future Proofing Plant Health scheme. R.J.A.B. and L.J.K. acknowledge support from the Erica Waltraud Albrecht Endowment Fund.

## Author contributions

R.J.A.B. conceived and oversaw the project. L.J.K. and R.J.A.B. wrote the manuscript, with input from J.L.K., W.J.P and S.J.R. L.J.K. conducted gene annotation, orthologue inference, the convergence analyses, calling and analysis of variants, the GO enrichment analysis and all phylogenetic analyses. L.J.K., W.C. and A.T.W. performed genome size estimation by flow cytometry. L.J.K., W.C., E.D.C. and D.W.C. extracted DNA. L.J.K. and E.D.C. assembled the genomes. J.L.K. conceived and oversaw the emerald ash borer (EAB) bioassays. D.W.C. conducted the EAB bioassays. J.L.K. and M.E.M. analysed data from the EAB bioassays. W.J.P. conducted the protein modelling analyses. S.J.R. advised on convergence analyses.

## Correspondence and requests for materials

should be addressed to L.J.K. and R.J.A.B; requests for DNA samples or plant materials may require the signing of a material transfer agreement, and permission from the original source in cases of intended commercial use.

## SUPPLEMENTARY INFORMATION

### Supplementary Tables

Supplementary Table 1. Results from EAB egg bioassays

Supplementary Table 2. Results of statistical analysis of bioassay results

Supplementary Table 3. Details of plant material used for whole genome sequencing and egg bioassays

Supplementary Table 4. Summary of genome size estimates for individuals used for whole genome sequencing, Illumina sequence data generated and genome assembly statistics

Supplementary Table 5. Details for the refined set of 53 candidate loci identified by the grand-conv analyses

Supplementary Table 6. Evidence for potential loss-of-function mutations in the candidate genes

Supplementary Table 7. Significantly enriched GO terms among the candidate genes

Supplementary Table 8. Presence of amino acid states associated with EAB-resistance in the 53 candidate genes across 29 *Fraxinus* genomes

Supplementary Table 9. Positions of candidate amino acid sites in *Fraxinus excelsior* diversity panel and evidence for polymorphism

Supplementary Table 10. Putative orthologues identified from the *Fraxinus pennsylvanica* transcriptome and evidence for differential expression of transcripts

### Supplementary Figures

**Supplementary Figure 1.**
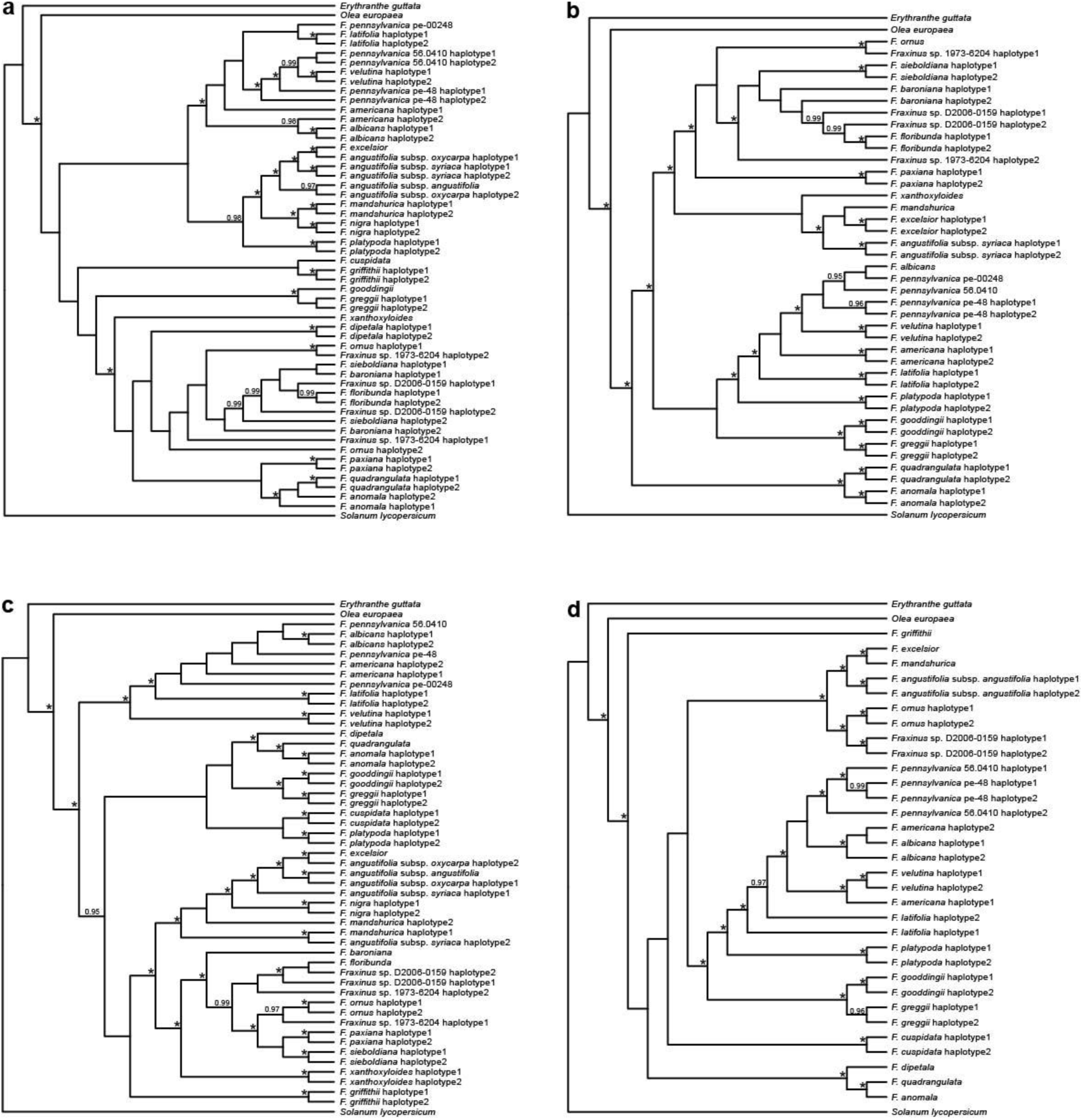
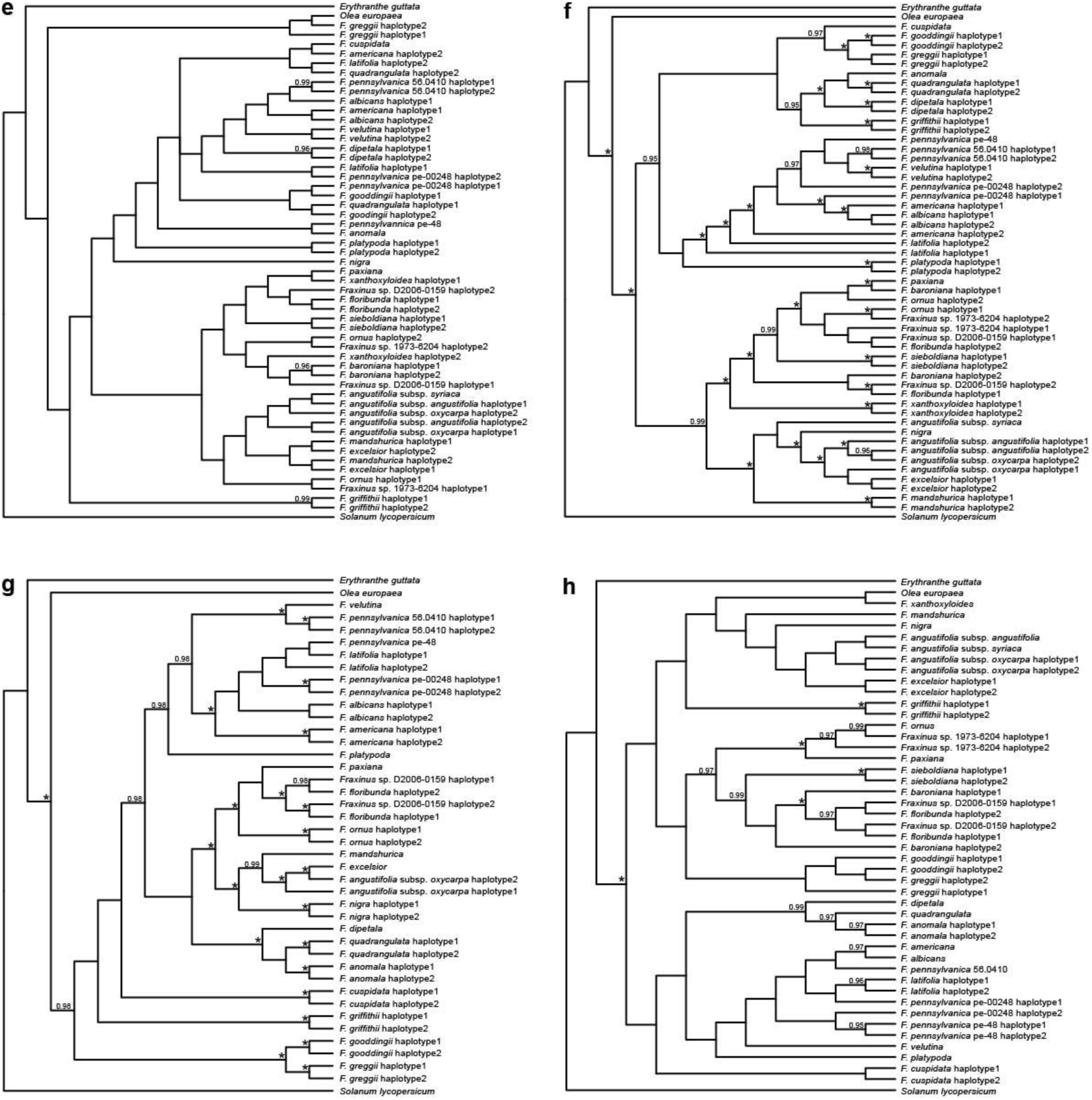
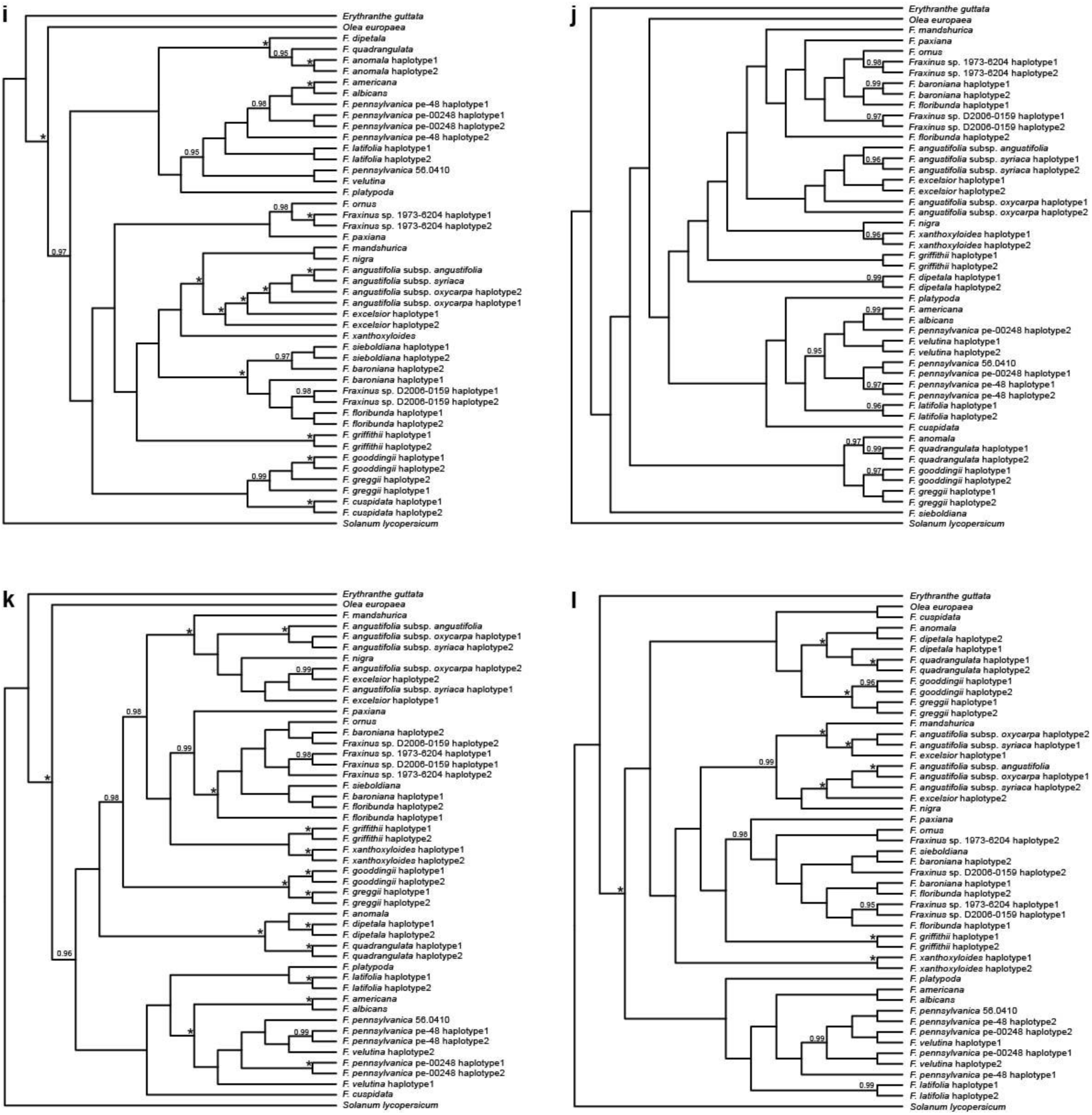
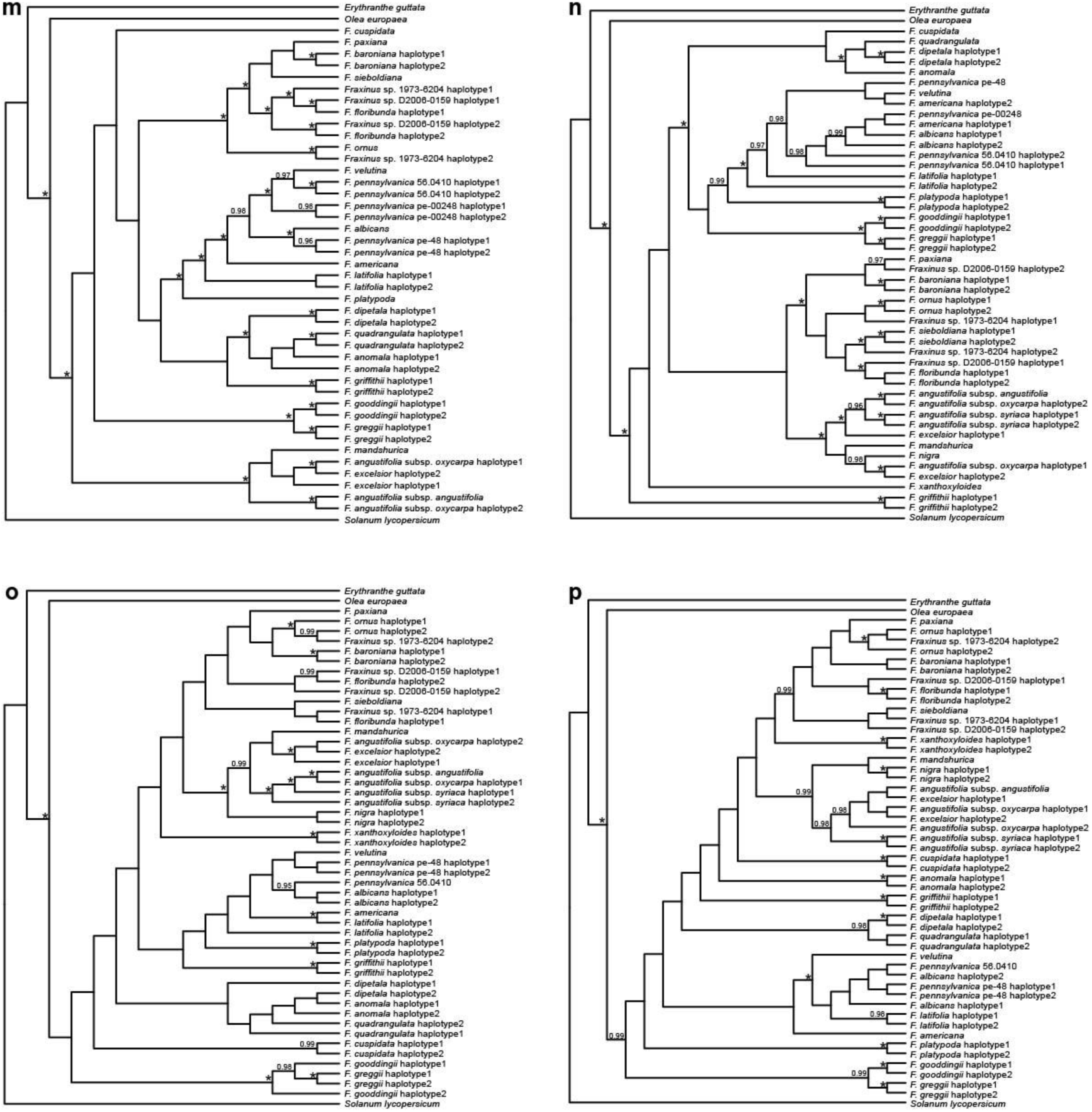
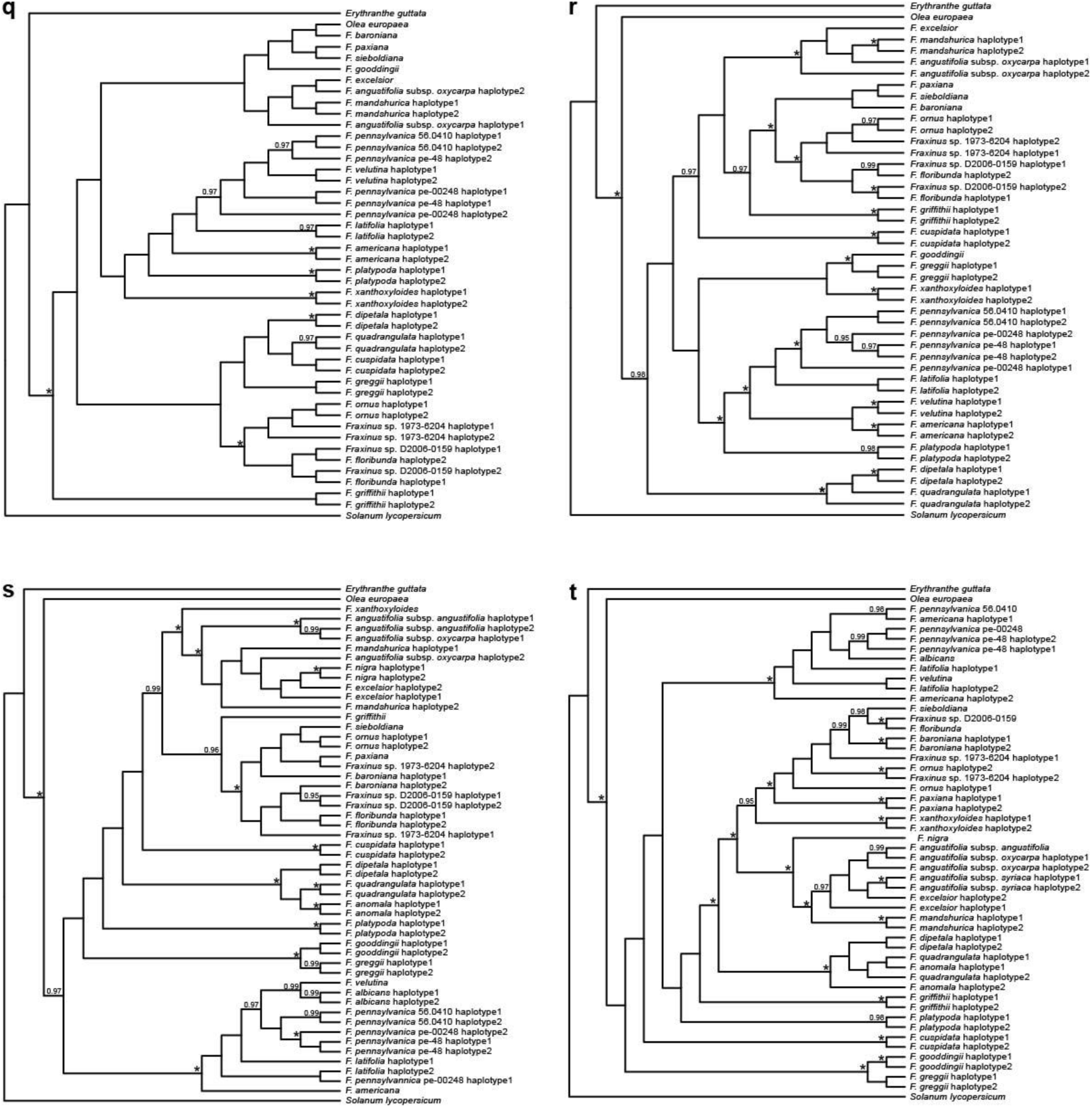
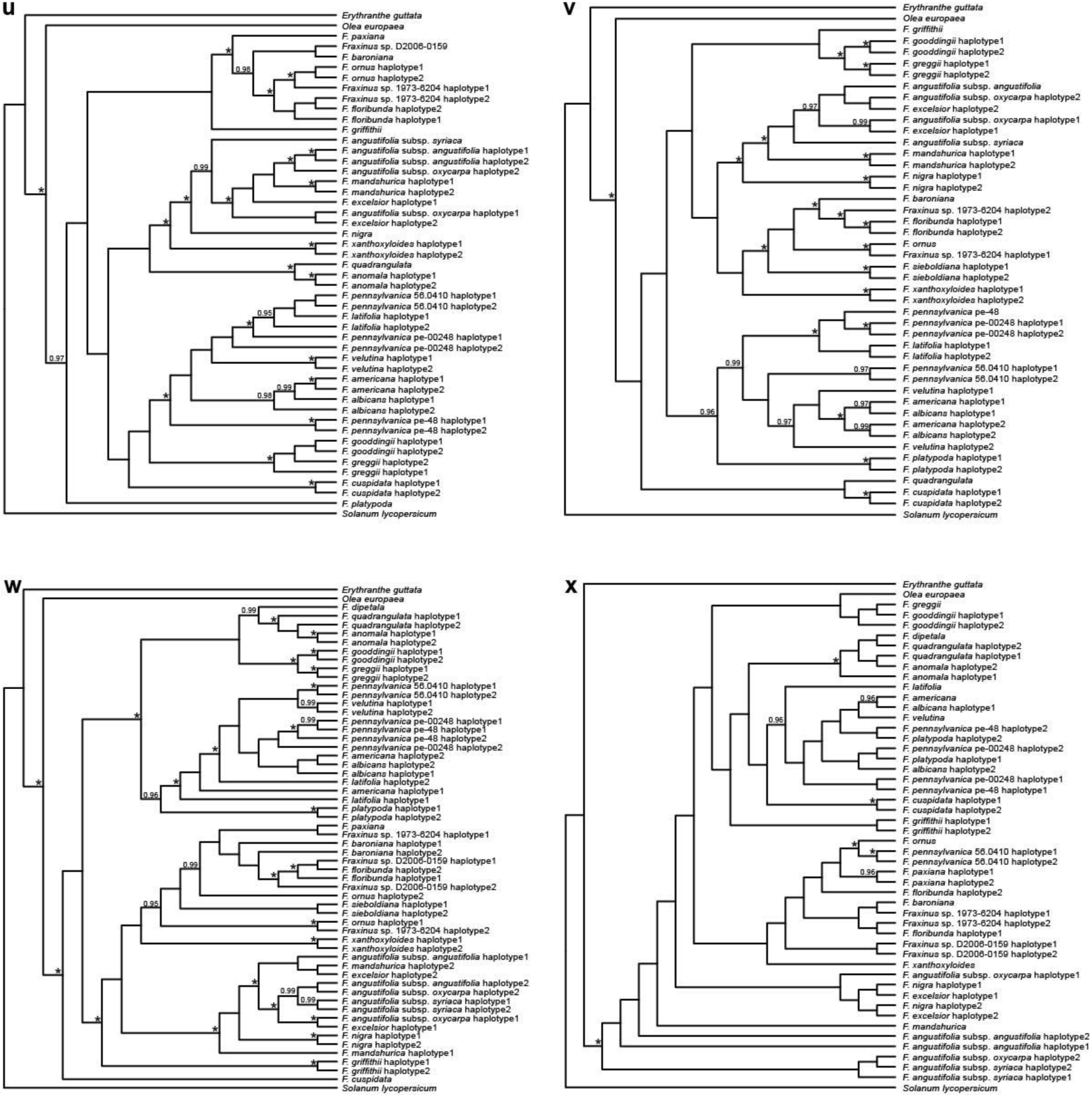
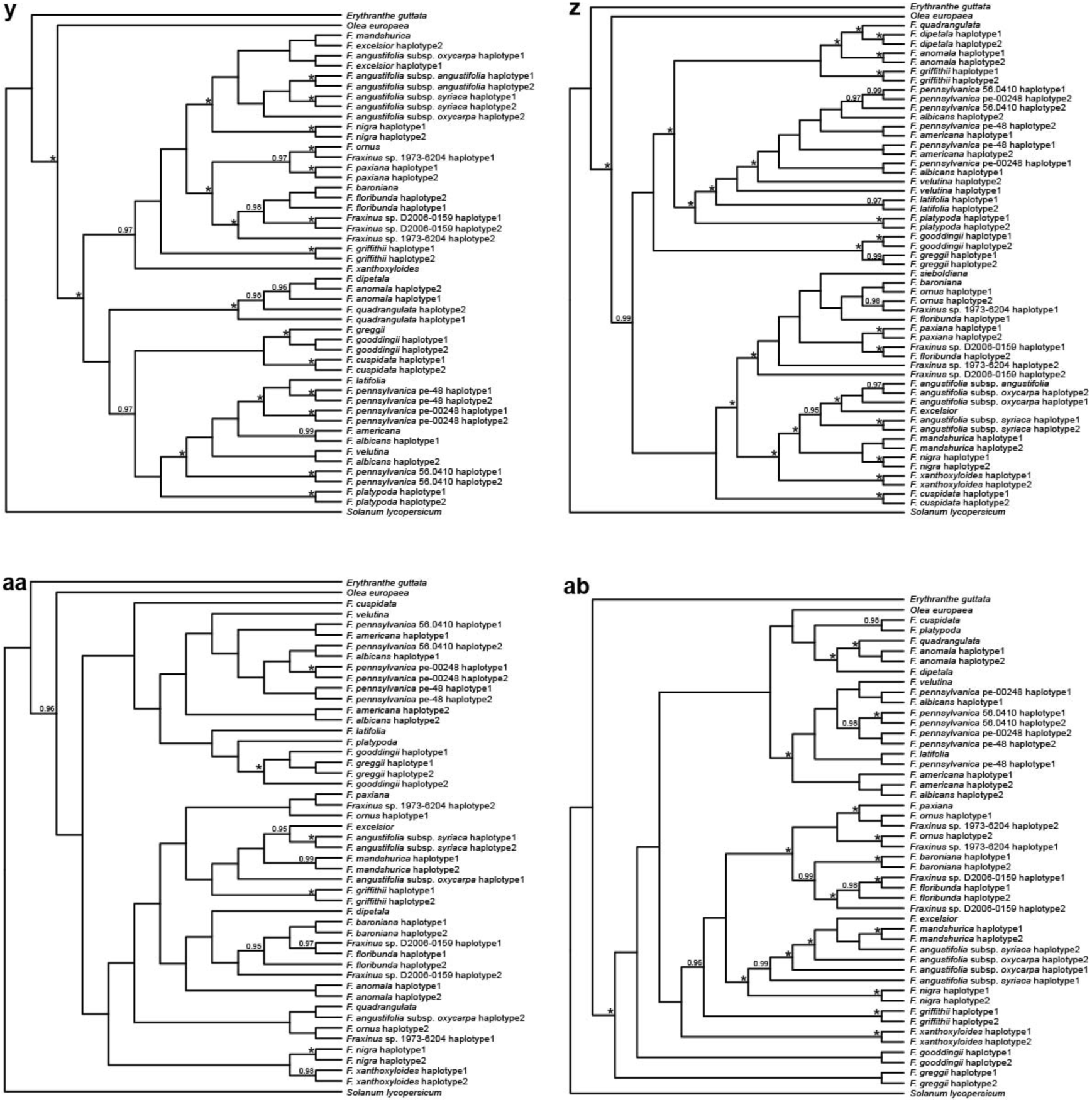
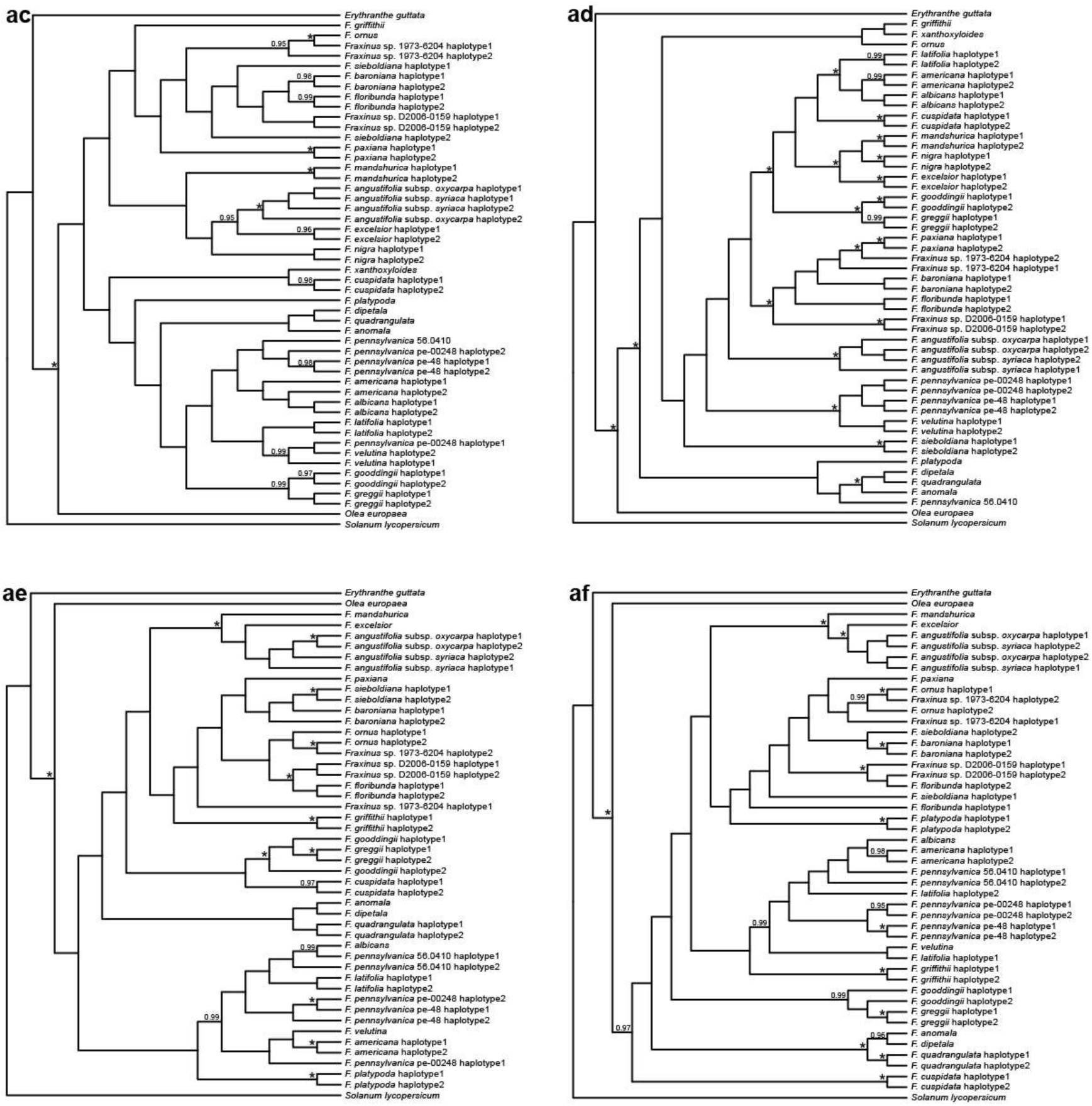
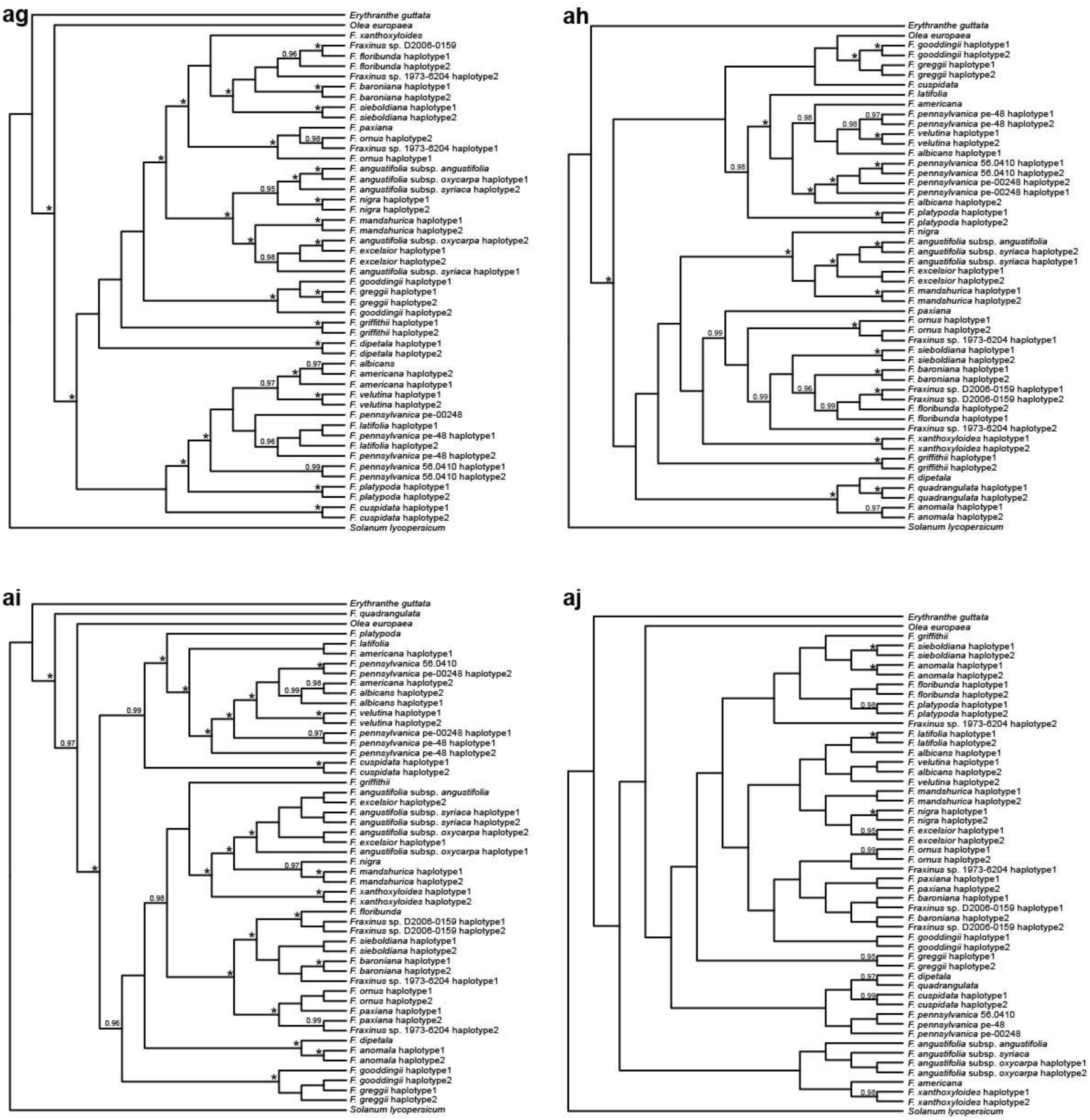
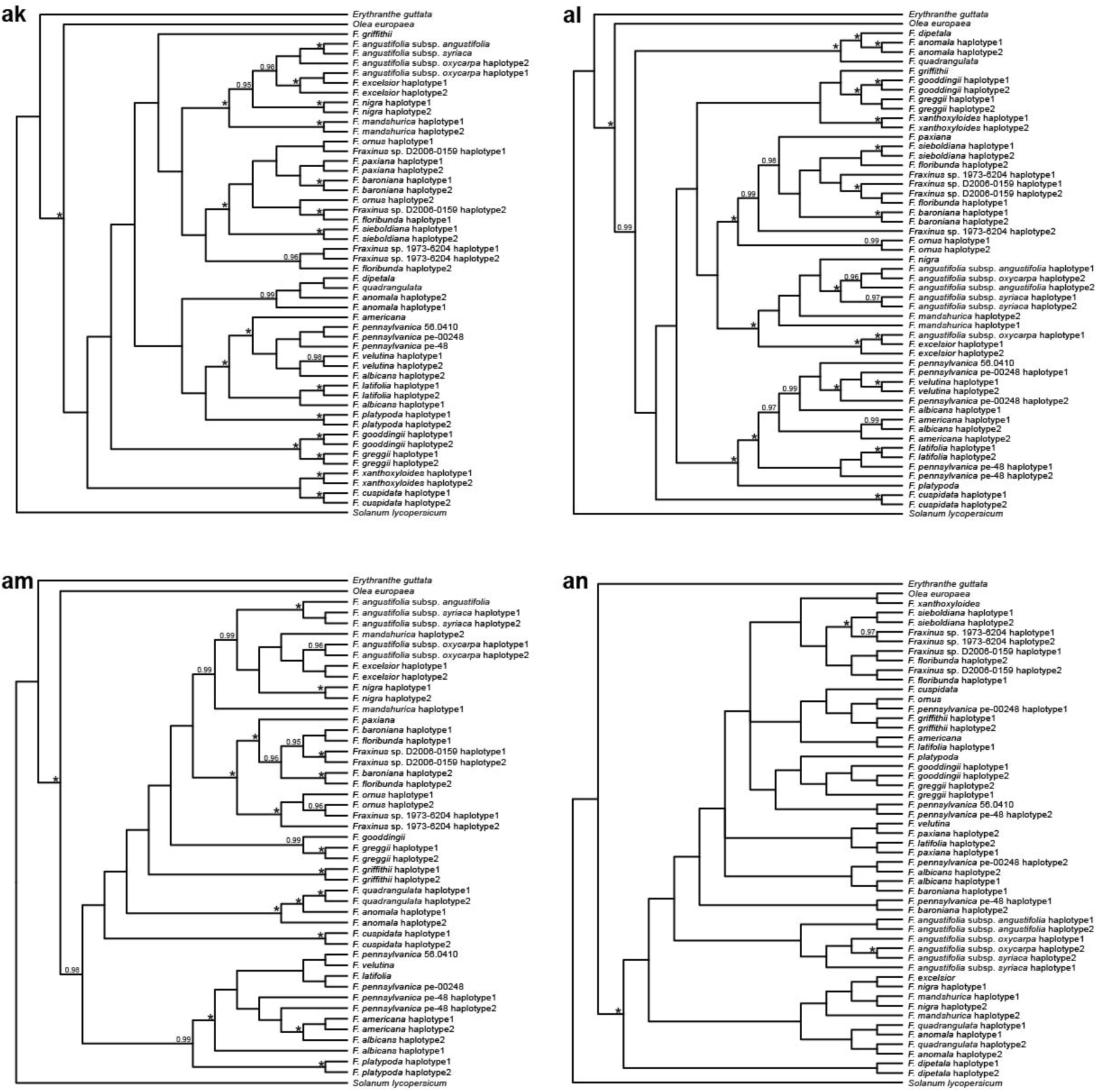
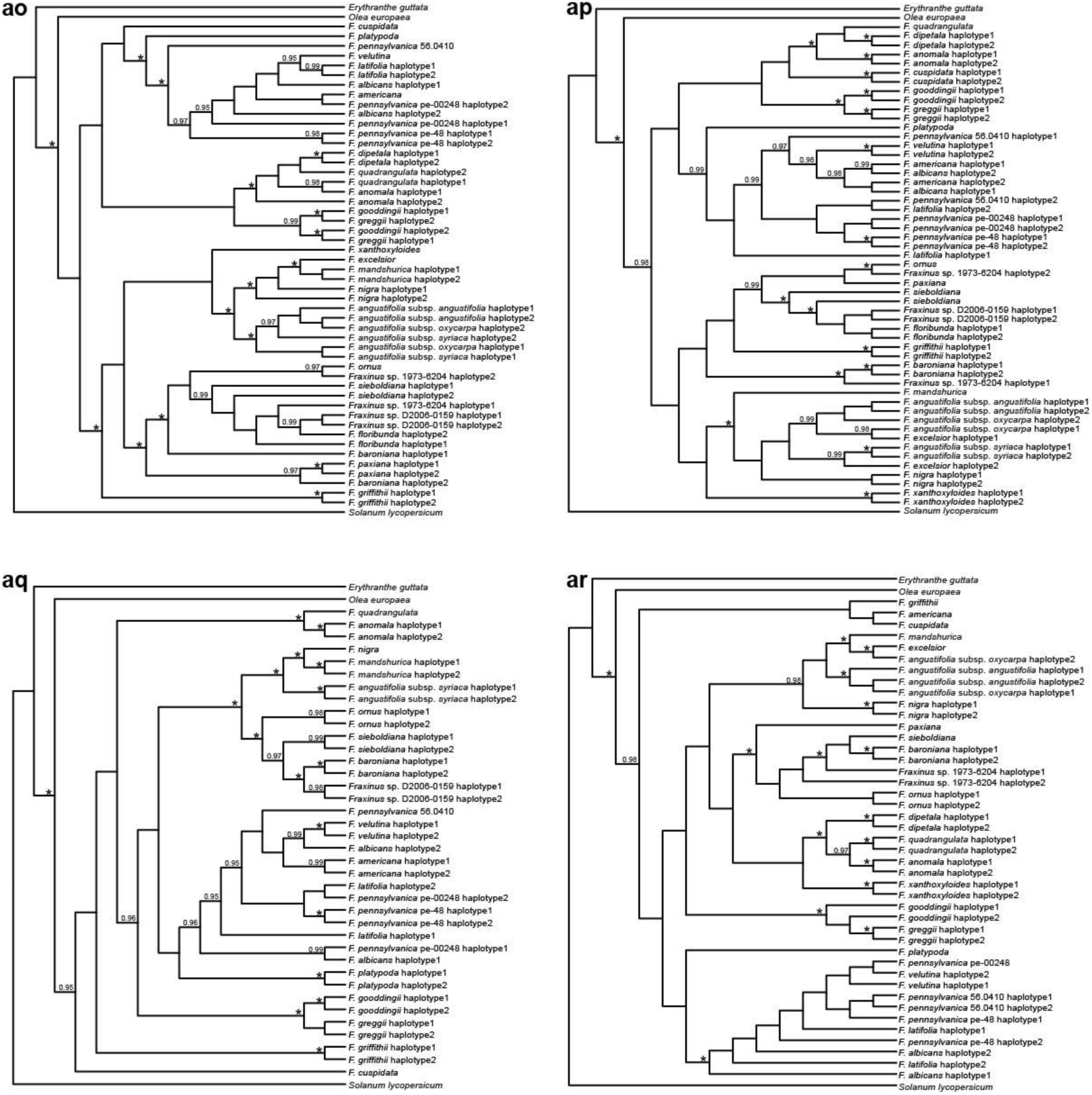
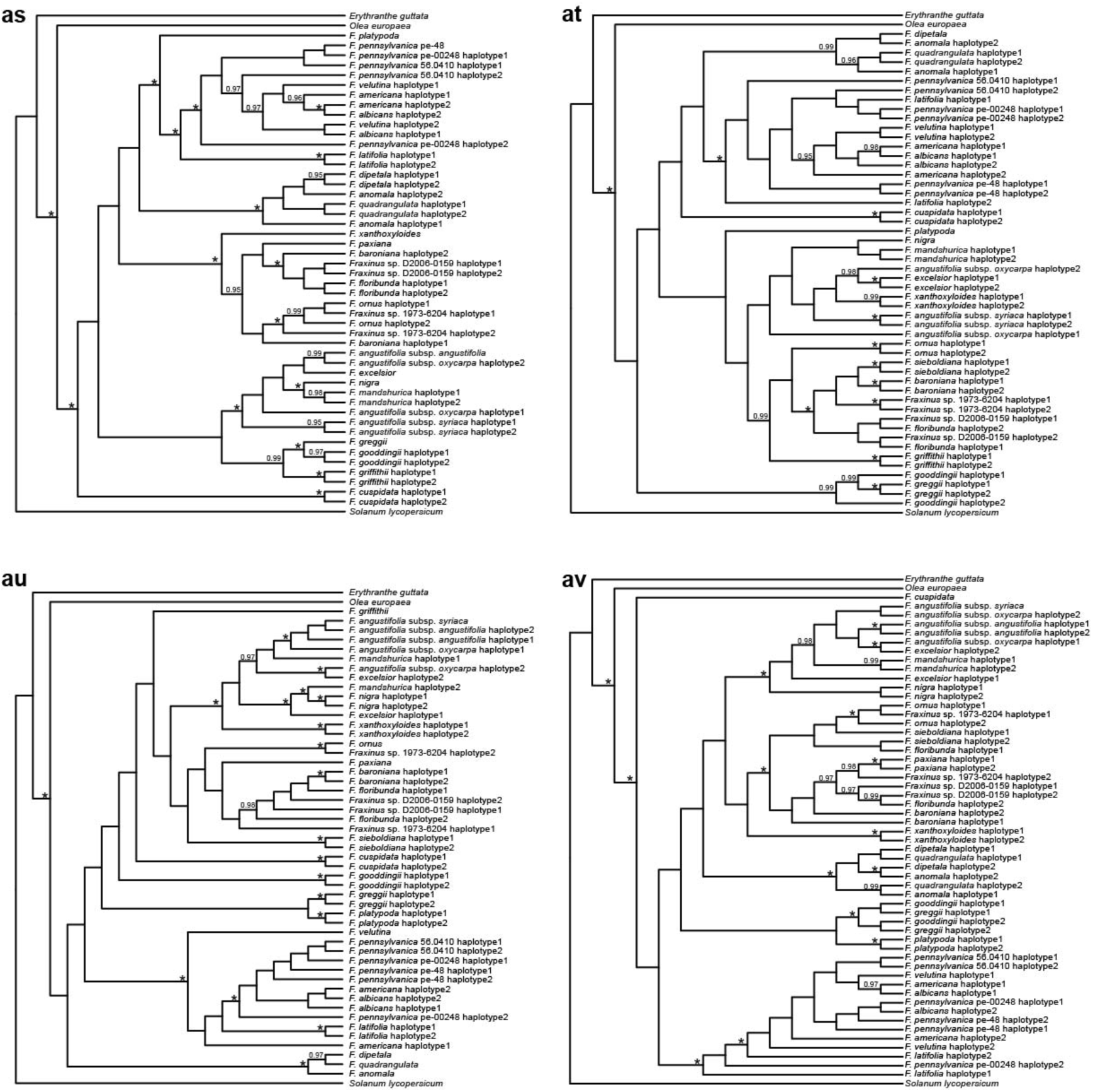
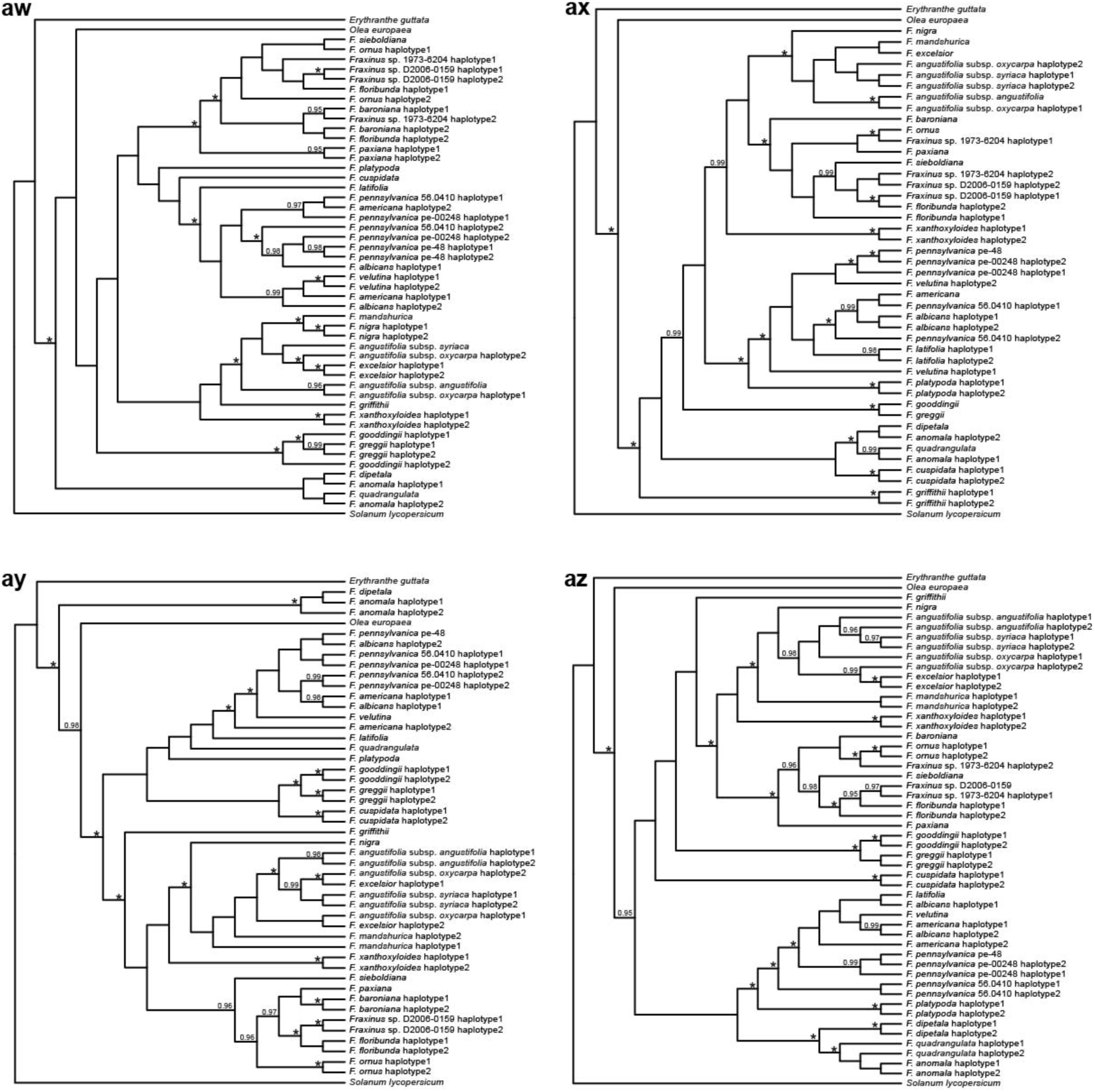
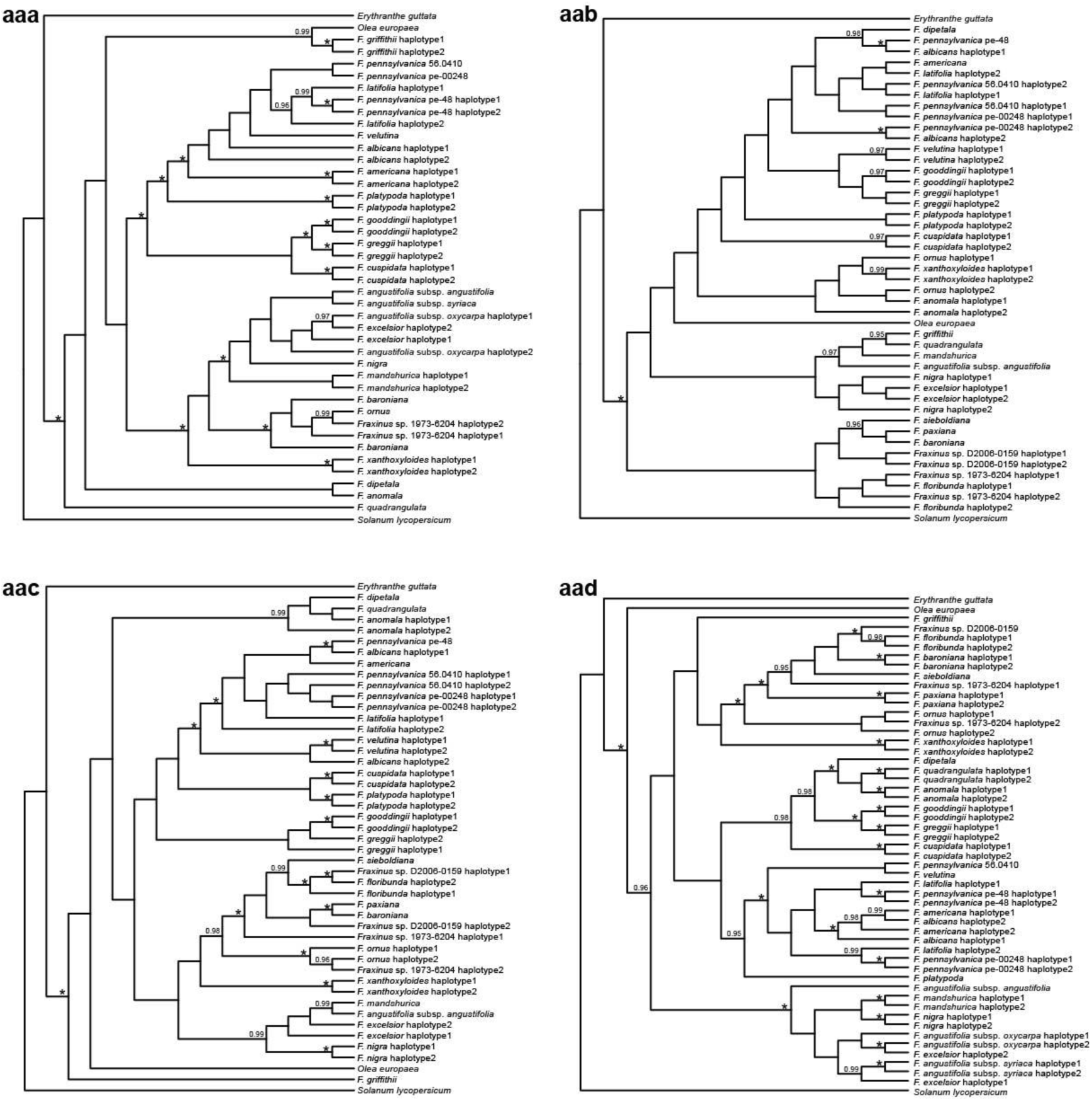
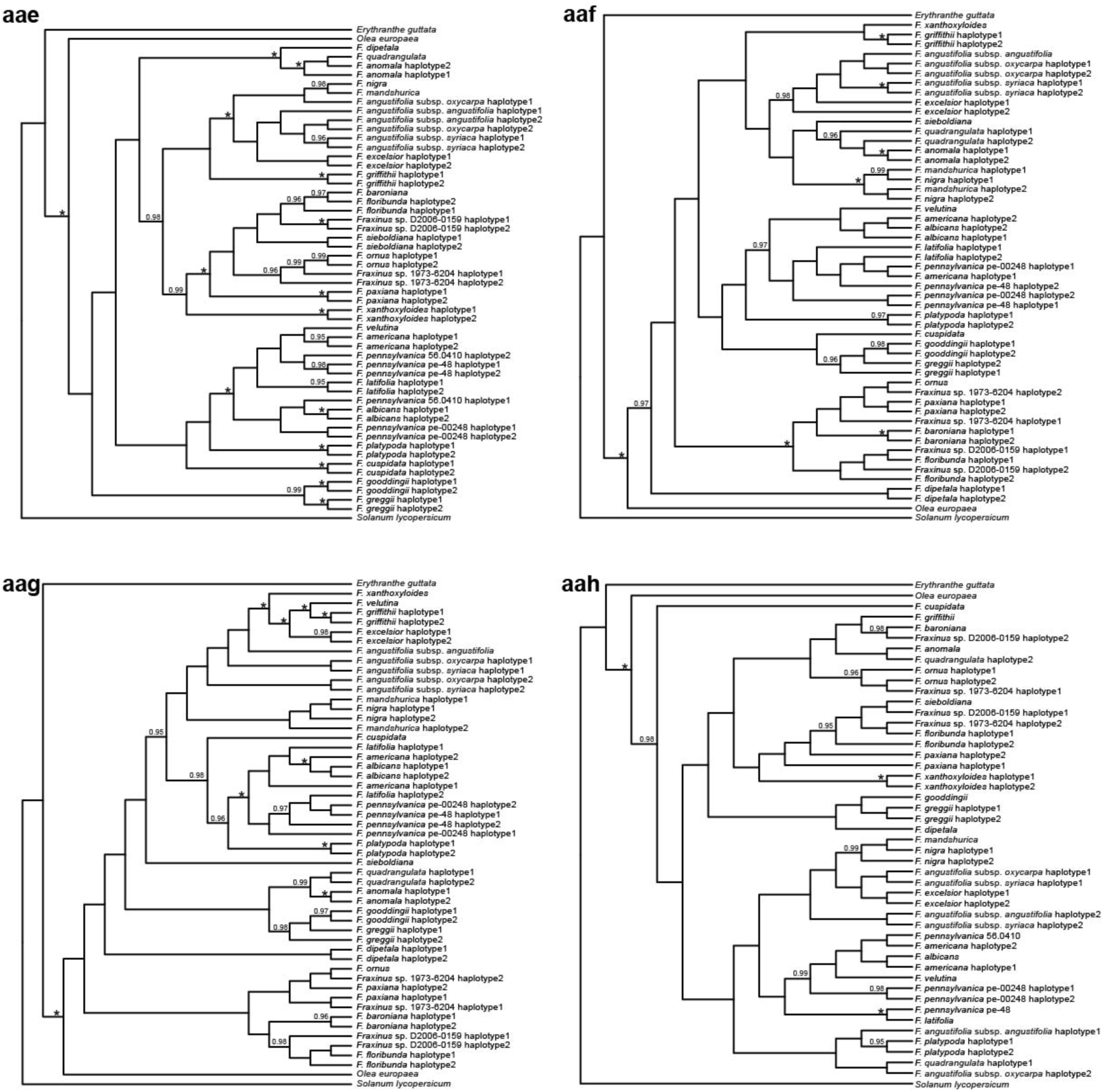
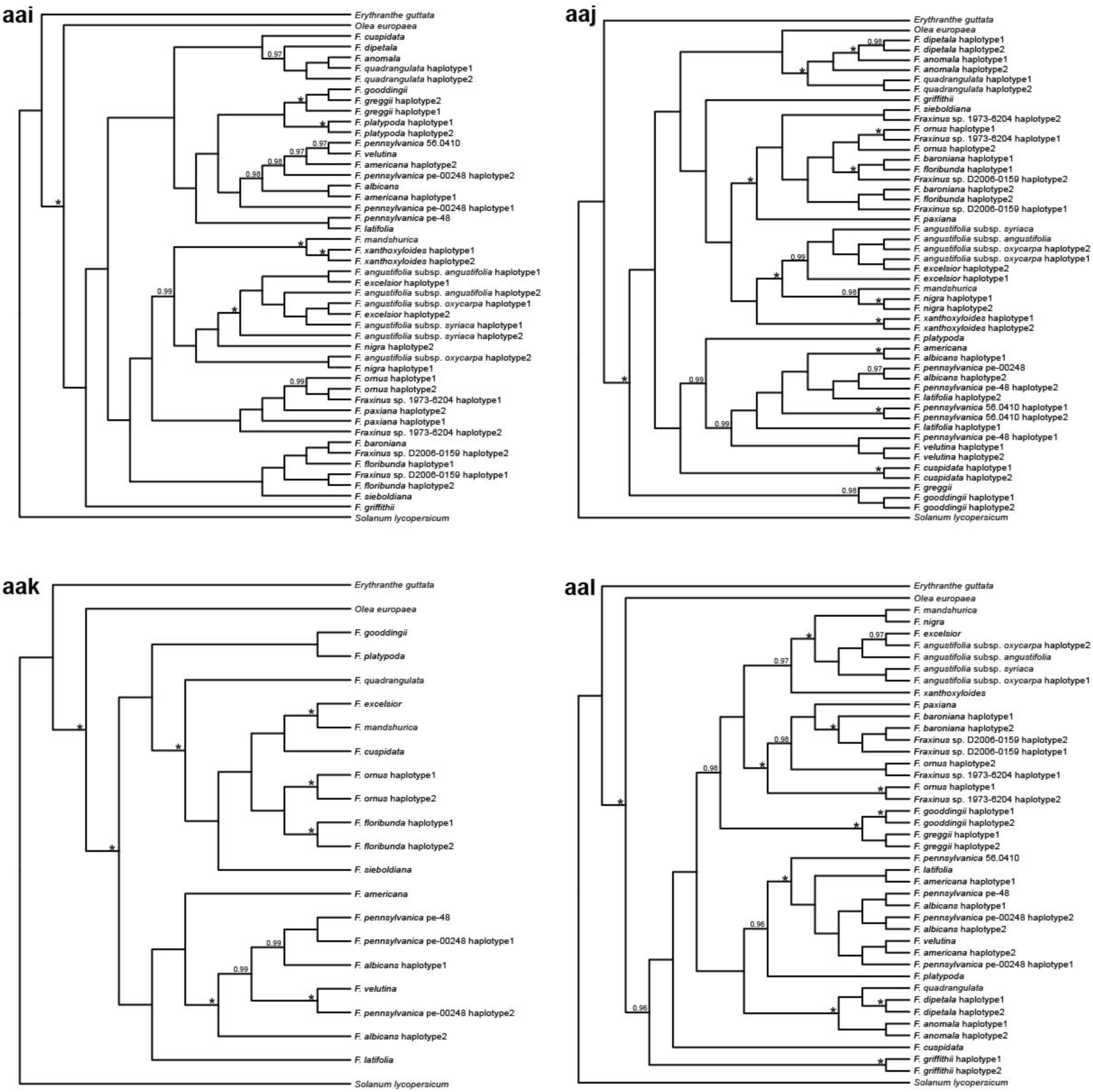
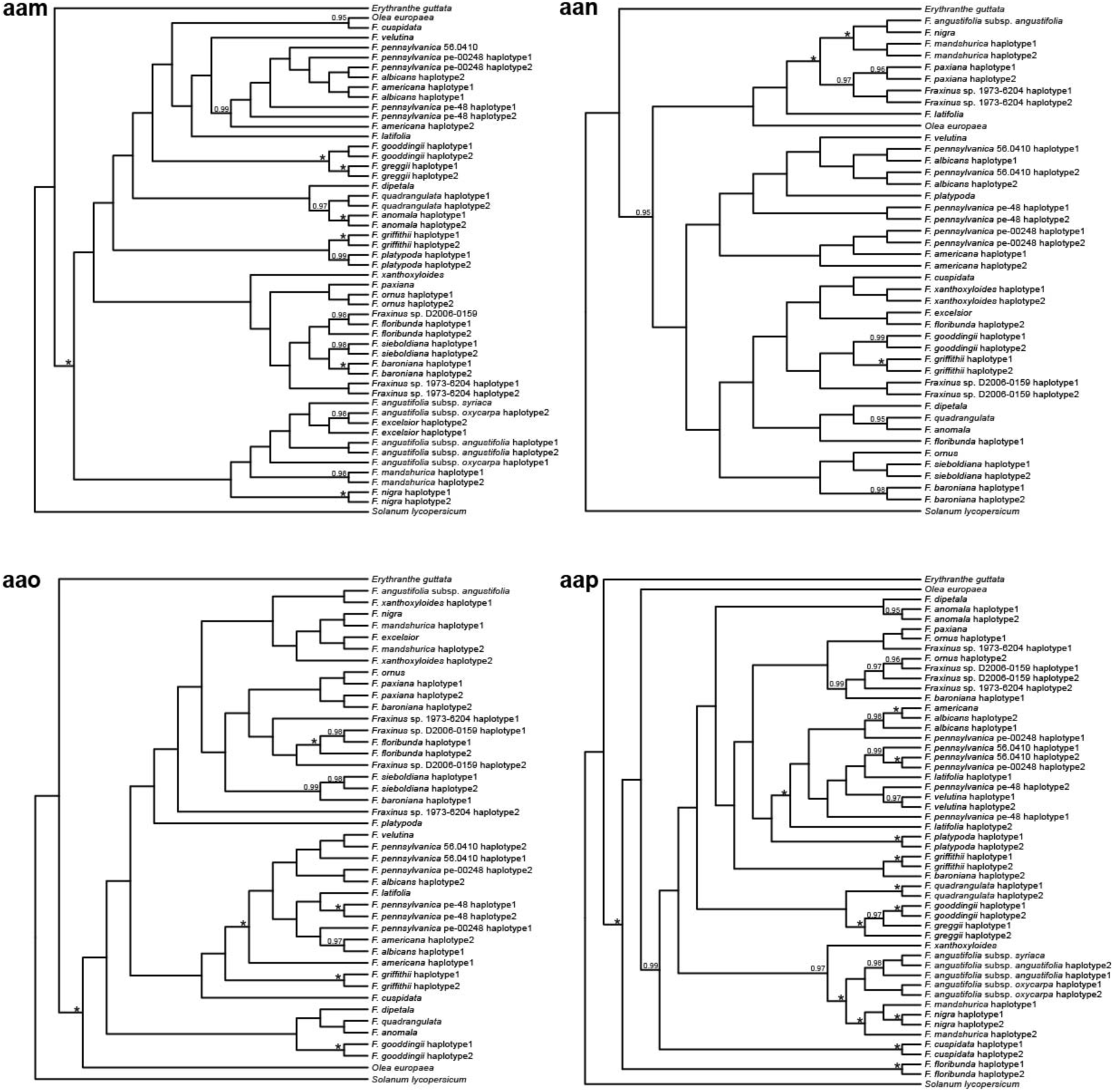
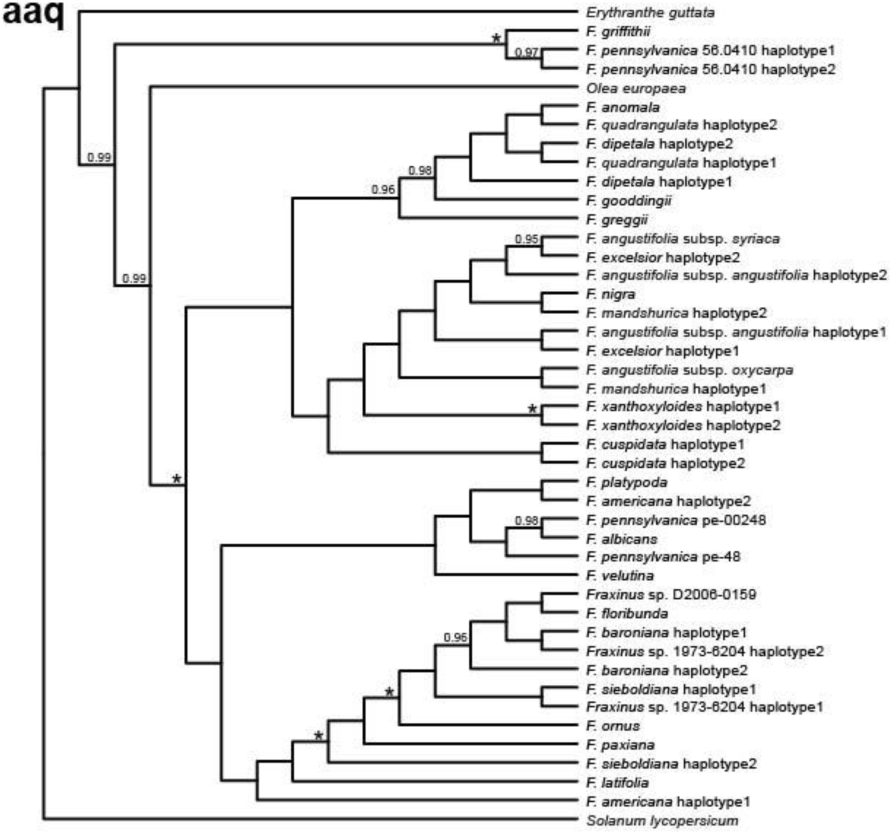
Gene-trees for the 53 candidate loci inferred from CDS alignments including phased alleles and accounting for evidence of recombination. **a**, Gene-tree for OG20252 inferred from positions 378-1524 of the CDS alignment excluding the codon for the amino acid site with evidence of convergence (the gene-tree inferred from positions 1-377 is not shown because the majority of nodes are poorly supported and the codon for the convergent site is within the other fragment). **b**, Gene-tree for OG853. **c**, Gene-tree for OG2897. **d**, Gene-tree for OG4372. **e**, Gene-tree for OG4469 inferred from positions 1-250 of the CDS alignment. **f**, Gene-tree for OG4469 inferred from positions 251-2595 of the CDS alignment. **g**, Gene-tree for OG5539. **h**, Gene-tree for OG6935 inferred from positions 1-794 of the CDS alignment. **i**, Gene-tree for OG6935 inferred from positions 795-2208 of the CDS alignment. **j**, Gene-tree for OG7454 inferred from positions 1-557 of the CDS alignment. **k**, Gene-tree for OG7454 inferred from positions 558-1405 of the CDS alignment. **l**, Gene-tree for OG7454 inferred from positions 1406-2352 of the CDS alignment excluding the codons for the amino acid sites with evidence of convergence. **m**, Gene-tree for OG10762 inferred from the CDS alignment with the codon for the amino acid site with evidence of convergence excluded. **n**, Gene-tree for OG11013. **o**, Gene-tree for OG11720 inferred from positions 1-868 of the CDS alignment excluding the codon for the amino acid site with evidence of convergence. **p**, Gene-tree for OG11720 inferred from positions 869-1773 of the CDS alignment. **q**, Gene-tree for OG13887 inferred from positions 1-269 of the CDS alignment. **r**, Gene-tree for OG13887 inferred from positions 270-1803 of the CDS alignment. **s**, Gene-tree for OG15551 inferred from the CDS alignment with the codons for the amino acid sites with evidence of convergence excluded. **t**, Gene-tree for OG16673 inferred from the CDS alignment with the codon for the amino acid site with evidence of convergence excluded. **u**, Gene-tree for OG16739. **v**, Gene-tree for OG17252. **w**, Gene-tree for OG19104. **x**, Gene-tree for OG20520 inferred from positions 1-372 of the CDS alignment. **y**, Gene-tree for OG20520 inferred from positions 373-1452 of the CDS alignment. **z**, Gene-tree for OG20859. **aa**, Gene-tree for OG21033 inferred from positions 1-327 of the CDS alignment. **ab**, Gene-tree for OG21033 inferred from positions 328-1539 of the CDS alignment. **ac**, Gene-tree for OG21449 inferred from positions 1-455 of the CDS alignment. **ad**, Gene-tree for OG21449 inferred from positions 456-1209 of the CDS alignment. **ae**, Gene-tree for OG23214 inferred from positions 1-836 of the CDS alignment. **af**, Gene-tree for OG23214 inferred from positions 837-1401 of the CDS alignment. **ag**, Gene-tree for OG23284. **ah**, Gene-tree for OG24614. **ai**, Gene-tree for OG24969. **aj**, Gene-tree for OG26964 inferred from positions 1-205 of the CDS alignment. **ak**, Gene-tree for OG26964 inferred from positions 206-1092 of the CDS alignment. **al**, Gene-tree for OG27080. **am**, Gene-tree for OG27693. **an**, Gene-tree for OG27838 inferred from positions 1-381 of the CDS alignment. **ao**, Gene-tree for OG27838 inferred from positions 382-1707 of the CDS alignment. **ap**, Gene-tree for OG28712. **aq**, Gene-tree for OG30208. **ar**, Gene-tree for OG32176 inferred from the CDS alignment with the codons for the amino acid sites with evidence of convergence excluded. **as**, Gene-tree for OG33348. **at**, Gene-tree for OG35707. **au**, Gene-tree for OG36502. **av**, Gene-tree for OG37560. **aw**, Gene-tree for OG37870. **ax**, Gene-tree for OG38407. **ay**, Gene-tree for OG38543. **Az**, Gene-tree for OG39275. **aaa**, Gene-tree for OG40061. **aab**, Gene-tree for OG41448 inferred from positions 1-243 of the CDS alignment. **aac**, Gene-tree for OG41448 inferred from positions 244-981 of the CDS alignment excluding the codon for the amino acid site with evidence of convergence. **aad**, Gene-tree for OG41488. **aae**, Gene-tree for OG43828. **aaf**, Gene-tree for OG46977 inferred from positions 1-291 of the CDS alignment. **aag**, Gene-tree for OG46977 inferred from positions 292-804 of the CDS alignment. **aah**, Gene-tree for OG47560 inferred from positions 1-280 of the CDS alignment. **aai**, Gene-tree for OG47560 inferred from positions 281-852 of the CDS alignment excluding the codons for the amino acid sites with evidence of convergence. **aaj**, Gene-tree for OG47629. **aak**, Gene-tree for OG49074. **aal**, Gene-tree for OG50989. **aam**, Gene-tree for OG56563. **aan**, Gene-tree for OG59564 inferred from positions 1-189 of the CDS alignment. **aao**, Gene-tree for OG59564 inferred from positions 190-558 of the CDS alignment excluding the codon for the amino acid site with evidence of convergence. **aap**, Gene-tree for OG60899. **aaq**, Gene-tree for OG64545. Gene-trees were inferred with MrBayes and rooted on *Solanum lycopersicum*. Numbers above branches are posterior probabilities (PP) of ≥0.95; asterisks indicate nodes with PP=1.

**Supplementary Figure 2.**
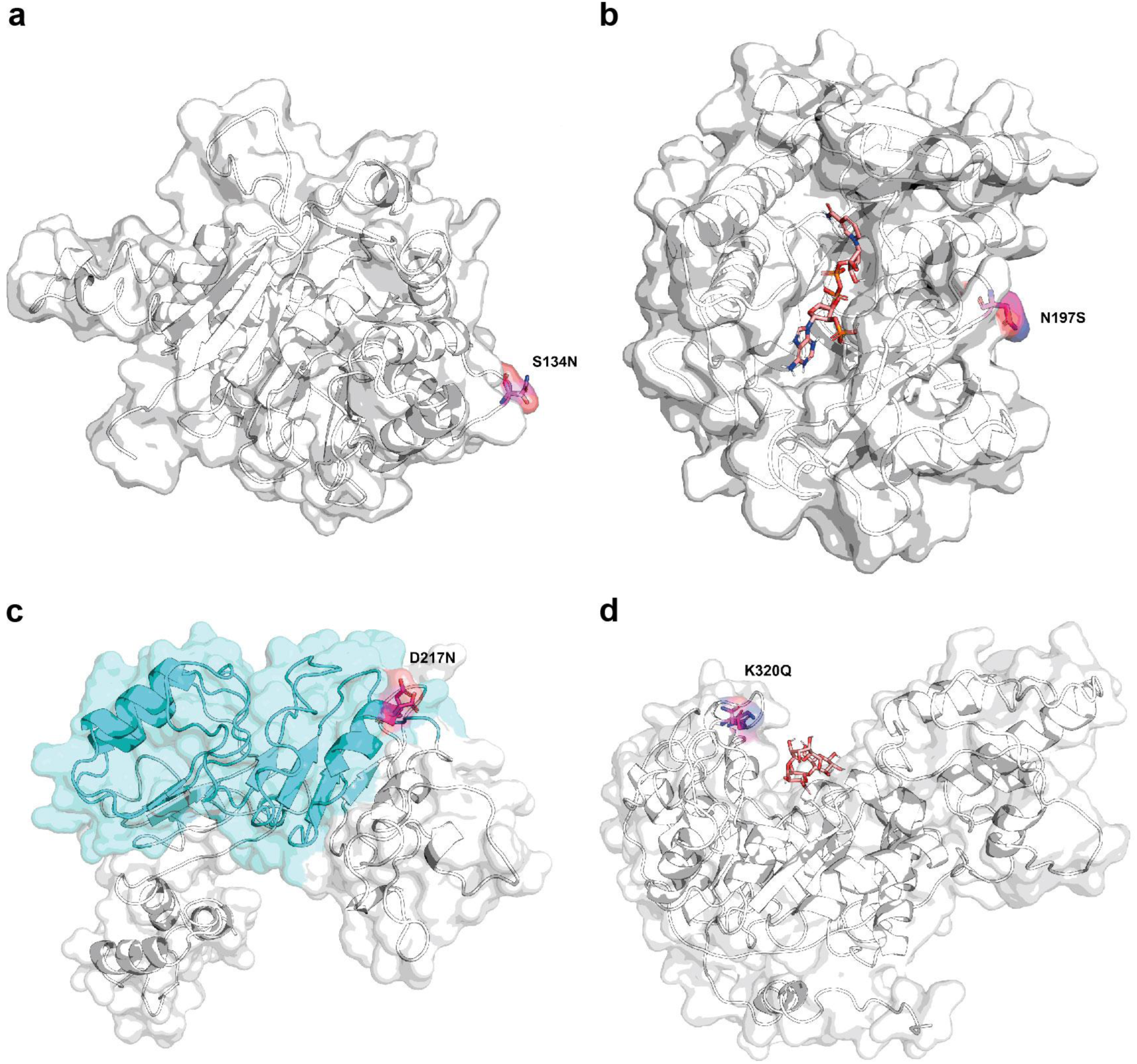
Predicted protein structures for selected candidate loci. **a**, Predicted protein structure for OG36502, modelled using the protein sequence for *Fraxinus platypoda*. The serine/asparagine variant at the site where convergence was detected is highlighted; the serine is a putative phosphorylation site. **b**, Predicted protein structure for OG40061, modelled using the protein sequence for *F. mandshurica*. The asparagine/serine variant at the site where convergence was detected is highlighted; the serine is a putative phosphorylation site. The putative substrate, NADP, is shown docked within the predicted active site. **c**, Predicted protein structure for OG38407, modelled using the protein sequence for *F. mandshurica*. The aspartic acid/asparagine variant at the site where convergence was detected is highlighted; the site falls within a leucine rich repeat region (LRR; shaded blue) which is predicted to span from position 111-237 within the protein sequence (detected using the GenomeNet MOTIF tool (www.genome.jp/tools/motif/), searching against the NCBI-CDD and Pfam databases with default parameters; the LRR region was identified as positions 111-237 with an e-value of 1e-05). **d**, Predicted protein structure for OG21033, modelled using the protein sequence for *F. platypoda*. The lysine/glutamine at the site where convergence was detected is highlighted. The putative substrate, β-D-Glcp-(1→3)-β-D-GlcpA-(1→4)-β-D-Glcp, is shown docked within the predicted active site.

**Supplementary Figure 3.**
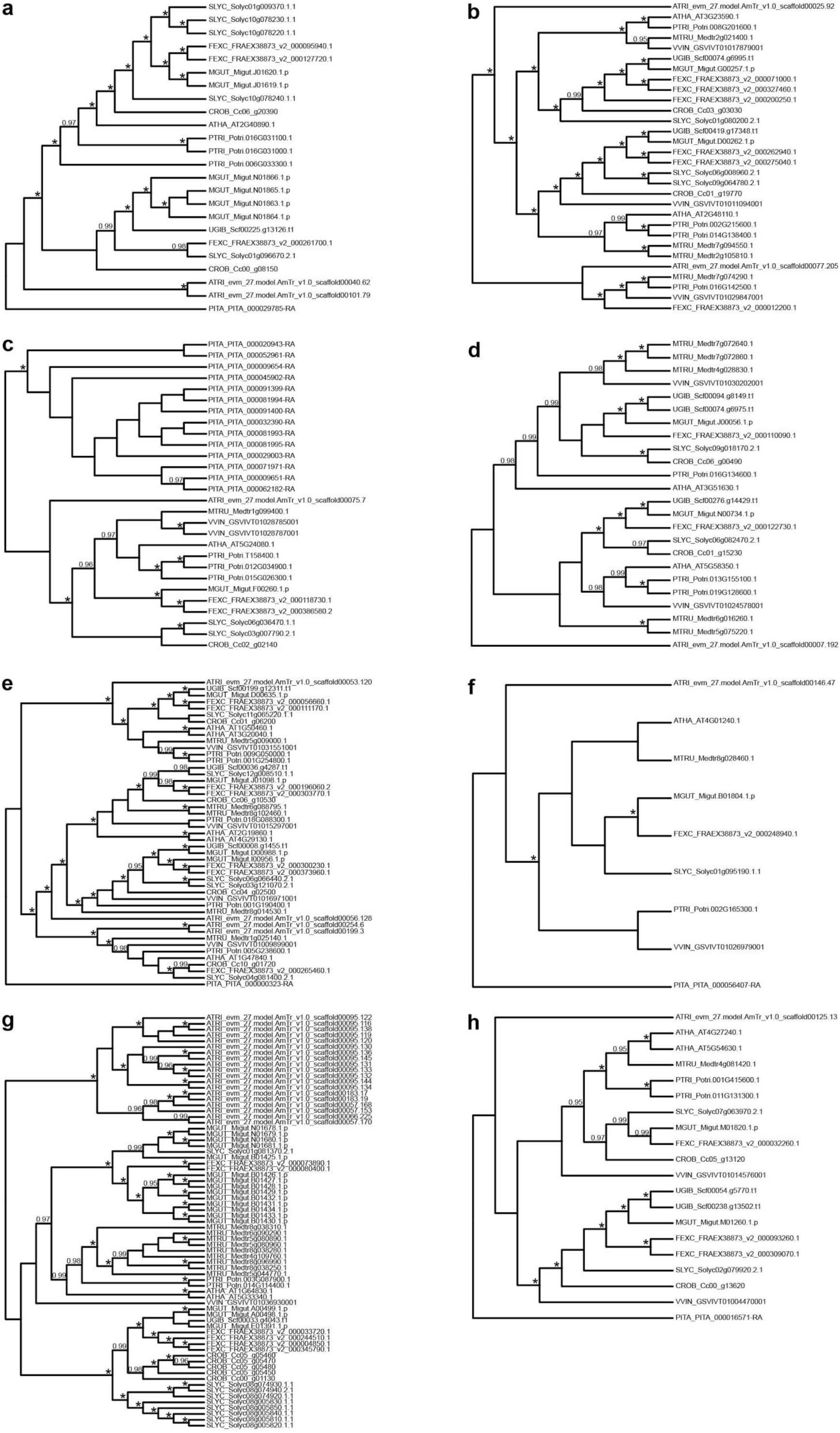
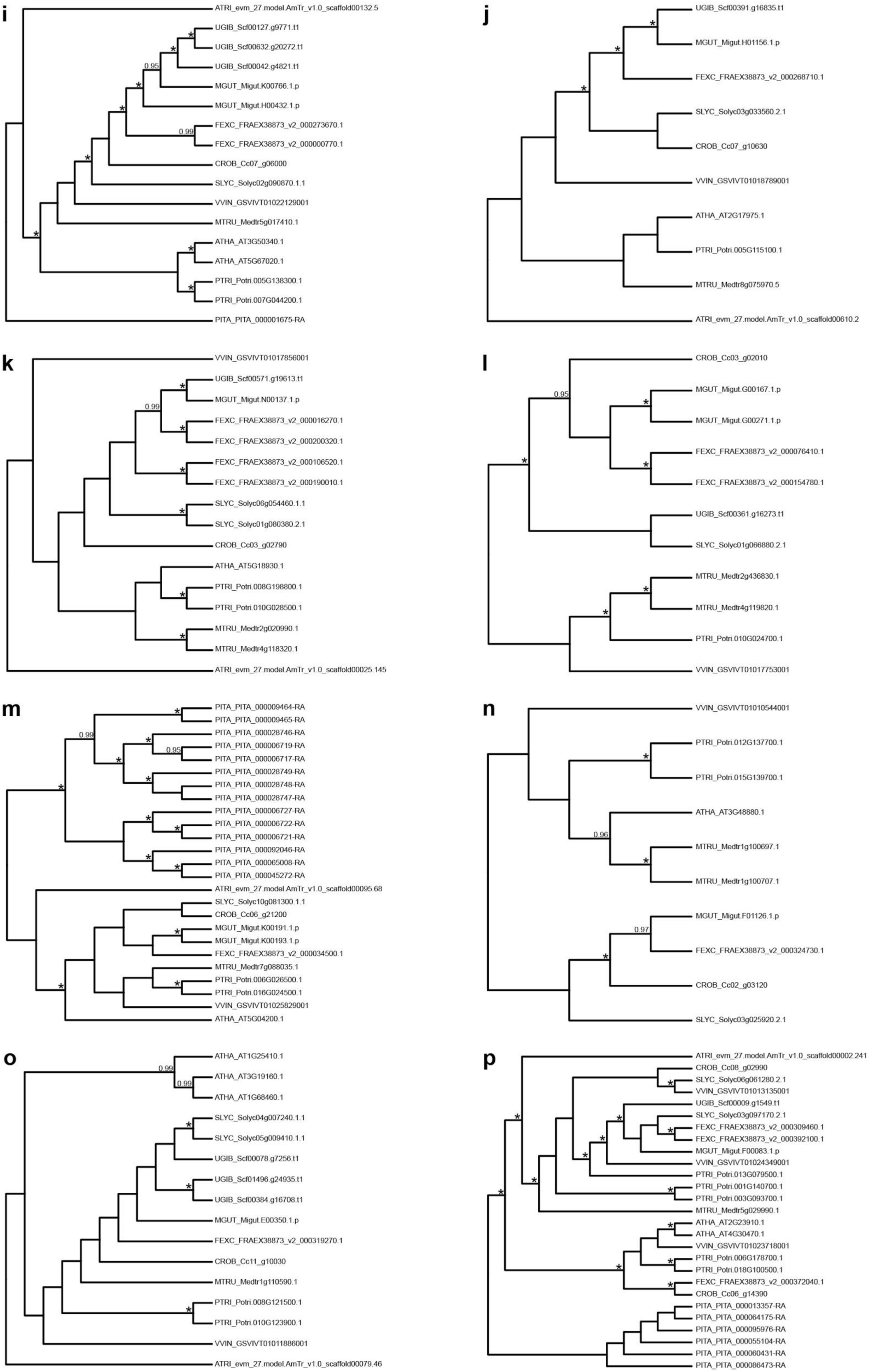
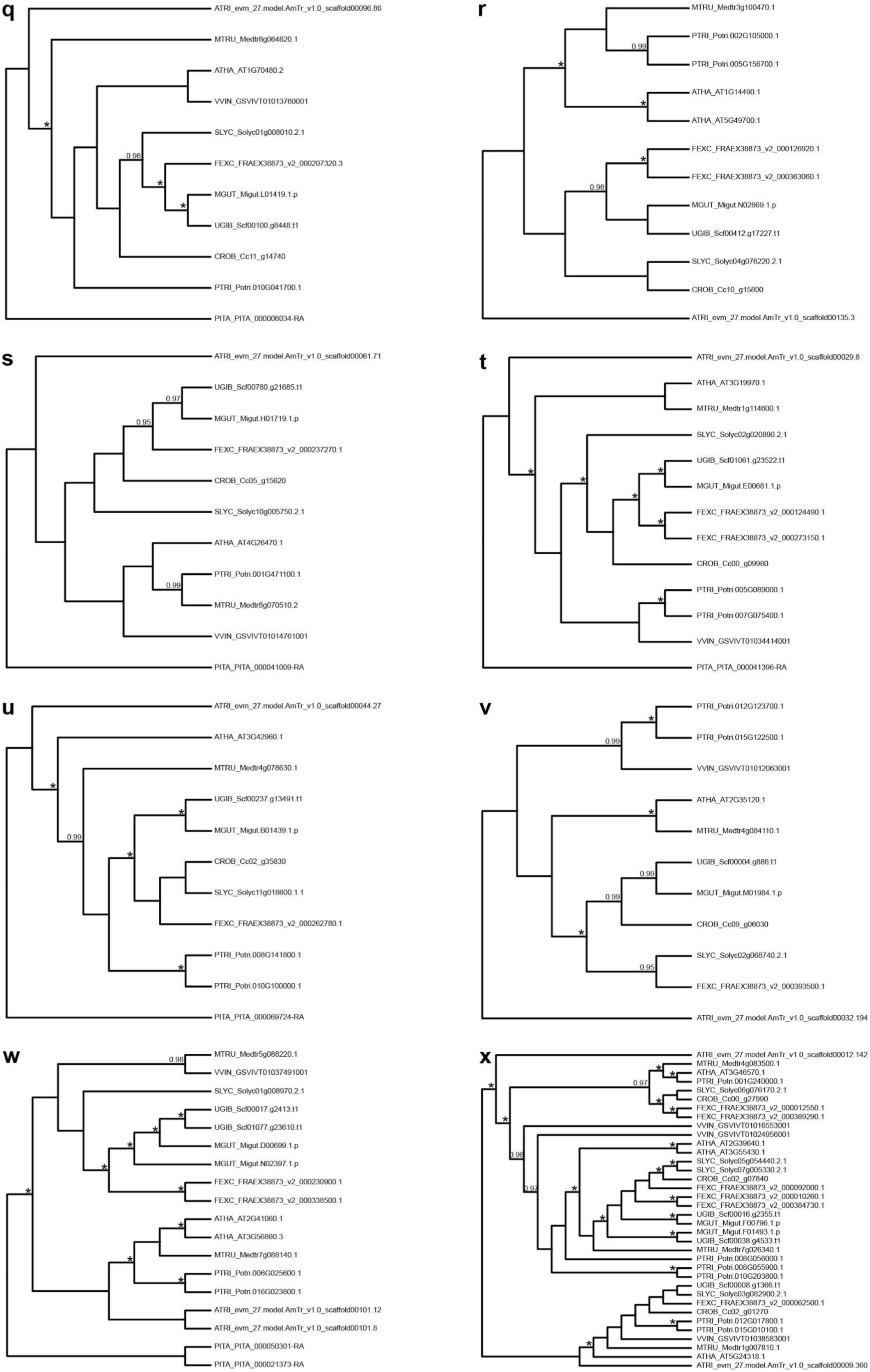
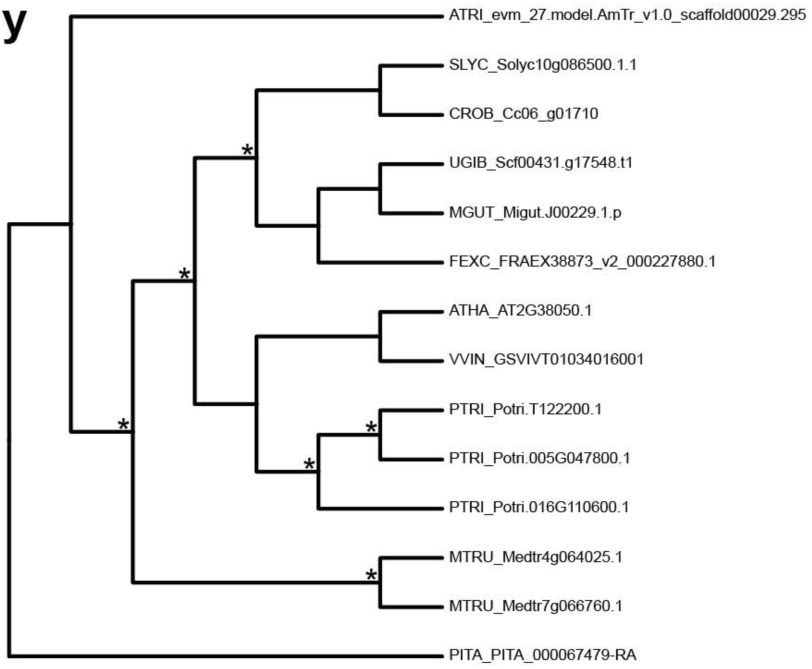
Gene-trees for OrthoMCL clusters including *Fraxinus excelsior* sequences from selected candidate genes. Gene-trees for OrthoMCL clusters including FRAEX38873_v2_000261700 (**a**), FRAEX38873_v2_000200250 (**b**), FRAEX38873_v2_000386580 (**c**), FRAEX38873_v2_000122730 (**d**), FRAEX38873_v2_000111170 (**e**), FRAEX38873_v2_000248940 (**f**), FRAEX38873_v2_000080400 (**g**), FRAEX38873_v2_000093260 (**h**), FRAEX38873_v2_000000770 (**i**), FRAEX38873_v2_000268710 (**j**), FRAEX38873_v2_000106520 (**k**), FRAEX38873_v2_000076410 (**l**), FRAEX38873_v2_000034500 (**m**), FRAEX38873_v2_000324730 (**n**), FRAEX38873_v2_000319270 (**o**), FRAEX38873_v2_000309460 (**p**), FRAEX38873_v2_000207320 (**q**), FRAEX38873_v2_000363060 (**r**), FRAEX38873_v2_000237270 (**s**), FRAEX38873_v2_000382300 (**t**), FRAEX38873_v2_000262780 (**u**), FRAEX38873_v2_000393500 (**v**), FRAEX38873_v2_000338500 (**w**), FRAEX38873_v2_000092000 (**x**) and FRAEX38873_v2_000227880 (**y**). Gene-trees were inferred with MrBayes and rooted on either *Pinus taeda*, *Amborella trichopoda* or midpoint rooted (in cases where branch lengths viewed when the tree was unrooted did not clearly indicate that sequences from either *P. taeda* or *A. trichopoda* represented suitable outgroups). Numbers above branches are posterior probabilities (PP) of ≥0.95; asterisks indicate nodes with PP=1. Species names are abbreviated as follows: ATHA, *Arabidopsis thaliana*; ATRI, *Amborella trichopoda*; CROB, *Coffea canephora*; FEXC, *Fraxinus excelsior*; MGUT, *Erythranthe guttata*; MTRU, *Medicago truncatula*; PITA, *Pinus taeda*; PTRI, *Populus trichocarpa*; SLYC, *Solanum lycopersicum*; UGIB, *Utricularia gibba*; VVIN, *Vitis vinifera*. Gene model names follow species abbreviations.

### Supplementary Notes

#### Supplementary Note 1

##### 1. Species-tree for Fraxinus

The primary concordance tree inferred with BUCKy (Fig. 2) suggests evidence for significant incongruence between individual gene-tree topologies, as demonstrated by the low concordance factors (CFs) for many of the nodes. Nevertheless, with the exception of sect. *Ornus*, all taxonomic sections represented by more than one taxon are resolved as monophyletic with a CF of ≥0.95, indicating that the majority of genes support these relationships. All species from sect. *Ornus* form a monophyletic group (CF=0.91) except for *F. griffithii*, which was also found to fall outside of the section on the basis of data from the plastid genome^68^.

#### Supplementary Note 2

##### 2. Results of grand-conv analyses

From grand-conv analyses of susceptible versus resistant *Fraxinus* lineages, we identified 64 loci with significant evidence of excess convergence (defined as loci with at least one amino acid site with a PP ≥0.90 of convergence and where the resistant branch pair has higher excess convergence than any other independent branch pair within the species-tree) in at least one of the three pairwise combinations of resistant lineages. The majority of loci (45) only show evidence of convergence between *F. mandshurica* and *F. platypoda*. Seven loci are only convergent between *F. mandshurica* and the sect. *Ornus* taxa (*F. baroniana*, *F. floribunda* and *Fraxinus* sp. D2006-0159), and nine between *F. platypoda* and the sect. *Ornus* taxa. Of the remaining three loci, one shows evidence for convergence in two of the three pairwise comparisons, with the final two loci having evidence of convergence in all three pairs of lineages. For all three loci with evidence of convergence between more than one pair of lineages, the same residues were inferred as being convergent in each case. Filtering of this initial list of 64 loci according to the criteria specified in the Methods (see section “Refining the initial list of candidate loci identified with grand-conv”), resulted in a refined list of 53 loci. Further details of each of these loci is provided in Supplementary Note 4.

#### Supplementary Note 3

##### 3. Evidence for loss-of-function mutations

Nineteen of our 53 loci have evidence for potential loss-of-function mutations (start codon losses, stop codon gains/premature stop codons or frameshift variants) that cannot be discounted by errors in gene model estimation (Supplementary Table 6). For ten of these loci, possible loss-of-function mutations are found only in susceptible taxa, for seven loci they are found only in resistant taxa, and for two loci they are found both in susceptible and resistant taxa. Three susceptible taxa (*F. latifolia*, *F. ornus* and *F. pennsylvanica* (susceptible genotype)) lack an allele without potential loss-of-function mutations (either they are homozygous for the mutation, or phasing indicates there are separate mutations on each allele) for two loci each (Supplementary Table 6) suggesting these genes could be non-functional, or generate a truncated protein product. For a single resistant taxon (*F. floribunda*) both alleles for OG60899 have potential loss-of-function mutations. In all other cases, there is either one allele that appears fully functional or it is uncertain whether multiple possible loss-of-function mutations are on the same or different alleles (Supplementary Table 6).

#### Supplementary Note 4

##### 4. Details of candidate loci

Seven candidate genes have putative roles relating to the phenylpropanoid biosynthesis pathway (OG853, OG15551, OG16673, OG19104, OG27080, OG40061 and OG64545), 15 have possible roles in perception and signalling that are relevant to defence response against insect herbivores (OG4469, OG11720, OG21033, OG23214, OG23284, OG24614, OG32176, OG33348, OG38407, OG39275, OG41448, OG41488, OG43828, OG47560 and OG50989) and two (OG16739 and OG37870) are candidates with putative roles related to hypersensitive like-response and programmed cell death. Further candidates also have potential roles relating to other aspects of defence response, including OG27693 and OG59564, which may play a role in response to oxidative stress, OG11013, an apparent member of a protein superfamily whose members have roles in plant stress and immunity, OG37560, a probable member of the *HIPP* gene family that may be involved in plant-pathogen interactions, and OG47629, which has a putative orthologue in *S. lycopersicum* that has been classified as a “disease-specific gene”.

Below we provide details of the individual candidate genes, including information from literature searches and results obtained from any phylogenetic analyses and protein modelling. Gene model numbers from the BATG0.5 reference genome for *F. excelsior* are shown in parentheses after the OMA group names.

###### OG6935 (FRAEX38873_v2_000218910)

Grand-conv detected evidence of a convergent methionine in all five resistant taxa, at a site where the other *Fraxinus* species in the analysis have a threonine (Supplementary Table 5). The top match for this gene in *Arabidopsis thaliana* is an RNA helicase family protein (AT3G62310) which is in the DExD/H-box RH family and has a putative orthologue in rice (OS03G0314100; both belong to the same OMA group (fingerprint: QWCVDFA), omabrowser.org/oma/info/ARATH24986/) that is drought inducible^102^. Expression of AT3G62310 has been shown to be induced in response to salt acclimation^103^.

###### OG11013 (FRAEX38873_v2_000029110)

Grand-conv detected evidence of a convergent arginine in all five resistant taxa, at a site where the other *Fraxinus* species in the analysis have a glutamine (Supplementary Table 5). The alignment of all sequences for OG11013 showed some susceptible species also have an arginine at the relevant amino acid site. However, the glutamine was only present in species that are susceptible to EAB, including *F. ornus* (sect. *Ornus*) and members of sect. *Melioides* (Supplementary Table 8), indicating possible convergence between a subset of the taxa with high susceptibility. It should be noted that both *F. pennsylvanica* individuals (susceptible and putatively resistant) have a glutamine at the relevant position. Among the outgroups, *Mimulus* had an arginine, *Olea* a glutamine and *Solanum* a histidine.The top match for this gene in *A. thaliana* is a tetratricopeptide repeat (TPR)-like superfamily protein encoding gene (AT3G49142). Although the function of AT3G49142 has not been fully characterised, the functions of some individual TPR proteins have been deduced, and include roles in plant stress, hormone signaling and immunity^104^.

###### OG27838 (FRAEX38873_v2_000134340)

Grand-conv detected evidence of a convergent arginine in the five resistant taxa, at a site where the susceptible *Fraxinus* species in the analysis have a glutamine. Although grand-conv identified the arginine as convergent between *F. mandshurica* and *F. platypoda*, and between *F. platypoda* and *F. baroniana, F. floribunda* and *Fraxinus* sp. D2006-0159, it was not detected as convergent in the pairwise comparison between *F. mandshurica* and *F. baroniana, F. floribunda* and *Fraxinus* sp. D2006-0159 (Supplementary Table 5). The top match for this gene in *A. thaliana* is *REDUCED DORMANCY 2* (AT2G38560), or *RDO2*, which is also known as *TRANSCRIPT ELONGATION FACTOR IIS (TFIIS)*. AT2G38560/*RDO2*/*TFIIS* encodes a transcription factor that is needed for RNA polymerase II (RNAPII) processivity, stimulating transcription elongation by RNAPII, and may also play a role in the control of alternative splicing^105^.

###### OG853 (FRAEX38873_v2_000200250)

Grand-conv detected evidence of a convergent phenylalanine in *F. mandshurica* and *F. platypoda*, at a site where the other *Fraxinus* species in the analysis have an isoleucine (Supplementary Table 5). The top match for this gene in *A. thaliana* is *REF4-RELATED 1* (AT3G23590), or *RFR1*; phylogenetic analysis of the OrthoMCL cluster containing both FRAEX38873_v2_000200250 and AT3G23590 indicates that they are orthologues (Supplementary Fig. 3b). AT3G23590/*RFR1* is also known as *MED5a* or *MED33a* and encodes a protein that belongs to the Mediator complex, which is a transcriptional regulator of nearly all cellular pathways^20^. AT3G23590/*RFR1*/*MED5a* acts in a partially redundant manner with its paralogue *REF4*/*MED5b* (AT2G48110) to limit phenylpropanoid accumulation^20^. Disruption of both the *MED5a* and *MED5b* genes in the lignin-deficient *ref8* mutant of *A. thaliana*, which has reduced activity of *REF8*/*CYP98A3* (AT2G40890; the gene encoding *p*-coumaroylshikimate 3’-hydroxylase (C3’H) and the best matching *A. thaliana* gene for one of our other candidates, OG15551/FRAEX38873_v2_000261700), restores lignin deposition to wild-type levels^21^. However, the disruption of the *MED5* genes does not restore the synthesis of guaiacyl and syringyl lignin subunits; the lignin content of the *med5a/5b ref8* triple mutant is almost entirely composed of *p*-hydroxyphenyl subunits, which account for < 2% of the lignin content in wild-type plants ^21^. It has also been found that *med5a/5b* loss-of-function mutant plants show increased expression of numerous genes involved in the phenylpropanoid biosynthesis pathway, including those encoding PAL, C4H, 4CL, C3’H, CCR, CAD and CSE, and that ‘phenylpropanoid biosynthesis’ was the most significantly enriched GO term for genes that were overexpressed in *med5a/5b* plants^21^. Although the direct targets of the *MED5* genes are yet to be identified, it has been suggested that *MED5a* and *MED5b* may limit phenylpropanoid biosynthesis via the transcriptional activation of genes that negatively regulate the pathway^106^. Further analysis of *med5a/5b* loss-of-function mutant plants found that genes with significantly decreased expression were enriched for GO terms related to defence response, such as “response to jasmonic acid”, “response to wounding” and “defence response to insect”, suggesting a role for *MED5a* and *MED5b* in regulation of defence response genes^22^. The orthologue of FRAEX38873_v2_000200250 in *S. lycopersicum* is Solyc01g080200 (Supplementary Fig. 3b), and has been shown to be upregulated in response to tomato yellow leaf curl virus^107^.

###### OG2897 (FRAEX38873_v2_000138720)

Grand-conv detected evidence of a convergent isoleucine and glutamic acid in *F. mandshurica* and *F. platypoda*, at sites where the other *Fraxinus* species in the analysis have a phenylalanine and glycine (Supplementary Table 5). The top match for this gene in *A. thaliana* is a gene encoding a tetratricopeptide repeat (TPR)-like superfamily protein (AT4G30825), also known as *PPRb*^108^. AT4G30825/*PPRb* is involved in nucleotide-excision repair, phosphorylation and regulation of transcription by RNA polymerase II, amongst other processes (www.arabidopsis.org/servlets/TairObject?id=500231848&type=locus). AT4G30825 is co-expressed with the disease resistance regulator *OVEREXPRESSOR OF CATIONIC PEROXIDASE3* (*OCP3*^108^), which also plays a role in drought tolerance^109^.

###### OG4372 (FRAEX38873_v2_000149840)

Grand-conv detected evidence of a convergent isoleucine in *F. mandshurica* and *F. platypoda*, at a site where the other *Fraxinus* species in the analysis have a serine or aspartic acid (Supplementary Table 5). The top match for this gene in *A. thaliana* is a gene encoding a transcription regulator NOT2/NOT3/NOT5 family protein (AT5G18230). NOT2 and NOT3/5 form a module that is part of the CCR4-NOT complex, which is a conserved multi-subunit complex that regulates gene expression at different levels^110^. The NOT2–3/5 module appears to be involved in coordination of transcriptional regulation and mRNA degradation, as well as assembly of cellular complexes such as the proteasome^110^.

###### OG4469 (FRAEX38873_v2_000386580)

Grand-conv detected evidence of a convergent threonine in *F. mandshurica* and *F. platypoda*, at a site where the other *Fraxinus* species in the analysis have a proline (Supplementary Table 5). The top match for this gene in *A. thaliana* is a gene encoding a protein kinase superfamily protein (AT5G24080); phylogenetic analysis of the OrthoMCL cluster containing FRAEX38873_v2_000386580 and AT5G24080 indicates they are likely to be orthologous (Supplementary Fig. 3c). The protein encoded by AT5G24080 is reported to be a G-type lectin S-receptor-like serine/threonine-protein kinase with an ATP binding site, belonging to the Ser/Thr protein kinase family (www.uniprot.org/uniprot/Q9FLV4). G-type lectin receptor kinases have various functions, including in self-incompatibility, with some having a role in plant defence^111,112^ such as in response to the tobacco hawk moth (*Manduca sexta*) in *Nicotiana attenuata*^113^ or to the brown planthopper (*Nilaparvata lugens*) in rice^114^. Extracellular ATP is associated with cell damage, and perception of ATP may play a role in defence response against insect herbivores via stimulation of the JA pathway^27^. AT5G24080 has also been named *AtG-LecRK-I.6* by Teixeira et al.^111^, who suggest it may be defective because its protein has an incomplete kinase domain and lacks some of the subdomains typically found in kinases. However, such proteins may still be functional even if they lack kinase activity^111^. Putative orthologues of OG4469/FRAEX38873_v2_000386580 in *S. lycopersicum* (Solyc03g007790 and Solyc06g036470) have been found to encode proteins that include the S-locus glycoprotein domain, transmembrane domain and kinase domains^111^, although their precise functions have not been characterised.

###### OG5539 (FRAEX38873_v2_000290300)

Grand-conv detected evidence of a convergent leucine in *F. mandshurica* and *F. platypoda*, at a site where the other *Fraxinus* species in the analysis have a phenylalanine (Supplementary Table 5). However, the leucine is also found in the alternate allele for susceptible *F. americana* (Supplementary Table 8). The top match for this gene in *A. thaliana* is *CHLORIDE CHANNEL G* (AT5G33280), *CLCG*. AT5G33280/*CLCG* is also known as *CBSCLC6* (www.uniprot.org/uniprot/P60300) and is suggested to function in chloride homeostasis during NaCl stress^115^.

###### OG7454 (FRAEX38873_v2_000218430)

Grand-conv detected evidence of a convergent histidine and lysine in *F. mandshurica* and *F. platypoda*, at sites where the other *Fraxinus* species in the analysis have arginine and glutamic acid (Supplementary Table 5). The top match for this gene in *A. thaliana* is *EMBRYO DEFECTIVE 3141* (AT5G50390), or *EMB3141*. AT5G50390/*EMB3141* is also known as *PCMP-H58* and encodes a pentatricopeptide repeat containing protein (www.uniprot.org/uniprot/Q9FK33); this gene appears to be required for embryo development^116^.

###### OG10762 (FRAEX38873_v2_000122730)

Grand-conv detected evidence of a convergent serine in *F. mandshurica* and *F. platypoda*, at a site where the other *Fraxinus* species in the analysis have an alanine (Supplementary Table 5). The top match for this gene in *A. thaliana* is *WITH NO LYSINE (K) KINASE 4* (AT5G58350), or *WNK4*. Phylogenetic analysis of the OrthoMCL cluster containing both FRAEX38873_v2_000122730 and AT5G58350 indicates that they are orthologues (Supplementary Fig. 3d). AT5G58350/*WNK4* is also known as *ZIK2* and encodes a serine/threonine kinase^117^. The function of AT5G58350/*WNK4* in not yet known, although its expression is influenced by circadian rhythms and other members of the *WNK* gene family in *A. thaliana* are involved in the regulation of flowering time^117^. It has also been suggested that plant *WNK* genes are involved in enhancing the oxidative defence system^117^. GO terms have been associated with AT5G58350/*WNK4* via identification of functional gene modules^118^, namely: CELLULAR RESPONSE TO PHOSPHATE STARVATION, GALACTOLIPID BIOSYNTHETIC PROCESS, CELLULAR RESPONSE TO WATER DEPRIVATION and ORGAN SENESCENCE (see http://bioinformatics.psb.ugent.be/cig_data/plant_modules/createResultsHTML2.php?geneID=AT5G58350&moduleTypes=repModules). The orthologue of FRAEX38873_v2_000122730 in *S. lycopersicum* appears to be Solyc06g082470 (Supplementary Fig. 3d) and is also known as SlMAPKKK42^119^. The function of Solyc06g082470/SlMAPKKK42 has not been fully characterised, but it has been suggested that SlMAPKKK genes in general might have a role in plant hormone signalling in relation to development and defence response^119^.

###### OG11720 (FRAEX38873_v2_000155730)

Grand-conv detected evidence of a convergent valine in *F. mandshurica* and *F. platypoda*, at a site where the other *Fraxinus* species in the analysis have an isoleucine (Supplementary Table 5). The top match for this gene in *A. thaliana* is *NITRATE TRANSPORTER 1.5* (AT1G32450), which is also known as *NRT1.5* or *NPF7.3*. AT1G32450/*NRT1.5* is a transmembrane nitrate transporter and has been shown to be involved in transport of nitrate from the root to shoot^32^. In addition to nitrate, some members of the NRT1 family (also known as the PTR or NPF family) are also involved in transport of some phytohormones and defence compounds^120^. Indeed, many of the 53 members of the NRT1 family in *A. thaliana* have multiple substrates and may be able transport jamonate, abscisic and/or gibberellins as well as nitrates, or other substrates^121^.

###### OG13887 (FRAEX38873_v2_000015960)

Grand-conv detected evidence of a convergent lysine n *F. mandshurica* and *F. platypoda*, at a site where the other *Fraxinus* species in the analysis have an arginine (Supplementary Table 5). The top match for this gene in *A. thaliana* i*s 54 CHLOROPLAST PROTEIN* (AT5G03940), or *54CP*, and is also known as *cpSRP54*. AT5G03940/*54CP*/*cpSRP54* encodes the 54kDa subunit of the chloroplast signal recognition particle and an *A. thaliana* mutant (*cbd*) with a T-DNA insertion in the gene has lower chlorophyll and carotenoid content compared with wild type plants, shows defects in chloroplast development and is deficient in abscisic acid^122^. The *cbd* mutant also shows evidence of cell death in leaves; it has been suggested that a reduced ability to quench reactive oxygen species as a result the lower levels of carotenoids in the mutant may contribute to the observed cell death^122^.

###### OG15551 (FRAEX38873_v2_000261700)

Grand-conv detected evidence of four convergent amino acids in *F. mandshurica* and *F. platypoda*, valine, isoleucine, isoleucine and methionine, where the other *Fraxinus* species in the analysis have an isoleucine at the first site and leucine at the other three (Supplementary Table 5). The top match for this gene in *A. thaliana* is the *CYP98A3* gene (AT2G40890; also known as *REF8*). In *A. thaliana* this gene is at a critical bottleneck in the phenylpropanoid pathway^18^. When *CYP98A3*/*REF8* is mutated in *A. thaliana*, it results in reduced lignin content, changes in lignin composition, a lack of soluble sinapoyl esters, and accumulation of flavonol glycosides^18^. Another study found a more than 90% reduction in the level of scopoletin and scopolin in the roots of *A. thaliana* mutants with T-DNA insertions in this gene compared with that found in wild type plants^123^. Phylogenetic analysis of the OrthoMCL cluster containing AT2G40890 and FRAEX38873_v2_000261700 indicates that they are close paralogues, rather than orthologues and suggests that the *Fraxinus* gene lacks a direct orthologue in *A. thaliana* (Supplementary Fig. 3a). There appears to be a small family of *CYP98A3-like* genes in *F. excelsior*, including FRAEX38873_v2_000095940 and FRAEX38873_v2_000127720 as well as FRAEX38873_v2_000261700 (Supplementary Fig. 3a). In a predicted protein model for OG15551/FRAEX38873_v2_000261700 all four variable sites identified by grand-conv are in close proximity to the active site (Fig. 3).

###### OG16673 (FRAEX38873_v2_000150510)

Grand-conv detected evidence of a convergent isoleucine in *F. mandshurica* and *F. platypoda*, at a site where the other *Fraxinus* species in the analysis have a valine (Supplementary Table 5). However, the isoleucine is also found in the alternate allele for susceptible *F. ornus* (Supplementary Table 8). The top match for this gene in *A. thaliana* is *Β-GLUCOSIDASE 41* (AT5G54570; also known as *BGLU41*). *BGLU41* belongs to the glycoside hydrolase family 1 (GH1) in *A. thaliana*, members of which function in processes such as chemical defence against herbivory, lignification, cell wall modification and control of phytohormone levels^23,124^. A number of *cis*-elements have been found in the region upstream of AT5G54570/*BGLU41*, including those that are responsive to abscisic acid and gibberellins^124^. The putative orthologue of OG16673/FRAEX38873_v2_000150510 in tomato is Solyc07g063880; this gene is differentially expressed in the *ovate* mutant and it had been suggested that its downregulation could be responsible for decreased glucose content in the fruit of *ovate* plants^125^. There has been debate regarding whether glucose should be considered as part of defence response, because it can reduce herbivore performance^126^.

###### OG17252 (FRAEX38873_v2_000111170)

Grand-conv detected evidence of a convergent asparagine and methionine in *F. mandshurica* and *F. platypoda*, at sites where the other *Fraxinus* species in the analysis have aspartic acid and threonine (Supplementary Table 5). The top match for this gene in *A. thaliana* is AT1G50460 (*HEXOKINASE-LIKE 1*/*HKL1*), which has a role in abiotic stress response in addition to its other functions (growth, root hair development^127^). AT4G29130 (*HXK1*) belongs to the same OrthoMCL cluster as AT1G50460, along with three other *A. thaliana* genes, and has response to oxidative stress and pathogen resistance among its functions^127^. Phylogenetic analysis of the OrthoMCL cluster indicates that FRAEX38873_v2_000111170 is orthologous to both AT1G50460/*HKL1* and another *A. thaliana* gene, AT3G20040 (*HEXOKINASE-LIKE 2*/*HKL2*; Supplementary Fig. 3e). AT3G20040/*HKL2* is also known as *ATHXK4*, but its function is apparently unknown^127^.

###### OG19104 (FRAEX38873_v2_000248940)

Grand-conv detected evidence of a convergent leucine in *F. mandshurica* and *F. platypoda*, at a site where the other *Fraxinus* species in the analysis have a serine (Supplementary Table 5). The top match for this gene in *A. thaliana* is AT4G01240, a gene encoding a S-adenosyl-L-methionine-dependent methyltransferases superfamily protein. Phylogenetic analysis of the OrthoMCL cluster containing FRAEX38873_v2_000248940 and AT4G01240 indicates they are likely to be orthologous (Supplementary Fig. 3f). The function of AT4G01240 has not been characterised but the gene was found to be within a region related to aphid feeding performance from a GWAS conducted in *A. thaliana*^128^. Plant S-adenosyl-L-methionine-dependent methyltransferases are also key enzymes in the phenylpropanoid, flavonoid and other metabolic pathways^129^. Methyltransferases in maize may function in generating volatile methyl esters (which function in plant defence) in response to herbivore (African cotton leaf worm) damage^130^. The serine variant in *Fraxinus* is a putative phosphorylation site (Supplementary Table 5); it was not possible to generate a predicted model for OG19104/FRAEX38873_v2_000248940 and therefore the position of the putative phosphorylation site within the structure of the protein is unknown.

###### OG20252 (FRAEX38873_v2_000397500)

Grand-conv detected evidence of a convergent valine in *F. mandshurica* and *F. platypoda*, at a site where the other *Fraxinus* species in the analysis have an isoleucine (Supplementary Table 5). However, the gene-tree inferred for OG20252 with the codon for the valine/isoleucine variant excluded groups sect. *Fraxinus* (all of which have the valine; Supplementary Table 8) and *F. platypoda* together (PP=0.98; Supplementary Fig. 1a) in conflict with the species-tree (Fig. 2), indicating that the evidence of convergence found by grand-conv is likely due to introgression or incomplete lineage sorting. The top match for this gene in *A. thaliana* is *BROMODOMAIN 4* (AT1G61215) which encodes a bromodomain protein with a DNA binding motif. The exact function of the bromodomain is unclear, but it may be involved in protein-protein interactions (http://www.ebi.ac.uk/interpro/entry/IPR001487). The putative orthologue of OG20522/FRAEX38873_v2_000397500 in *S. lycopersicum* is Solyc02g078810.2 and is annotated with a MYB-like DNA binding domain in PhytoMine (https://phytozome.jgi.doe.gov/phytomine/report.do?id=285298920&trail=%7c285298920).

###### OG20859 (FRAEX38873_v2_000138760)

Grand-conv detected evidence of a convergent alanine and glutamic acid in *F. mandshurica* and *F. platypoda*, at sites where the other *Fraxinus* species in the analysis have threonine or serine and aspartic acid (Supplementary Table 5). The top match for this gene in *A. thaliana* is AT5G57830, which encodes a zein-binding protein of unknown function.

###### OG23214 (FRAEX38873_v2_000080400)

Grand-conv detected evidence of a convergent valine and isoleucine in *F. mandshurica* and *F. platypoda*, at sites where the other *Fraxinus* species in the analysis have a methionine and threonine (Supplementary Table 5). The top match for this gene in *A. thaliana* is a gene encoding an eukaryotic aspartyl protease family protein (AT1G64830). Aspartic proteases are involved in many biological processes, including senescence, stress responses, and programmed cell death^131^. Phylogenetic analysis of the OrthoMCL cluster that contains FRAEX38873_v2_000080400 indicates it is orthologous to both AT1G64830 and another *A. thaliana* gene, AT5G33340 (*CDR1*/*CONSTITUTIVE DISEASE RESISTANCE 1*; see Supplementary Fig. 3g). AT5G33340/*CDR1* is an extracelluar aspartic protease that is involved in disease signalling via salicylic acid-dependent inducible resistance^132^; overexpression of AT5G33340/*CDR1* leads increased expression of other defence-related genes and enhanced resistance to a bacterial pathogen^132^. Salicylic acid is more often linked with defense against biotrophic pathogens, but is involved in response to oviposition by insect herbivores^17^ and plays a key role in pathogen-triggered programmed cell death^33^.

###### OG24614 (FRAEX38873_v2_000093260)

Grand-conv detected evidence of a convergent isoleucine in *F. mandshurica* and *F. platypoda*, at a site where the other *Fraxinus* species in the analysis have a threonine (Supplementary Table 5). The top match for this gene in *A. thaliana* is a gene encoding a zinc finger protein (AT5G54630). However, phylogenetic analysis of the relevant OrthoMCL cluster indicates another *A. thaliana* gene, AT4G27240, is equally closely related to FRAEX38873_v2_000093260, with both appearing to be paralogues of the *F. excelsior* gene (Supplementary Fig. 3h). AT4G27240 also encodes a zinc finger protein; both AT4G27240 and AT5G54630 are transcription factors^133^, but their functions are not fully characterised. AT5G54630 is expressed in response to brassinolide^134^, a brassinosteroid, which have a crucial role in regulating the growth-immunity trade-off^135^; transcriptions factors act both up and downstream of phytohormone signalling to regulate defence responses^27^. One of the susceptible taxa included in our convergence analyses, *F. ornus*, may lack a functional copy of this gene (Supplementary Table 6).

###### OG24969 (FRAEX38873_v2_000395930)

Grand-conv detected evidence of a convergent serine in *F. mandshurica* and *F. platypoda*, at a site where the other *Fraxinus* species in the analysis have a threonine (Supplementary Table 5). The top match for this gene in *A. thaliana* is *ROH1* (AT1G63930). AT1G63930 has been suggested to play a role in seed coat development, but it is possible that it does not have this function in wild-type *A. thaliana*^136^. One of the susceptible taxa included in our convergence analyses, *F. pennsylvanica*, may lack a functional copy of this gene (Supplementary Table 6).

###### OG26964 (FRAEX38873_v2_000042410)

Grand-conv detected evidence of a convergent histidine and methionine in *F. mandshurica*, at sites where the other *Fraxinus* species in the analysis have an arginine and threonine (Supplementary Table 5). The top match for this gene in *A. thaliana* is *C5orf35* (AT5G23200); the function of this gene has not been characterised.

###### OG27080 (FRAEX38873_v2_000000770)

Grand-conv detected evidence of a convergent aspartic acid and methionine in *F. mandshurica* and *F. platypoda*, at sites where the other *Fraxinus* species in the analysis have an asparagine and leucine (Supplementary Table 5). The top match for this gene in *A. thaliana* is the uncharacterised gene AT5G67020. However, phylogenetic analysis of the OrthoMCL cluster containing FRAEX38873_v2_000000770 indicates it is orthologous to both AT5G67020 and AT3G50340 (Supplementary Fig. 3i). Although the function of AT5G67020 is unknown it is suggested to be a possible target of members of the *R2R3-MYB* gene family, which appear to control accumulation of flavonols^137^. AT5G67020 has reduced expression in a mutant where three *R2R3-MYB* genes have been knocked out; other genes that also show a reduction in expression include those encoding known flavonoid biosynthesis enzymes, as well as the *4CL3* gene (AT1G65060) that functions in the general phenylpropanoid pathway^137^. AT3G50340 is also not fully characterised but has been identified as an auxin-inducible gene^138^ and is also suggested to be regulated by drought, abscisic acid and jasmonic acid^139^.

###### OG28712 (FRAEX38873_v2_000047370)

Grand-conv detected evidence of a convergent histidine and methionine in *F. mandshurica* and *F. platypoda*, at sites where the other *Fraxinus* species in the analysis have alanine and threonine (Supplementary Table 5). The top match for this gene in *A. thaliana* is *PROTOCHLOROPHYLLIDE OXIDOREDUCTASE A* (AT5G54190), also known as *PORA*. AT5G54190/*PORA* encodes light-dependent NADPH:protochlorophyllide oxidoreductase A and is involved in chlorophyll biosynthesis (www.arabidopsis.org/servlets/TairObject?type=locus&name=AT5G54190). The putative orthologue of OG28712/FRAEX38873_v2_000047370 in *S. lycopersicum* is Solyc12g013710; this gene has been identified as a mycorrhiza-regulated gene^140^, and is upregulated in plants colonised with a mycorrhizal fungus along with a number of other genes involved in photosynthesis^140^. Expression of Solyc12g013710 is repressed by overexpression of *SlRBZ* (Solyc03g033560^141^) which is the putative orthologue of another of the candidate loci, OG32176. One of the susceptible taxa included in our convergence analyses, *F. pennsylvanica*, may lack a functional copy of this gene (see Supplementary Table 6).

###### OG30208 (FRAEX38873_v2_000228080)

Grand-conv detected evidence of a convergent histidine and valine in *F. mandshurica* and *F. platypoda*, at sites where the other *Fraxinus* species in the analysis have a glutamine and alanine (Supplementary Table 5). The top match for this gene in *A. thaliana* is a gene encoded an uncharacterised transmembrane protein, DUF677 (AT1G20180). AT1G20180 is upregulated in two *A. thaliana* autophagy mutant lines in response to differing amounts of nitrogen^142^; other genes upregulated at the same time were predominantly involved in response to biotic stress, chemical and abiotic stress and salicylic acid^142^. The FRAEX38873_v2_000228080 reference gene model from *F. excelsior* is truncated relative to the other sequences in the OG, which may be due to a misassembly introducing a premature stop codon or might indicate that the *F. excelsior* reference individual does not have a full-length copy of the gene (Supplementary Table 5). One of the susceptible taxa included in our convergence analyses, *F. ornus*, may lack a functional copy of this gene (Supplementary Table 6).

###### OG32176 (FRAEX38873_v2_000268710)

Grand-conv detected evidence of a convergent histidine and leucine in *F. mandshurica* and *F. platypoda*, at sites where the other *Fraxinus* species in the analysis have an glutamine and serine (Supplementary Table 5). The top match for this gene in *A. thaliana* is *STRESS ASSOCIATED RNA-BINDING PROTEIN 1* (AT2G17975), also known as *SRP1*; FRAEX38873_v2_000268710 and AT2G17975/*SRP1* appear orthologous on the basis of phylogenetic analysis of the OrthoMCL cluster to which they belong (Supplementary Fig. 3j). AT2G17975/*SRP1* is an RNA-binding protein involved in the post-transcriptional regulation of abscisic acid (ABA) signalling via the modulation of *ABI* genes^143^; expression of AT2G17975 is down regulated in response to ABA and abiotic stress^143^. The apparent orthologue of OG32176/FRAEX38873_v2_000268710 in *S. lycopersicum* is Solyc03g033560 (Supplementary Fig. 3j); this gene is also known as *SlRBZ* and is a RanBP2-type zinc finger protein gene^141^. Mutants that overexpress *SlRBZ* have a dwarf phenotype and are chlorotic^141^. Overexpression of *SlRBZ* also leads to decreased expression of photosynthesis genes, including Solyc12g013710 which is the putative orthologue of another of the candidate loci, OG28712; gibberellic acid (GA) biosynthesis genes were also repressed^141^. It is suggested that *SlRBZ* regulates the formation of chloroplasts, and as a result also controls chlorophyll, carotenoid and GA biosynthesis, which take place in the chloroplast^141^. GA may play a role in regulating anti-herbivore defense^27^.

###### OG33348 (FRAEX38873_v2_000106520)

Grand-conv detected evidence of a convergent serine in *F. mandshurica* and *F. platypoda*, at a site where the other *Fraxinus* species in the analysis have an asparagine (Supplementary Table 5). The top match for this gene in *A. thaliana* is *S-ADENOSYLMETHIONINE DECARBOX*YLA*SE 4* (AT5G18930), or *SAMDC4*. AT5G18930/*SAMDC4* is also known as *BUSHY AND DWARF 2* (*BUD2)* and is involved in the synthesis of S-adenosylmethioninamine from S-adenosyl-L-methionine within the S-adenosylmethioninamine biosynthesis pathway (https://www.uniprot.org/uniprot/Q3E9D5). Phylogenetic analysis of the OrthoMCL cluster including FRAEX38873_v2_000106520 and AT5G18930 indicates they are likely to be orthologous, albeit the relevant node is not well supported (PP<0.95; Supplementary Fig. 3k). AT5G18930/*SAMDC4*/*BUD2* is essential for biosynthesis of the polyamines spermidine and spermine^144^; polyamines in higher plants can be involved in mediating biotic and abiotic stress responses, such as pathogen infection, osmotic stress and wounding ^144^. Spermine in particular is suggested to play a role in defence response signalling^36^. Loss of function of AT5G18930/*SAMDC4*/*BUD2* alters growth and development^144^ and analysis of its promoter region found elements that are associated with response to auxin, dehydration, drought, salt, cold, wounding, defence related gene expression and wound response, amongst other physiological responses^145^.

###### OG37560 (FRAEX38873_v2_000076410)

Grand-conv detected evidence of a convergent alanine in *F. mandshurica* and *F. platypoda*, at a site where the other *Fraxinus* species in the analysis have a proline (Supplementary Table 5). The top match for this gene in *A. thaliana* is a gene encoding a heavy metal transport/detoxification superfamily protein (AT5G27690). AT5G27690 has also been called *AthHIPP36* or *AthHIP36* and is a member of a heavy metal-associated isoprenylated plant protein (HIPP) family^146^. HIPPs may have roles in heavy metal homeostasis and detoxification, transcriptional responses to cold and drought, and plant-pathogen interactions^146^. However, FRAEX38873_v2_000076410 does not belong to the same OrthoMCL cluster as AT5G27690, and appears not to have an orthologue in *A. thaliana*. FRAEX38873_v2_000076410 does belong to the same OrthoMCL cluster as, and appears orthologous to (Supplementary Fig. 3l), the *P. trichocarpa* gene Potri.010G024700 (also called POPTR_0010s02550), which encodes a putative copper (Cu) transport protein (https://phytozome.jgi.doe.gov/phytomine/report.do?id=49365037&trail=%7c49365037) and belongs to the HIPP family (*PtrHIP43*^146^). It has been suggested that mechanisms for signalling and responding to Cu stress could overlap with those involved in biotic stress, possibly relating to both types of stress causing the production of reactive oxygen species^147^; exposure to high Cu can prime maize plants for increased production of volatile organic compounds and faster phytohormone signalling upon subsequent herbivory by an insect herbivore^147^. Moreover, copper transport genes have been implicated in the regulation of defence response to whitefly (*Bemisia tabaci*) in cotton^148^. One of the susceptible taxa included in our convergence analyses, *F. latifolia*, may lack a functional copy of this gene (see Supplementary Table 6).

###### OG37870 (FRAEX38873_v2_000034500)

Grand-conv detected evidence of a convergent glutamine in *F. mandshurica* and *F. platypoda*, at a site where the other *Fraxinus* species in the analysis have an arginine (Supplementary Table 5). The top match for this gene in *A. thaliana* is *METACASPASE 9* (AT5G04200), or *AtMC9*. Results of phylogenetic analysis of the OrthoMCL cluster including FRAEX38873_v2_000034500 and AT5G04200/*AtMC9* are compatible with the inference that these genes are orthologues (although the topology does not fit exactly with expected species relationships, suggesting the *A. thaliana* and *F. excelsior* sequences may be from different paralogues, this is not well supported; Supplementary Fig. 3m). AT5G04200/*AtMC9* plays a role in controlling autophagy in tracheary elements, ensuring that cell death does not spread to surrounding non-target cells^149^. The site with evidence of convergence in *Fraxinus* is located within a linker region between p20 and p10-like domains of the AtMC9 protein^150^; the linker region appears to be important to the function of AtMC9, as its removal causes a significant reduction in the enzyme’s activity^150^. Two poplar genes, Potri.006G026500.1 and Potri.016G024500.1, appear orthologous to FRAEX38873_v2_000034500 (Supplementary Fig. 3m) and are known as *PtMC13* and *PtMC14*, respectively^42^. Both poplar genes are suggested to play a role in controlling cell death in xylem elements^42^.

###### OG38407 (FRAEX38873_v2_000324730)

Grand-conv detected evidence of a convergent aspartic acid in *F. mandshurica* and *F. platypoda*, at a site where the other *Fraxinus* species in the analysis have an asparagine (Supplementary Table 5). The top match for this gene in *A. thaliana* is *SNC1-INFLUENCING PLANT E3 LIGASE REVERSE GENETIC SCREEN 4* (AT3G48880), or *SNIPER4*, which encodes an F-box protein. Phylogenetic analysis of their OrthoMCL cluster indicates that FRAEX38873_v2_000324730 and AT3G48880/*SNIPER4* are likely be to orthologues (Supplementary Fig. 3n). AT3G48880/*SNIPER4* was found to be a positive regulator of effector triggered immunity (ETI) in *A. thaliana* and has a role in maintaining the balance of immune response by forming part of a complex directing the MUSE13 and MUSE14 proteins for degradation^28^. The targeted degradation of MUSE13/14 ensures sufficient levels of SNC1, an immune sensor, are available to trigger defence response against a bacterial pathogen^28^.

###### OG38543 (FRAEX38873_v2_000362960)

Grand-conv detected evidence of a convergent aspartic acid in *F. mandshurica* and *F. platypoda*, at a site where the other *Fraxinus* species in the analysis have an asparagine (Supplementary Table 5). The top match for this gene in *A. thaliana* is a gene encoding a cyclin-like protein (AT3G19650). Specific functional information is lacking for AT3G19650, however, cyclins regulate activity of cyclin-dependent kinases, which play critical roles in the control of cell cycle progression^151^.

###### OG39275 (FRAEX38873_v2_000319270)

Grand-conv detected evidence of a convergent glutamic acid in *F. mandshurica* and *F. platypoda*, at a site where the other *Fraxinus* species in the analysis have a glutamine (Supplementary Table 5). The top match for this gene in *A. thaliana* is *ISOPENTENYLTRANSFERASE 6* (AT1G25410), also called *IPT6*. Phylogenetic analysis of the OrthoMCL cluster including FRAEX38873_v2_000319270 indicates that, in addition to AT1G25410/*IPT6*, it is equally closely related to two other *A. thaliana* genes (AT1G68460/*IPT1* and AT3G19160/*IPT8*) and may be orthologous to all three (Supplementary Fig. 3o). AT1G68460/*IPT1*, AT1G25410/*IPT6* and AT3G19160/*IPT8* belong to a plant-specific ATP/ADP IPT clade of genes, which synthesize iP- and tZ-type cytokinins^152^. Cytokinins play a role in plant growth, defence response and immunity; they help to mediate trade-offs between growth and defence^31,153^. The FRAEX38873_v2_000319270 reference gene model from *F. excelsior* appears to be incorrect, possibly due to a misassembly causing an internal inversion (Supplementary Table 5).

###### OG40061 (FRAEX38873_v2_000309460)

Grand-conv detected evidence of a convergent asparagine in *F. mandshurica* and *F. platypoda*, at a site where the other *Fraxinus* species in the analysis have a serine (Supplementary Table 5). The top match for this gene in *A. thaliana* is a gene encoding a NAD(P)-binding Rossmann-fold superfamily protein (AT2G23910), which is reported to be involved in lignin biosynthetic process and response to karrikin and to cinnamoyl-CoA reductase activity and oxidoreductase activity (https://www.arabidopsis.org/servlets/TairObject?name=AT2G23910&type=locus). However, phylogenetic analysis indicates that FRAEX38873_v2_000309460 is paralogous to AT2G23910, along with AT4G30470 (another NAD(P)-binding Rossmann-fold superfamily protein), and may lack a direct orthologue in *A. thaliana* (Supplementary Fig. 3p). AT2G23910 is also known as *CCR(Cinnamoyl-CoA reductase)6*^154^, *CCR-like8*^155^ or *CCRL9*^156^ and is co-expressed with genes involved in flavonoid metabolism^156^. AT4G30470 is also known as *CCR-like6*^155^ and, in common with the best matching *A. thaliana* gene for another of our candidates (OG27080, see above), is a possible target of transcription factors in the *R2R3-MYB* gene family that appear to control accumulation of flavonols^137^. The apparent orthologue of OG40061/FRAEX38873_v2_000309460 in *S. lycopersicum* is Solyc03g097170 (Supplementary Fig. 3p); Solyc03g097170 was found to be upregulated in response to gibberellin and has been defined as a DELLA-dependent gene^157^. The serine variant in *Fraxinus* is a putative phosphorylation site (Supplementary Table 5), and is found on the exterior of the protein in the predicted model for OG40061/FRAEX38873_v2_000309460 in close proximity to the putative active site (Supplementary Fig. 2b). The FRAEX38873_v2_000309460 reference gene model from *F. excelsior* appears to be truncated on the 3′ end due to the genome assembly being incomplete (Supplementary Table 5).

###### OG41448 (FRAEX38873_v2_000207320)

Grand-conv detected evidence of a convergent alanine in *F. mandshurica* and *F. platypoda*, at a site where the other *Fraxinus* species in the analysis have glycine (Supplementary Table 5). The top match for this gene in *A. thaliana* is *JASSY* (AT1G70480); phylogenetic analysis of the OrthoMCL cluster including FRAEX38873_v2_000207320 and AT1G70480/*JASSY* indicates they are likely to be orthologous (Supplementary Fig. 3q). AT1G70480/*JASSY* encodes a chloroplast outer membrane protein that is involved in the transport of the JA precursor 2-oxophytodienoic acid (OPDA) from the chloroplast^158^. AT1G70480/*JASSY* loss-of-function of mutants show increased susceptibility to a fungal pathogen compared with wild-type plants and also lacked activation of JA-responsive genes following wounding, suggesting that the gene is essential for initiation of JA-signalling pathways^158^. Further analysis revealed that the defects observed in the mutants result from a failure of JA accumulation in the absence of AT1G70480/*JASSY* expression^158^. JASSY belongs to the Bet v1-like protein superfamily^158^; a related protein has been proposed to play a role in resistance to EAB in *F. mandshurica*^75^.

###### OG41488 (FRAEX38873_v2_000363060)

Grand-conv detected evidence of a convergent threonine in *F. mandshurica* and *F. platypoda*, at a site where the other *Fraxinus* species in the analysis have alanine (Supplementary Table 5). The top match for this gene in *A. thaliana* is *AT-HOOK MOTIF NUCLEAR LOCALIZED PROTEIN 17* (AT5G49700), or *AHL17*. Phylogenetic analysis of the OrthoMCL cluster including FRAEX38873_v2_000363060 indicates that it is likely to be orthologous to both AT5G49700/*AHL17* and AT1G14490/*AHL28* (Supplementary Fig. 3r). AT5G49700/*AHL17* and belong AT1G14490/*AHL28* to the AHL gene family in *A. thaliana*, members of which are involved in modulating plant growth and development^159^. *AHL* genes may also play a role in regulating homeostasis of gibberellins, jasmonic acid and cytokinins, and some *AHL* genes may be involved in regulating plant defence response (reviewed by Zhao et al.,^160^). The apparent orthologue of OG41488/FRAEX38873_v2_000363060.1 in *S. lycopersicum* is Solyc04g076220 and is also known as *AHL17a*^161^ (Howden et al., 2017); the potential role of Solyc04g076220/*AHL17a* in immunity to a fungal pathogen in tomato has been investigated, but over-expression this gene did not visibly impact immunity^161^.

###### OG46977 (FRAEX38873_v2_000304400)

Grand-conv detected evidence of a convergent arginine in *F. mandshurica* and *F. platypoda*, at a site where the other *Fraxinus* species in the analysis have a serine (two taxa with a glutamine instead were found to have errors in their predicted gene models; Supplementary Table 5). The top match for this gene in *A. thaliana* is a transmembrane proteins 14C gene (AT3G43520). AT3G43520 is also known as *FATTY ACID EXPORT 2* (*AtFAX2*) and may be involved in free fatty acids export from the plastids (https://www.uniprot.org/uniprot/Q94A32). The FRAEX38873_v2_000304400 reference gene model from *F. excelsior* appears to be incorrect on the 3′ end, possibly due to a misassembly (Supplementary Table 5).

###### OG47629 (FRAEX38873_v2_000173180)

Grand-conv detected evidence of a convergent aspartic acid in *F. mandshurica* and *F. platypoda*, at a site where the other *Fraxinus* species in the analysis have a glycine (Supplementary Table 5). The top match for this gene in *A. thaliana* is a gene encoding a MIZU-KUSSEI-like protein of unknown function (AT2G37880). AT2G37880 has been identified as an early high-light responsive gene, showing increased expression in an *A. thaliana* cell culture 30 min after exposure to high light stress conditions^162^. The putative orthologue of OG47629/FRAEX38873_v2_000173180 in *S. lycopersicum* is Solyc09g011350 and has been identified as a “disease-specific gene” that is differentially regulated in tomato leaves at 24 hours post inoculation with three pathogens^163^. Solyc09g011350 was upregulated in the infected leaves compared with the control, specifically in response to *Botrytis cinerea*^163^.

###### OG49074 (FRAEX38873_v2_000159740)

Grand-conv detected evidence of a convergent valine in *F. mandshurica* and *F. platypoda*, at a site where the other *Fraxinus* species in the analysis have a glycine (Supplementary Table 5). The top match for this gene in *A. thaliana* is *MEMBRANE-ASSOCIATED PROGESTERONE BINDING PROTEIN 4* (AT4G14965), or *MAPR4.* Although the function of AT4G14965/*MAPR4* has not been fully characterised, GO terms have been associated with it via identification of functional gene modules^118^, which are: NEGATIVE REGULATION OF CELLULAR PROCESS and POSTREPLICATION REPAIR (see http://bioinformatics.psb.ugent.be/cig_data/plant_modules/createResultsHTML.php?geneID=AT4G14965).

###### OG50989 (FRAEX38873_v2_000237270)

Grand-conv detected evidence of a convergent serine in *F. mandshurica* and *F. platypoda*, at a site where the other *Fraxinus* species in the analysis have a proline (Supplementary Table 5). The top match for this gene in *A. thaliana* is a gene encoding a calcium-binding EF-hand family protein (AT4G26470), which functions in calcium ion binding and calcium-mediated signaling (www.arabidopsis.org/servlets/TairObject?type=locus&name=At4g26470) and appears orthologous to FRAEX38873_v2_000237270 on the basis of phylogenetic analysis of their OrthoMCL cluster (Supplementary Fig. 3s). AT4G26470 is also known as *CML21* (short for *Calmodulin-like protein 21*)^164^ and is transcriptionally upregulated during pollen tube growth^165^. AT4G26470/*CML21* also shows increased expression in response to melatonin^166^; many of the other genes that also showed altered expression in response to melatonin levels were involved in stress defence response, indicating an important role for melatonin triggering defence response to both abiotic and biotic stresses^166^. Calcium signalling plays a key role in plant defence response^167^ including in pathogen-triggered programmed cell death^33^ and Ca^2+^ influx is one of the early signals of feeding insects and is involved in triggering defense response^17^.

###### OG59564 (FRAEX38873_v2_000174000)

Grand-conv detected evidence of a convergent aspartic acid in *F. mandshurica* and *F. platypoda*, at a site where the other *Fraxinus* species in the analysis have a glutamic acid (Supplementary Table 5). The top match for this gene in *A. thaliana* is *Na-translocating NADH-quinone reductase subunit A* (AT5G55640). AT5G55640 is also known as *MDF20.8* (www.uniprot.org/uniprot/Q9FM74). Information on the precise function of AT5G55640/*MDF20.8* is lacking, however, quinone reductases are considered to be detoxifying enzymes and have a role in protecting organisms from oxidative stress^168^. One of the susceptible taxa included in our convergence analyses, *F. latifolia*, may lack a functional copy of this gene (Supplementary Table 6).

###### OG60899 (FRAEX38873_v2_000255120)

Grand-conv detected evidence of a convergent serine in *F. mandshurica* and *F. platypoda*, at a site where the other *Fraxinus* species in the analysis have a glycine. However, the serine is also found in the alternate allele for susceptible *F. ornus* (Supplementary Table 5). The convergent site is predicted to be within a signal peptide, but the amino acid variant is not suggested to alter the localisation of the protein (Supplementary Table 5). The top match for this gene in *A. thaliana* is *EARLY NODULIN-LIKE PROTEIN 7* (AT1G79800), or *ENODL7*. Although the function of AT1G79800 has not been fully characterised, GO terms have been associated with it via identification of functional gene modules^118^, which are: COPPER ION BINDING and ELECTRON CARRIER ACTIVITY (http://bioinformatics.psb.ugent.be/cig_data/plant_modules/createResultsHTML.php?geneID=AT1G79800). Two *P. trichocarpa* genes belong to the same OrthoMCL cluster as FRAEX38873_v2_000255120 (Sollars et al.^11^) both of which (Potri.001G187700 and Potri.003G050500) have been categorised as AGP genes (highly glycosylated arabinogalactan-proteins), and are known as *PtPAG36* and *PtPAG37*^169^. However, there is no specific information on the function of these genes. One of the resistant taxa included in our convergence analyses, *F. floribunda*, may lack a functional copy of this gene (Supplementary Table 6). The FRAEX38873_v2_000255120 reference gene model from *F. excelsior* appears to be incorrect, possibly due to a misassembly (Supplementary Table 5).

###### OG21449 (FRAEX38873_v2_000266260)

Grand-conv detected evidence of a convergent threonine in *F. mandshurica*, *F. baroniana, F. floribunda* and *Fraxinus* sp. D2006-0159, at a site where the other *Fraxinus* species in the analysis have an isoleucine (Supplementary Table 5). The top match for this gene in *A. thaliana* is *PHOTOLYASE/BLUE-LIGHT RECEPTOR 2* (AT2G47590), also known as *PHR2*. Photolyases are light activated and repair damage caused to DNA by UV radiation; *PHR2* photolyases are specific to plants^170^.

###### OG23284 (FRAEX38873_v2_000124490)

Grand-conv detected evidence of a convergent phenylalanine and serine in *F. mandshurica*, *F. baroniana, F. floribunda* and *Fraxinus* sp. D2006-0159, at sites where the other *Fraxinus* species in the analysis have a cysteine and a leucine (Supplementary Table 5). The top match for this gene in *A. thaliana* is a gene encoding an alpha/beta-Hydrolases superfamily member (AT3G19970); phylogenetic analysis of the OrthoMCL cluster including FRAEX38873_v2_000124490 and AT3G19970 indicates they are likely to be orthologous (Supplementary Fig. 3t). Although the function of AT3G19970 has not been fully characterised, GO terms have been associated with it via identification of functional gene modules^118^, including: CALLOSE DEPOSITION DURING DEFENSE RESPONSE, SALICYLIC ACID MEDIATED SIGNALING PATHWAY, JASMONIC ACID MEDIATED SIGNALING PATHWAY, REGULATION OF PLANT-TYPE HYPERSENSITIVE RESPONSE, REGULATION OF IMMUNE RESPONSE, RESPONSE TO CHITIN (see http://bioinformatics.psb.ugent.be/cig_data/plant_modules/createResultsHTML.php?geneID=AT3G19970) indicating that this gene may potentially have a role related to plant defence response. The apparent orthologue of FRAEX38873_v2_000382300 in *S. lycopersicum* is Solyc02g020890 (Supplementary Fig. 3t); Solyc02g020890 was among 4774 genes in tomato whose expression was significantly affected by application of methyl jasmonate^171^.

###### OG47560 (FRAEX38873_v2_000262780)

Grand-conv detected evidence of a convergent lysine and phenylalanine in *F. mandshurica*, *F. baroniana, F. floribunda* and *Fraxinus* sp. D2006-0159, at sites where the other *Fraxinus* species in the analysis have a glutamic acid and tyrosine (Supplementary Table 5). The top match for this gene in *A. thaliana* is *TAPETUM 1* (AT3G42960), or *TA1*. AT3G42960/*TA1* is also known as *ASD* or *ATA1* and is expressed in tapetal cells; it has been reported to be involved in flower development and to have oxidoreductase activity (www.arabidopsis.org/servlets/TairObject?name=AT3G42960&type=locus). Phylogenetic analysis of the OrthoMCL cluster including FRAEX38873_v2_000262780 and AT3G42960/*TA1* suggests they may be paralogues, although this result is not conclusive (Supplementary Fig. 3u). The apparent orthologue of OG47560/FRAEX38873_v2_000262780 in *S. lycopersicum* is Solyc11g018600 (Supplementary Fig. 3u) and has decreased expression in a male-sterile tomato mutant^172^. The PhytoMine entry for Solyc11g018600 predicts that this gene is involved in the abscisic acid (ABA) biosynthesis pathway (https://phytozome.jgi.doe.gov/phytomine/report.do?id=286588936&trail=%7c286588936). AT3G42960/*TA1* is also predicted to be involved in the ABA biosynthesis pathway, although this has not yet been confirmed with experimental evidence (https://pmn.plantcyc.org/ARA/NEW-IMAGE?type=PATHWAY&object=PWY-695). A major function of ABA is response to abiotic stress^31^; ABA also plays an important role in triggering stomatal closure, which represents a plant defence mechanism^31^. Evidence suggests that ABA interacts with other phytohormones to influence response to herbivory and plant defence against pathogens^30,31^.

###### OG56563 (FRAEX38873_v2_000135960)

Grand-conv detected evidence of a convergent valine in *F. mandshurica*, *F. baroniana, F. floribunda* and *Fraxinus* sp. D2006-0159, at sites where the other *Fraxinus* species in the analysis have an alanine or threonine (Supplementary Table 5). However, the valine is also found in the alternate allele for susceptible *F. ornus* (Supplementary Table 8). The top match for this gene in *A. thaliana* is *CASP-LIKE PROTEIN 4D1* (AT2G39530), *CASPL4D1*. The function of AT2G39530/*CASPL4D1* is not well characterised, but it has been shown to have reduced expression in response to drought stress^173^. AT2G39530/*CASPL4D1* also showed upregulation in *A. thaliana* plants infiltrated with an avirulent strain of *Pseudomonas syringae* pv. tomato but downregulation after ABA induced susceptibility followed by pathogen inoculation^174^. However, FRAEX38873_v2_000135960 does not belong to the same OrthoMCL cluster as AT2G39530 (Sollars et al.^11^), and appears not to have an orthologue in *A. thaliana*. The putative orthologue of OG56563/FRAEX38873_v2_000135960 in *S. lycopersicum* is Solyc03g098090, but information on the function of this gene is apparently lacking.

###### OG64545 (FRAEX38873_v2_000393500)

Grand-conv detected evidence of a convergent serine in *F. mandshurica*, *F. baroniana, F. floribunda* and *Fraxinus* sp. D2006-0159 in a position where the other *Fraxinus* species in the test have an asparagine (Supplementary Table 5). The top match for this gene in *A. thaliana* is a gene encoding a single hybrid motif superfamily protein (AT2G35120) and is involved in glycine decarboxylation via the glycine cleavage system (www.arabidopsis.org/servlets/TairObject?name=AT2G35120&type=locus). AT2G35120 is also known as *GDCH2*^175^ or *GDH2* (short for *Glycine cleavage system H protein 2*; www.uniprot.org/uniprot/O82179). The glycine decarboxylase (GDC) multienzyme system (also known as the glycine cleavage system; www.uniprot.org/uniprot/O82179) consists of four enzymes (P, H, T and L proteins) and catalyses the destruction of glycine molecules emitted from the peroxisomes during photorespiration^176^. The H protein shuttles the methylamine group of glycine from the P protein to the T protein (www.uniprot.org/uniprot/O82179). GDC might be involved in plant stress-response, via cross-talk between nitric oxide and reactive oxygen species^177^. Two *P. trichocarpa* genes belong to the same OrthoMCL cluster as FRAEX38873_v2_000393500^11^ (Supplementary Fig. 3v) both of which (Potri.012G123700 and Potri.015G122500, also known as *gdcH1* and *gdcH2* respectively; https://phytozome.jgi.doe.gov/) have correlated expression with several genes involved in the phenylpropanoid pathway, including a *P. trichocarpa CYP98A3* putative orthologue (Potri.006G033300; https://phytozome.jgi.doe.gov/phytomine/report.do?id=48451623) which belongs to the same OrthoMCL cluster as another of our candidate loci, OG15551/FRAEX38873_v2_000261700 (Supplementary Fig. 3a). Potri.012G123700/*gdcH1* was found to be most highly expressed in xylem tissues and Potri.015G122500/*gdcH2* also shows evidence of higher expression in wood tissues than in numerous other tissue types examined^178^ (see http://popgenie.org/gene?id=Potri.T122200). Both poplar genes contain the AC element within their promoters^179^. This element also occurs within many phenylpropanoid biosynthesis genes in *P. trichocarpa* (including *PAL*, *C4H*, *C3’H*, *CCR* and *CAD*), suggesting that the expression of Potri.012G123700/*gdcH1* and Potri.015G122500/*gdcH2* is coordinated with that of phenylpropanoid genes to support lignin and flavonoid biosynthesis^179^. This role for GDC H protein genes may be a tree-specific adaptation in response to high demands for one-carbon units during lignification, which is distinct from the role of *GDC* genes in *A. thaliana*^179^.

###### OG16739 (FRAEX38873_v2_000338500)

Grand-conv detected evidence of a convergent alanine in *F. platypoda*, *F. baroniana, F. floribunda* and *Fraxinus* sp. D2006-0159, at a site where the other *Fraxinus* species in the analysis have a threonine (Supplementary Table 5). However, the alanine is also found in the alternate allele for susceptible *F. ornus* (Supplementary Table 8). The top match for this gene in *A. thaliana* is a gene encoding a RNA-binding (RRM/RBD/RNP motifs) protein family member (AT2G41060), which is also known as *UBA2B*. Phylogenetic analysis of the OrthoMCL cluster including FRAEX38873_v2_000338500 and AT2G41060 indicates they are paralogues rather than orthologues (Supplementary Fig. 3w) and that FRAEX38873_v2_000338500 is equally closely related to AT3G56860, also known as *UBA2A*. Overexpression of both *UBA2A* and *UBA2B* leads to increased expression of a number of senescence-associated and defence related genes, as well as to increased ethylene biosynthesis, hypersentive-like patterns of cell death and callose deposition^41^. Expression of the *UBA2* genes was not found to increase in relation to natural, age-related, senescence, but instead appears connected to wound-induced cell-death and senescence pathways, supporting the suggestion that these genes play a role in regulating senescence in response to wounding and other environmental cues^41^.

###### OG20520 (FRAEX38873_v2_000382300)

Grand-conv detected evidence of a convergent isoleucine in *F. platypoda*, *F. baroniana, F. floribunda* and *Fraxinus* sp. D2006-0159, at a site where the other *Fraxinus* species in the analysis have a serine (Supplementary Table 5). The top match for this gene in *A. thaliana* is *SHOOT GRAVITROPISM 7* (AT4G37650), which is also known as *SHORT ROOT (SHR)*. AT4G37650/*SHR* is essential for root ground tissue patterning and interacts with *SCARECROW/SCR*^180^; they are thought to function as transcriptional regulators^180^. The putative orthologue of OG20520/FRAEX38873_v2_000382300 in *S. lycopersicum* is Solyc02g092370; this gene is also known as *SlSHRa* and is involved in root radial patterning^181^. The putative orthologue from *Populus trichocarpa*, Potri.007G063300 (being the only *P. trichocarpa* gene belonging to the same OrthoMCL cluster as FRAEX38873_v2_000382300^11^), is also known as *PtSHR1*^182^ and down-regulation of this gene in transgenic hybrid poplar leads to increased height and girth, indicating that *SHR* is involved in regulation of cell division and meristem activity in shoots as well as roots^183^.

###### OG21033 (FRAEX38873_v2_000092000)

Grand-conv detected evidence of a convergent lysine in *F. platypoda*, *F. baroniana, F. floribunda* and *Fraxinus* sp. D2006-0159, at a site where the other *Fraxinus* species in the analysis have a glutamine (Supplementary Table 5). The top match for this gene in *A. thaliana* is an O-Glycosyl hydrolases family 17 protein encoding gene (AT3G55430); proteins in this family include those with a number of β-glucosidase related activities (www.cazy.org/GH17.html) and are involved in hydrolysis of glycosidic bonds in carbohydrates (http://www.ebi.ac.uk/interpro/entry/IPR000490). Phylogenetic analysis of the OrthoMCL cluster including FRAEX38873_v2_000092000 indicates it is equally closely related to AT3G55430 and another O-Glycosyl hydrolases family 17 protein encoding gene from *A. thaliana*, AT2G39640, and appears orthologous to both (Supplementary Fig. 3x). Expression of AT3G55430 seems to be at least in part dependent on nitric oxide, and *A. thaliana* mutants where AT3G55430 function is impaired have increased susceptibility to the fungal pathogen *Botrytis cinerea*^184^. It is suggested that the control of AT3G55430 expression via nitric oxide might form part of the mechanism for basal resistance to *B. cinerea* in *A. thaliana*^184^. In the predicted protein model for OG21033/FRAEX38873_v2_000092000 the lysine/glutamine variant is close to the active site (Supplementary Fig. 2d).

###### OG27693 (FRAEX38873_v2_000056890)

Grand-conv detected evidence of a convergent isoleucine in *F. platypoda*, *F. baroniana, F. floribunda* and *Fraxinus* sp. D2006-0159, at a site where the other *Fraxinus* species in the analysis have a threonine or alanine (Supplementary Table 5). The top match for this gene in *A. thaliana* is *HESPERIN* (AT1G31500); this gene is a transcriptional regulator of circadian rhythms but apparently also has a role in response to oxidative stress^185^.

###### OG35707 (FRAEX38873_v2_000303580)

Grand-conv detected evidence of a convergent leucine in *F. platypoda*, *F. baroniana, F. floribunda* and *Fraxinus* sp. D2006-0159, at a site where the other *Fraxinus* species in the analysis have a phenylalanine (Supplementary Table 5). The top match for this gene in *A. thaliana* is *ZRT/IRT-LIKE PROTEIN 2* (AT5G59520), or *ZIP2*. AT5G59520/*ZIP2* probably plays a role in Mn (and possibly Zn) transport into the root vasculature for translocation to the shoot^186^. In plants, Mn is a cofactor in processes such as photosynthesis, lipid biosynthesis and oxidative stress^187^. Three *P. trichocarpa* genes belong to the same OrthoMCL cluster as FRAEX38873_v2_000303580 (Sollars et al.^11^) one of which, Potri.009G034600, is also known as *ZIP2* and is involved in Cd^2+^ uptake^188^.

###### OG36502 (FRAEX38873_v2_000266620)

Grand-conv detected evidence of a convergent serine in *F. platypoda*, *F. baroniana, F. floribunda* and *Fraxinus* sp. D2006-0159, at a site where the other *Fraxinus* species in the analysis have an asparagine (Supplementary Table 5). The top match for this gene in *A. thaliana* is *TRNA METHYLTRANSFERASE 140B* (AT1G54650), which encodes a methyltransferase family protein. AT1G54650/*TRM140B* is proposed to have a role in tRNA nucleoside methylation, generating m3C (3-methylcytidine) modifications^189^. The serine variant in *Fraxinus* is a putative phosphorylation site (Supplementary Table 5), and is found on the exterior of the protein in the predicted model for OG36502/FRAEX38873_v2_000266620 (Supplementary Fig. 2a).

###### OG43828 (FRAEX38873_v2_000227880)

Grand-conv detected evidence of a convergent leucine in *F. platypoda*, *F. baroniana, F. floribunda* and *Fraxinus* sp. D2006-0159, at a site where the other *Fraxinus* species in the analysis have a serine (Supplementary Table 5). The top match for this gene in *A. thaliana* is *ATDET2* (AT2G38050), a gene involved in the brassinolide (brassinosteroid) biosynthetic pathway. Phylogenetic analysis indicates that FRAEX38873_v2_000227880 and AT2G38050/*ATDET2* are likely to be orthologues (Supplementary Fig. 3y).

AT2G38050/*ATDET2* is also known as *DWARF 6* (*DWF6*) and is involved in multiple reactions within the brassinosteriod biosynthesis pathway^190^. Brassinosteroids are hormones whose functions in plants include regulating growth and development^191^ and which also play an important part in modulating growth–defence trade-offs^135^. Brassinosteriods in plants have potential role in mediating response to stresses such as freezing, drought, salinity, disease, heat and nutrient deficiency^191^. There is also evidence for a role for brassinosteriods in regulating glucosinolate profiles, which function in defence response against insects in *A. thaliana*^192^. The *det2-1* mutant in *A. thaliana* has impaired stomatal opening in response to blue light, suggesting a further possible role for brassinosteroids in stomatal opening^193^. One of the apparent orthologues of OG43828/FRAEX38873_v2_000227880 in *P. trichocarpa*, Potri.T122200 (Supplementary Fig. 3y), was found to be preferentially expressed in beetle damaged leaves^178^ (see http://popgenie.org/gene?id=Potri.T122200). The FRAEX38873_v2_000227880 reference gene model from *F. excelsior* appears to be incomplete on the 5′ end, apparently due to a frameshift induced by the assembly of a chimeric haplotype combining variants from both alleles (Supplementary Table 5).

#### Supplementary Note 5

##### 5. Significant enrichment of GO terms

Three GO terms are significantly enriched among the refined list of 53 candidates (*p*-value < 0.01) relative to the set of genes included in the grand-conv analyses, according to Fisher’s exact test run with the weight algorithm, all within the biological process (BP) domain and all related to hormone metabolic processes: estrogen metabolic process, C21-steroid hormone metabolic process and androgen metabolic process (Supplementary Table 7). These terms are associated with two of the 53 candidates, OG40061 (FRAEX38873_v2_000309460) and OG43828 (FRAEX38873_v2_000227880), the latter of which appears to be involved in brassinosteroid biosynthesis (see Supplementary Note 4.5). The same terms were found to be significantly enriched when running Fisher’s exact test run with the elim algorithm. However, two additional terms within the BP domain were also significantly enriched: hormone biosynthetic process and organic cyclic compound biosynthetic process (Supplementary Table 7). Three of the candidate genes are associated with the term “hormone biosynthetic process”, OG16739 (FRAEX38873_v2_000338500.1), OG39275 (FRAEX38873_v2_000319270.1) and OG43828 (FRAEX38873_v2_000227880). Nine candidate genes are associated with the term “organic cyclic compound biosynthetic process” including some that appear to be involved in phytohormone biosynthesis (Supplementary Table 7 and Supplementary Note 4). Also, two terms within the molecular function (MP) domain were found to be significantly enriched when using the elim algorithm: steroid dehydrogenase activity and carboxy-lyase activity (Supplementary Table 7). Both of these terms are associated with OG40061 (FRAEX38873_v2_000309460) and OG43828 (FRAEX38873_v2_000227880). Some of the genes among the top 53 candidates did not have any GO terms annotated by the analysis performed by (Sollars et al.^11^), which may explain why some of the candidates that appear to be involved in phytohormone biosynthesis (Supplementary Note 4) are not associated with the significantly enriched GO terms.

#### Supplementary Note 6

##### 6. Differential expression in response to EAB-feeding

For 29 the 53 loci in our refined set of candidates we were able to identify the likely orthologue from the published *F. pennsylvanica* transcriptome^25^; of these 29 *F. pennsylvanica* genes, 11 showed evidence for differential expression subsequent to EAB-feeding (four with decreased expression, and seven with increased expression; Supplementary Table 10)^25^. However, none of these genes showed a difference in expression pattern between susceptible and resistant *F. pennsylvanica* individuals, suggesting that the majority of these loci may not play a critical role in governing the degree to which this species can defend itself against EAB. Nevertheless, loci that show a response to EAB, but where expression patterns are not differentiated between individuals with contrasting levels of susceptibility, may form part of a suite of genes that are required for defence, with trees that succumb fully to the insect lacking further essential components. Alternatively, these differentially expressed genes may reflect other changes that impacted the seedlings during the eight week period between collection of the “pre” and “post” EAB-feeding samples^25^.

